# “Self-inactivating” rabies viruses are susceptible to loss of their intended attenuating modification

**DOI:** 10.1101/550640

**Authors:** Lei Jin, Makoto Matsuyama, Heather A. Sullivan, Mulangma Zhu, Thomas K. Lavin, YuanYuan Hou, Nicholas E. Lea, Maxwell T. Pruner, María Lucía Dam Ferdínez, Ian R. Wickersham

**Author notes:** **Competing interests statement** I.R.W. is a consultant for Monosynaptix, LLC, advising on design of neuroscientific experiments.

## Abstract

Monosynaptic tracing using rabies virus is an important technique in neuroscience, allowing brain-wide labeling of neurons directly presynaptic to a targeted neuronal population. A 2017 article reported development of a noncytotoxic version – a major advance – based on attenuating the rabies virus by addition of a destabilization domain to the C-terminus of a viral protein. However, this modification did not appear to hinder the ability of the virus to spread between neurons. We analyzed two viruses provided by the authors and show here that both were mutants that had lost the intended modification, explaining the paper’s paradoxical results. We then made a virus that actually did have the intended modification in at least the majority of virions and found that it did not spread efficiently under the conditions described in the original paper, namely, without an exogenous protease being expressed in order to remove the destabilization domain. We found that it did spread when the protease was supplied, although this also appeared to result in the deaths of most source cells by three weeks postinjection. We conclude that the new approach is not robust but that it could become a viable technique given further optimization and validation.

**SIGNIFICANCE STATEMENT:** Rabies virus, which spreads between synaptically-connected neurons, has been one of the primary tools used by neuroscientists to reveal the organization of the brain. A new modification to rabies virus was recently reported to allow the mapping of connected neurons without adverse effects on the cells’ health, unlike earlier versions. Here we show that the conclusions of that study were probably incorrect and based on having used viruses that had lost the intended modification because of mutations. We also show that a rabies virus that does retain the intended modification does not spread efficiently between neurons under the conditions reported previously; however, it does spread between neurons under different conditions, suggesting that the approach may be successful if refined further.

## INTRODUCTION

Viruses have become important tools for neuroscience (1–16), and “monosynaptic tracing” based on rabies virus (5) has become the primary method of labeling neurons directly presynaptic to some targeted group of neurons (17–20). Its core components are, first, selective infection of the targeted neuronal group with a recombinant rabies virus with a deleted gene (which in all work published to date is the “G” gene encoding the glycoprotein that coats the viral envelope) and, second, complementation of the deletion in the targeted starting neurons, by expression of the deleted gene in *trans*. With all of its gene products therefore present in the starting cells, the virus can fully replicate within them and spreads, as wild-type rabies virus does, to cells directly presynaptic to the initially-infected neurons. Assuming that G has not been provided in *trans* in these presynaptic cells too, the deletion-mutant (“ΔG”, denoting the deletion of G) virus is unable to spread beyond them, resulting in labeling of just the neurons in the initially-targeted population and ones that are directly presynaptic to them (5).

A drawback of these ΔG (or “first-generation” (21)) rabies viruses is that they are cytotoxic (4, 21, 22), which has spurred several labs to develop less toxic versions. Reardon, Murray, and colleagues (22) showed that simply using ΔG rabies virus of a different parent strain — switching from the original SAD B19 strain to the more neuroinvasive CVS-N2c strain (23) — decreased toxicity and increased the efficiency of transneuronal spread. Our own group has taken a more drastic approach, introducing nontoxic “second-generation” rabies viruses from which both G and a second gene, “L”, encoding the viral polymerase, have been deleted (21); when complemented by expression of both deleted genes *in trans*, these double-deletion-mutant vectors spread to presynaptic neurons, just as with the first-generation system (24).

Taking a quite different approach, Ciabatti et al. (25) introduced “self-inactivating rabies” (“SiR”) viruses, which differed from simple first-generation (ΔG, SAD B19 strain) ones by the addition of a destabilization domain to the C-terminus of one of the viral proteins, so that the protein would be rapidly degraded soon after it was produced. Because the protein in question (the nucleoprotein, encoded by the “N” gene) is essential for viral gene expression and replication, its destabilization was intended to “silence” viral gene expression and prevent replication, making the viruses nontoxic.

These SiR viruses were designed to be unable to replicate unless an exogenous protease (tobacco etch virus protease, TEVP) was expressed in infected cells in order to remove the destabilization (or “PEST” (26)) domain. However, the paper reported that they were able to spread between neurons – which requires replication – just as efficiently as unmodified first-generation viruses did, without the protease being provided at all.

We hypothesized that the viruses that were used for the reported transsynaptic tracing experiments (25) were mutants with premature stop codons at or near the end of the native nucleoprotein gene and before the sequence of the destabilization domain. Rhabdoviruses have high mutation rates (27–32), and production of high-titer rabies virus stocks for *in vivo* injection typically involves repeated passaging on complementing cell lines (33–35), which affords ample opportunity for accumulation of mutants with a selective replication advantage.

Here we show that, in both of the two SiR virus samples that we analyzed, the great majority of viral particles did have mutations in their genomes that caused the complete loss of the intended C-terminal addition to the nucleoprotein, so that they were effectively just ordinary first-generation ΔG rabies viral vectors. We also tested the SiR-CRE virus *in vivo* and found that it was rapidly cytotoxic.

We also show that a ΔG virus that does have the intended modification to the nucleoprotein does not spread efficiently *in vivo* in the absence of TEVP, but that it does spread efficiently when TEVP is provided.

## RESULTS

We analyzed samples of two viruses sent directly from the Tripodi lab to MIT two months after their publication(25) and given directly to the Wickersham lab soon afterward, still frozen and unopened, with express permission from Marco Tripodi by email on December 5, 2017: “EnvA/SiR-CRE” (made from genome plasmid Addgene 99608, pSAD-F3-NPEST-iCRE-2A- mCherryPEST) and “EnvA/SiR-FLPo” (made from genome plasmid Addgene 99609, pSAD- F3-NPEST-FLPo-2A-mCherryPEST). Both had the SAD B19 strain of rabies virus as their parent strain and had been packaged with the avian and sarcoma virus subgroup A envelope glycoprotein (“EnvA”) for targeted infection of cells expressing EnvA’s receptor, TVA (5).

For comparison with the two SiR viruses, we made five control viruses in our own laboratory: three first-generation vectors, RVΔG-4Cre (21), RVΔG-4FLPo (see Methods), and RVΔG-4mCherry (36), and two second-generation vectors, RVΔGL-4Cre and RVΔGL-4FLPo (21). All of these viruses are on the SAD B19 background, like the SiR viruses. For each of the four recombinase-expressing viruses from our laboratory, we made one preparation packaged with the EnvA envelope protein and one preparation packaged with the native rabies virus (SAD B19 strain) glycoprotein (denoted as “B19G”); RVΔG-4mCherry (used only as a control for the Sanger sequencing) was only packaged with the EnvA envelope protein.

### Sequencing of viral genomes: Sanger sequencing

In order to directly test our hypothesis that the SiR viruses had developed premature stop codons removing the PEST domain in a majority of viral particles, we sequenced the genomes of a large number of individual viral particles using two different techniques.

First, we used ordinary Sanger sequencing to determine the sequence in the vicinity of the end of the nucleoprotein gene for 50 to 51 individual viral particles of each of the two SiR viruses and of a first-generation virus from our own laboratory, RVΔG-4mCherry (Figure 1 and Supplementary File S1). We ensured the isolation of individual viral genomes by using a primer with a random 8-base index for the reverse transcription step, so that the cDNA copy of each RNA viral genome would have a unique index. Following the reverse transcription step, we amplified the genomes by standard PCR, cloned the amplicons into a generic plasmid, transformed this library into E.coli and sequenced plasmids purified from individual colonies.

**Figure 1:**
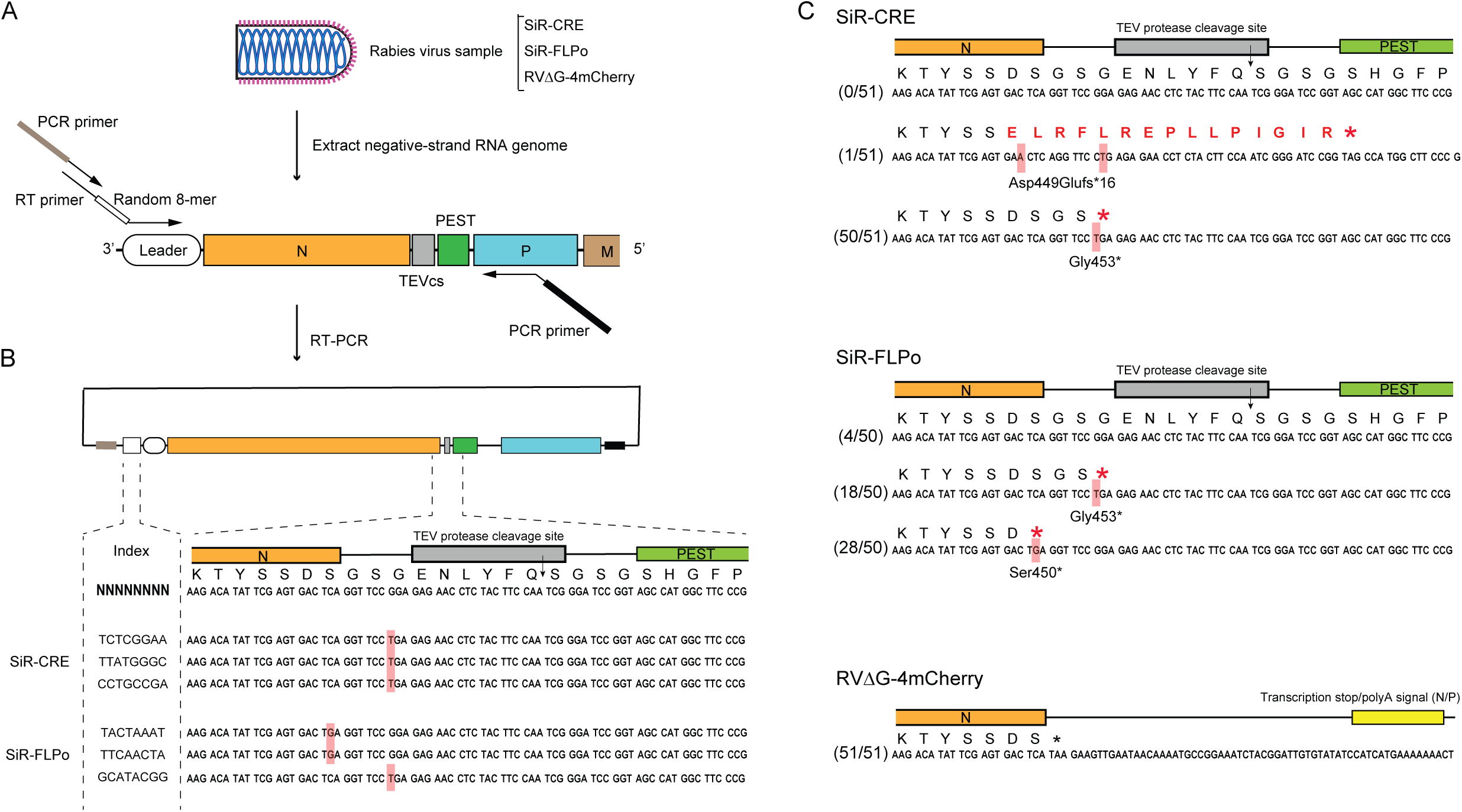
Single-molecule Sanger sequencing of individual viral genomes revealed that most SiR virions had lost the intended C-terminal modification to the nucleoprotein. (A) Schematic of the RT-PCR workflow. In the reverse transcription (RT) step, the RT primer, containing a random 8-nucleotide sequence, anneals to the 3’ rabies virus leader, adding a unique random index to the 5’ end of the cDNA molecule corresponding to each individual viral particle’s RNA genome. In the PCR step, the forward PCR primer anneals to the RT primer sequence and the reverse PCR primer anneals within the viral phosphoprotein gene P. Both PCR primers have 15-base sequences homologous to those flanking an insertion site in a plasmid used for sequencing, allowing the amplicons to be cloned into the plasmid using a seamless cloning method before transformation into bacteria. The resulting plasmid library consists of plasmids containing up to 4^8^ different index sequences, allowing confirmation that the sequences of plasmids purified from individual picked colonies correspond to the sequences of different individual rabies viral particles’ genomes. (B) Representative Sanger sequencing data of the 8-bp index and the TEV-PEST sequence. Mutations are highlighted in red. (C) Mutation variants and their frequencies in each viral vector sample based on Sanger sequencing data. No unmutated genomes were found in the SiR-CRE sample: 50 out of 51 had a substitution creating an opal stop codon just before the TEV cleavage site, and the 51^st^ genome contained a frameshift which also removed the C-terminal addition. In the SiR-FLPo sample, only 4 out of 50 clones had an intact sequence of the C-terminal addition; the other 46 out of 50 had one of two *de novo* stop codons at the end of N or the beginning of the TEV cleavage site. In the sample of RVΔG-4mCherry, a virus from our laboratory included as a control to distinguish true mutations on the rabies genomes from mutations due to the RT-PCR process, none of the 51 clones analyzed had mutations in the sequenced region.

As shown in Figure 1, the results confirmed our hypothesis that SiR viruses are prone to loss of the 3’ addition to the nucleoprotein gene. Specifically, in the SiR-CRE sample, 100% of the 51 sequenced viral particles had lost the PEST domain. Fifty out of the 51 had the same point mutation in the linker between the end of the native nucleoprotein gene and the TEVP cleavage site, converting a glycine codon (GGA) to a stop codon (TGA) so that the only modification to the C-terminus of the nucleoprotein was the addition of two amino acids (a glycine and a serine). The one sequenced viral particle that did not have this point mutation had a single-base insertion in the second-to-last codon of the native nucleoprotein gene, frameshifting the rest of the sequence and resulting in 15 amino acids of nonsense followed by a stop codon before the start of the PEST domain sequence.

In the SiR-FLPo sample, the population was more heterogeneous: out of 50 sequenced viral particles, 18 had the same stop codon that was found in almost all genomes in the Cre sample, while another 28 had a different stop codon three amino acids upstream, immediately at the end of the native nucleoprotein gene (converting a serine codon (TCA) to a stop codon (TGA)). Four viral particles had no mutations in the sequenced region. Thus 46/50 (92%) of the SiR-FLPo viral particles sequenced had lost the PEST domain.

In contrast, in the first-generation virus from our own lab, RVΔG-4mCherry, none of the 50 viral particles sequenced had mutations in the sequenced region containing the end of the nucleoprotein gene.

### Sequencing of viral genomes: Single-molecule, real-time (SMRT) sequencing

As a second approach to analyzing the mutations present in the SiR viruses, we employed a large-scale sequencing technology: single-molecule, real-time (“SMRT”) sequencing, which provides independent sequences of tens of thousands of individual single molecules in a sample in parallel (Figure 2 and Supplementary File S2). The results from this advanced sequencing method were quite consistent with the results from the Sanger sequencing presented above. As with the sample preparation for Sanger sequencing, we included a random index (10 bases, in this case) in the reverse transcription primer, so that again the cDNA copy of each RNA viral genome molecule would be labeled with a unique index.

**Figure 2:**
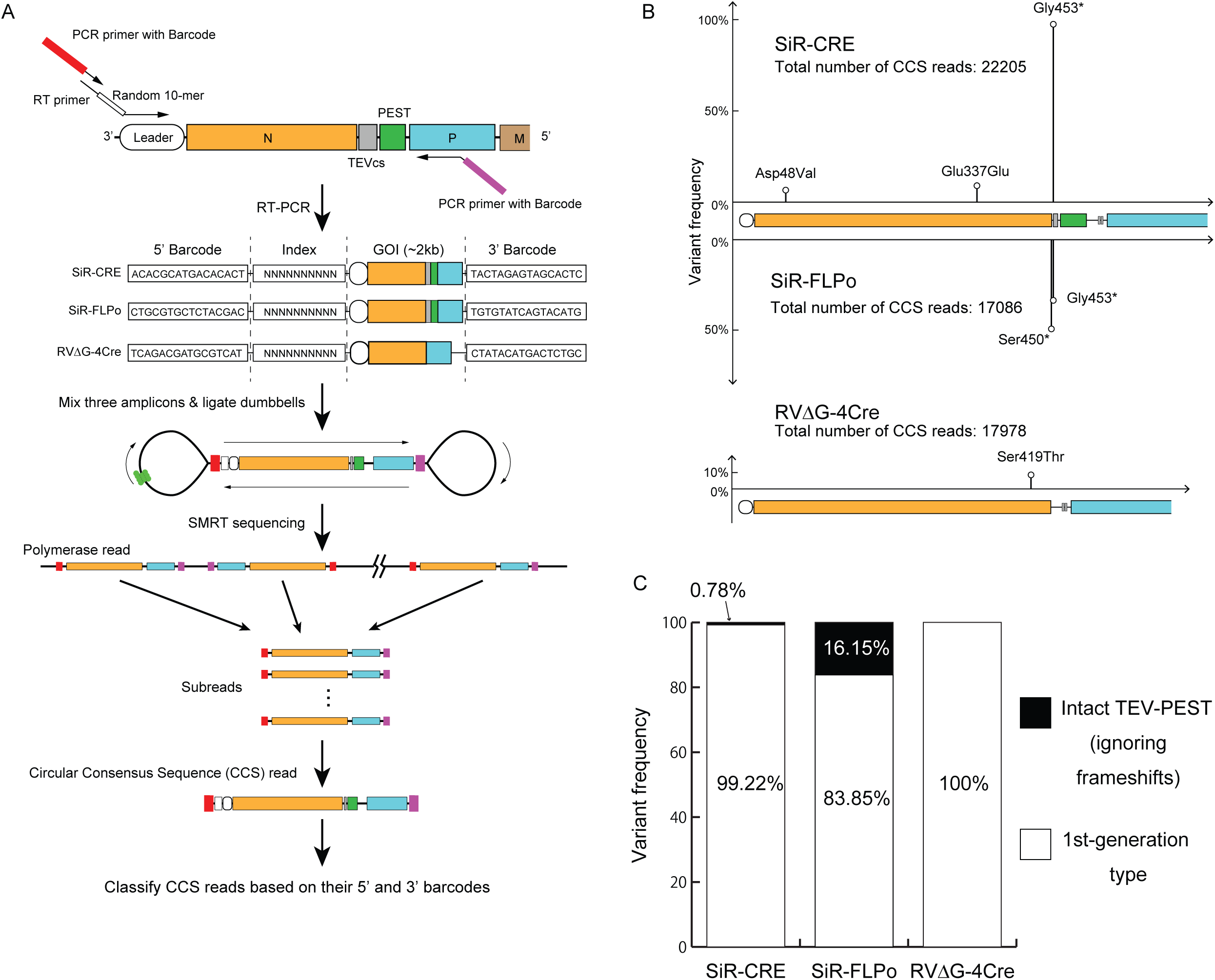
Single-molecule, real-time (SMRT) sequencing of thousands of barcoded viral genomes confirmed that most SiR virions had lost the intended C-terminal modification to the nucleoprotein. (A) Schematic of workflow for SMRT sequencing. An RT primer with a random 10-nucleotide sequence anneals to the leader sequence on the negative-sense single-stranded RNA genome. Forward and reverse PCR primers have distinct SMRT barcodes at their 5’ ends for the three different virus samples. After RT-PCR, each amplicon library consists of amplicons each containing a SMRT barcode to identify the sample of origin as well as the 10-nucleotide index (i.e., with a potential diversity of 4^10^ different indices) to uniquely label the individual genome of origin. SMRT “dumbbell” adaptors are then ligated to the amplicons’ ends, making circular templates which are then repeatedly traversed by a DNA polymerase, resulting in long polymerase reads each containing multiple reads of the positive and negative strands. The individual subreads for a given molecule are combined to form circular consensus sequence (CCS) reads. (B) High-frequency (>2%) point mutations found in the rabies vector samples based on SMRT sequencing. Horizontal axis represents nucleotide position along the reference sequences (see text); vertical axis represents variant frequency. Total number of CCS3 reads (i.e., with at least 3 subreads for each position) are 22,205 for SiR-CRE, 17,086 reads for SiR-FLPo, and 17,978 reads for RVΔG-4Cre. The great majority of SiR-CRE and SiR-FLPo genomes have point mutations creating premature stop codons at or just after the end of N and before the C-terminal addition. The only frequent (>2%) mutation found in the control virus, RVΔG-4Cre, was a single amino acid substitution at position 419 in 9.49% of virions. Insertions and deletions are not shown here (see text). (C) Summary of results. In the SiR virus samples, 99.22% of SiR-CRE virions and 83.85% of SiR- FLPo virions had point mutations creating premature stop codons that completely removed the intended C-terminal addition to the nucleoprotein, making them simply first-generation (ΔG) rabies viral vectors. This does not include any insertions or deletions causing frameshifts (see text), which would further increase the percentage of first-generation-type virions in these samples. In the RVΔG-4Cre sample, there were no premature stop codons at or near the end of the nucleoprotein gene.

SMRT sequencing entails circularization of the DNA amplicons and multiple consecutive passes around the resulting circular molecule, with the redundancy provided by this repeated sequencing of each position increasing the signal to noise ratio and statistical significance of the results. The numbers presented in Figure 2 and below use the default of including only clones that had at least three reads of each base (“circular consensus sequence 3”, or “CCS3” in Supplementary File S2). Using the increasingly stringent criteria of requiring either five or eight reads per base (CCS5 or CCS8) reduced the numbers of qualifying genomes in all cases and changed the percentages slightly but gave very similar results overall. Because read accuracy for SMRT sequencing is ≥98% for circular consensus sequencing with 3 passes (see https://www.mscience.com.au/upload/pages/pacbio/technical-note---experimental-design-for-targeted-sequencing.pdf), we used a conservative threshold of 2% frequency of any given point mutation position in order to screen out false positives. Also to be very conservative, for Figure 2 we ignored all apparent frame shifts caused by insertions and deletions, because insertions in particular are prone to false positives with SMRT sequencing (37). See Supplementary File S2 for details, including details of frameshifts due to insertions; Supplementary Files S3-S5 contain the sequences of the PCR amplicons that would be expected based on published sequences of the three viruses, but to summarize here:

As a control, we used a virus from our own laboratory, RVΔG-4Cre (21) (see Addgene #98034 for reference sequence). Out of 17,978 sequenced genomes of this virus, we found no mutations above threshold frequency at the end of N. We did find that 1,706 viral particles (9.49%) had a nonsynonymous mutation (TCT (Ser) è ACT (Thr)) farther up in N at amino acid position 419 (31 amino acids upstream of the end of the 450-aa native protein). We do not know if this mutation is functionally significant, although it is not present in CVS-N2c (38), HEP-Flury (39), ERA (40), or Pasteur strains (Genbank GU992320), so these particles may effectively be N- knockouts that were propagated by coinfection with virions with intact N (see Discussion for more on such parasitic co-propagating mutants).

For the SiR-CRE virus, out of 22,205 viral genomes sequenced, 22,032 had the premature stop codon (GGA -> TGA) in the linker between the native nucleoprotein gene and the TEVP cleavage site sequence. In other words, even without including frameshifts, at least 99.22% of the individual viral particles in the SiR-CRE sample were essentially first-generation ΔG vectors, with the only modification of the nucleoprotein being an additional two amino acids at the C-terminus.

For the SiR-FLPo virus, out of 17,086 viral genomes sequenced, 5,979 had the stop codon (GGA -> TGA) in the linker, 8,624 had the stop codon (TCA -> TGA) at the end of N, and a further 28 had a different stop codon (TCA -> TAA) at the same position at the end of N. Of these, 305 viral particles had premature stop codons at both of these two positions, so that the total number of viral particles with one or both stop codons immediately before the PEST domain was (8624 + 5979 + 28 - 305 = 14,326. In other words, again even without including frameshifts, at least 83.85% of the individual viral particles in the SiR-FLPo sample were essentially first-generation ΔG vectors, with the only modification of the nucleoprotein being either two amino acids added to, or one amino acid lost from, the C-terminus.

### Anti-nucleoprotein immunostaining

We infected reporter cell lines with serial dilutions of the two EnvA-enveloped SiR viruses as well as the eight recombinase-expressing ones from our own lab: ΔG vs. ΔGL, Cre vs. FLPo, EnvA vs. B19G envelopes. Three days later, we immunostained the cells for rabies virus nucleoprotein and imaged the cells with confocal microscopy.

As seen in Supplementary Figure S1, we found that the cells infected with the SiR viruses looked very similar to those infected with the first-generation, ΔG viruses. Notably, the viral nucleoprotein, which in the SiR viruses is intended to be destabilized and degrade rapidly in the absence of TEVP, accumulated in the SiR-infected cells in clumpy distributions that looked very similar to those in the cells infected with the first-generation, ΔG viruses. By contrast, the cells infected with the second-generation, ΔGL viruses, which we have shown to be noncytotoxic (21), did not show any such nucleoprotein accumulation, clumped or otherwise, only punctate labeling presumably indicating isolated viral particles or post-infection uncoated viral particles (ribonucleoprotein complexes) that are not replicating.

### Longitudinal two-photon imaging *in vivo*

To see whether the SiR viruses kill neurons in the brain, we conducted longitudinal two-photon imaging *in vivo* of virus-labeled neurons in visual cortex of tdTomato reporter mice, as we had done previously to demonstrate the nontoxicity of second-generation rabies virus (21) (Figure 3). Because the SiR viruses were EnvA-enveloped, we first injected a lentivirus expressing EnvA’s receptor TVA, then one week later we injected either SiR-CRE or one of two EnvA-enveloped viruses made in our laboratory: the first-generation virus RVΔG-4Cre(EnvA) or the second-generation virus RVΔGL-4Cre(EnvA). Beginning one week after rabies virus injection, we imaged labeled neurons at the injection site every seven days for four weeks, so that we could track the fate of individual neurons over time.

**Figure 3:**
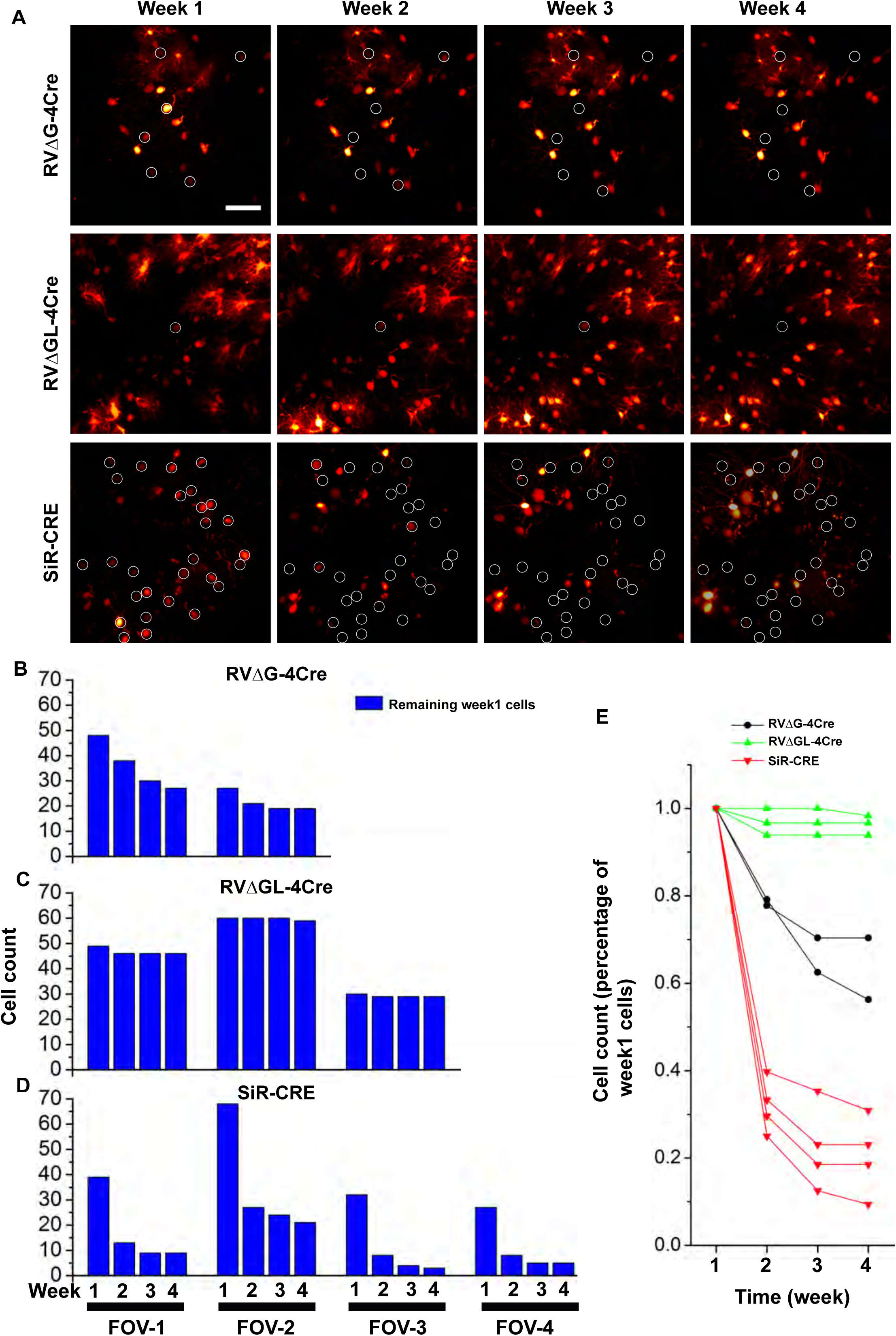
Longitudinal two-photon imaging *in vivo* showed that SiR virus killed approximately 80% of infected neurons in vivo within 2-4 weeks. A) Representative fields of view (FOVs) of visual cortical neurons labeled with RVΔG-4Cre (top row), RVΔGL-4Cre (middle row), or SiR-CRE (bottom row) in Ai14 mice (Cre-dependent expression of tdTomato). Images within each row are of the same FOV imaged at the four different time points in the same mouse. Circles indicate cells that are present at 7 days postinjection but no longer visible at a subsequent time point. Scale bar: 50 µm, applies to all images. B-D) Numbers of cells present at week 1 that were still present in subsequent weeks. While very few cells labeled with RVΔGL-4Cre were lost, and RVΔG-4Cre killed a significant minority of cells, SiR-CRE killed the majority of labeled neurons within 14 days following injection. Each FOV is from a single mouse: n = 2 mice in B, 3 mice in C, and 4 mice in D. E) Percentages of cells present at week 1 that were still present in subsequent imaging sessions. By 28 days postinjection, an average of only 20.5% of cells labeled by SiR-CRE remained.

As we found in our previous work (21), our second-generation virus RVΔGL-4Cre did not kill neurons to any appreciable degree: all but a tiny handful of the neurons labeled by this virus at seven days after injection were still present three weeks later in all mice. Also as we have found previously (21), our first-generation virus RVΔG-4Cre did kill neurons, but by no means all of them (see the Discussion for possible reasons for this).

However, we found that the putatively nontoxic SiR-CRE caused a steep loss of neurons much more pronounced than even our first-generation virus did. By 14 days after injection, 70% of cells seen at seven days were dead; by 28 days, 81% were.

There is a possible confound from our use of the tdTomato reporter line Ai14 (which we used primarily because we already had large numbers of mice of this line): because SiR-CRE is actually “SiR-iCRE-2A-mCherryPEST”, designed to coexpress mCherry (with an added C-terminal PEST domain intended to destabilize it, as for the nucleoprotein) along with Cre, it is conceivable that some of the SiR-CRE-labeled red cells at seven days were only expressing mCherry and not tdTomato. If the destabilized mCherry were expressed only transiently, as intended(25), and a significant fraction of SiR-CRE virions had mutations in the Cre gene so that they did not express functioning Cre, then it is possible that some of the red cells seen at seven days were labeled only with mCherry that stopped being visible by 14 days, so that it would only look like those cells had died.

We viewed this alternative explanation as unlikely, because the designers of SiR-CRE had injected it in an EYFP reporter line and found no cells labeled only with mCherry and not EYFP at six days and nine days postinjection (see Figure S4 in Ciabatti et al. (25)). Nevertheless, we addressed this potential objection in several ways.

First, we sequenced the transgene inserts (iCre-P2A-mCherryPEST) of 21 individual SiR- CRE viral particles (see Supplementary File S6) and found that only two out of 21 had mutations in the Cre gene, suggesting that there would not have been a large population of cells only labeled by mCherry and not by tdTomato.

Second, we repeated some of the SiR-CRE injections and imaging in a different reporter line: Ai35, expressing Arch-EGFP-ER2 after Cre recombination (41) (Jax 012735). Although we found that the membrane-localized green fluorescence from the Arch-EGFP-ER2 fusion protein was too dim and diffuse at seven days postinjection to be imaged clearly, we were able to obtain clear images of a number of cells at 11 days postinjection. We found that 46% of them had disappeared only three days later (see Supplementary Figure S2 and Supplementary Video S1), and 86% had disappeared by 28 days postinjection, consistent with a rapid die-off. Furthermore, we found that the red fluorescence in Ai35 mice, which was due only to the mCherry expressed by the virus, was much dimmer than the red fluorescence in Ai14 mice at the same time point of seven days postinjection and with the same imaging parameters (see Supplementary Figure S3): the mean intensity was 45.86 (arbitrary units, or “a.u.”) in Ai14 but only 16.29 a.u. in Ai35. This is consistent with the published findings that tdTomato is a much brighter fluorophore than mCherry (42), particularly with two-photon excitation (43), and it is also consistent with Ciabatti et al.’s addition of a destabilization domain to mCherry’s C-terminus. We therefore redid the counts of labeled cells in our Ai14 datasets to include only cells with fluorescence at seven days of more than 32.33 a.u., the midpoint of the mean intensities in Ai35 versus Ai14 mice, in order to exclude neurons that might have been labeled with mCherry alone. As seen in Supplementary Figure S4, restricting the analysis to the cells that were brightest at seven days (and therefore almost certainly not labeled with just mCherry instead of either just tdTomato or a combination of both mCherry and tdTomato) made no major difference: 70.0% of SiR-labeled neurons had disappeared by 14 days, and 80.8% were gone by 21 days.

Although in theory it is possible that the disappearance of the infected cells could be due to cessation of tdTomato or Arch-EGFP-ER2 expression rather than to the cells’ deaths, because of downregulation by rabies virus of host cell gene expression (44), we view this as highly unlikely. Downregulation of host cell gene expression by rabies virus is neither total (“cells with high expression of RbV transcripts retain sufficient transcriptional information for their classification into a specific cell type.” (44)) nor uniform (45); in practice, we saw no evidence of a decline in reporter expression in the infected cells but in fact found the exact opposite. As can be seen in a number of cells in Figures 3 and S4, the cells got brighter and brighter over time, unless they abruptly disappeared. In our experience, including in this case, cells infected with rabies virus increase in brightness until they die, often blebbing and coming apart into brightly labeled pieces, regardless of whether the fluorophore is expressed from a reporter allele (as in this case) or directly by the virus (see Chatterjee et al. 2018 for many more examples of this (21)).

### Construction and testing of a virus with an intact PEST domain

We decided to directly test whether a rabies virus with an intact PEST domain fused to its nucleoprotein can spread transsynaptically, with or without TEVP (Figure 4). Beginning with our lab’s first-generation virus RVΔG-4Cre (21), we constructed a “self-inactivating” version, “RVΔG- NPEST-4Cre”, by adding the coding sequence for the C-terminal addition from Ciabatti et al. (25) to the 3’ end of the nucleoprotein gene. To reduce the chance of the PEST domain being lost to nonsense mutations during production of the virus, we made synonymous changes to five codons near the junction of the end of the native nucleoprotein gene and the beginning of the addition, so that those codons were no longer a single mutation away from being stop codons; apart from these five synonymous changes, the nucleotide sequence of the addition was identical to that used by Ciabatti et al. (25) (Figure 4A). We also made a matched “revertant” version, “RVΔG- N*PEST-4Cre”, with exactly the same sequence as RVΔG-NPEST-4Cre except with a stop codon three codons into the linker, at the same location as the stop codon that we had found in the overwhelming majority of viral particles in the original SiR-CRE. While the genomes of these two new viruses therefore only differed from each other by one codon, at the protein level one virus (RVΔG-NPEST-4Cre) had a full-length PEST domain on the end of its nucleoprotein, while the other (RVΔG-N*PEST-4Cre) had only a two-amino-acid (Gly-Ser) addition to the end of its nucleoprotein and otherwise was an ordinary first-generation ΔG virus.

**Figure 4:**
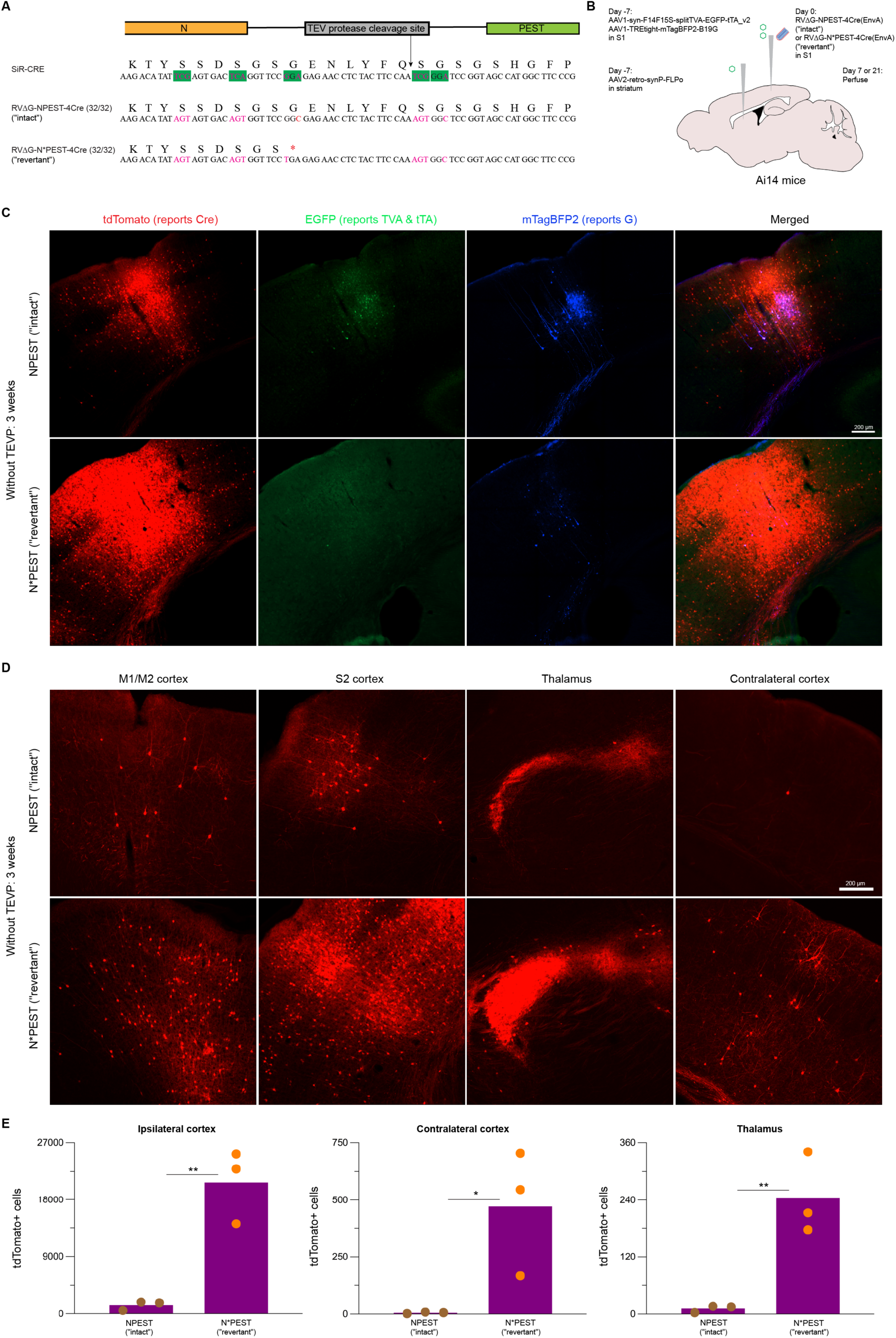
In the absence of TEVP, rabies virus with an apparently intact PEST domain spreads much less than one without a PEST domain. A) Sequences of the region of the junction between the nucleoprotein gene N and its 3’ addition in SiR-CRE (intended sequence) as well as in our two new viruses RVΔG-NPEST-4Cre and RVΔG-N*PEST-4Cre. Apart from synonymous mutations we made in five codons (highlighted in green) to make them less likely to mutate to stop codons, RVΔG-NPEST-4Cre has exactly the same 3’ addition to N as was intended for SiR. RVΔG-N*PEST-4Cre has exactly the same sequence as RVΔG-NPEST-4Cre apart from a stop codon that we deliberately introduced at the same location as the nonsense mutation that we had found in almost every SiR-CRE virion, so that the only modification to the native nucleoprotein is an additional two amino acids (Gly-Ser) on its C-terminus. Sanger sequencing of 32 viral particles of each virus confirmed that the intended additions were still present in the final stocks. B) Diagram of virus injections for monosynaptic tracing experiments. An AAV2-retro expressing FLPo was injected in dorsolateral striatum, and a FLP-dependent helper virus combination was injected into barrel cortex to express G in corticostriatal cells. Rabies virus was injected seven days later at the same location in barrel cortex, and mice were perfused either 7 days or 21 days after rabies virus injection. C-D) Results at 21 days after injection (see Supplementary Fig. S6 for similar results at 7 days). At 21 days after injection, the NPEST virus (top row), with an intact PEST domain, shows very limited spread, whereas the N*PEST virus (bottom row), without the PEST domain, has labeled many thousands of neurons in ipsilateral cortex (C) and also spread to many cells in contralateral cortex and thalamus (D) (see quantification in E). For all panels in this figure as well as Figure 5 and the related supplementary figures, the red channel shows tdTomato, reporting Cre expression from the rabies viruses; the green channel shows EGFP, coexpressed with tTA by the first helper virus; the blue channel shows mTagBFP2, coexpressed with G by the second helper virus. Scale bar: 200 µm, applies to all images. E) Counts of labeled neurons in ipsilateral cortex (left), contralateral cortex (center), and thalamus (right) at 21 days for the two viruses. Each dot indicates the total number of labeled cells found in a given mouse brain when examining every sixth 50 µm section, so that the corresponding numbers in the entire brain would be roughly six times as high (see Methods). All differences in numbers of cells labeled by the two viruses are significant for all conditions (see Supplementary File S8 for all counts and statistical analyses).

Following production of high-titer EnvA-enveloped virus (see Methods), we confirmed that the final stocks retained the intended 3’ additions to the nucleoprotein gene by extracting the genomic RNA and used Sanger sequencing on the genomes of 32 viral particles for each virus. All of the sequenced clones for each of the two viruses had the respective intended sequences (Figure 4, panel A; details in Supplementary File S7), except for one clone of RVΔG-NPEST- 4Cre(EnvA) with a point mutation in the transcriptional termination signal of the nucleoprotein gene (from CATGAAAAAAA to CATAAAAAAAA), and two clones of RVΔG-N*PEST-4Cre(EnvA) that had one synonymous mutation each, one in the nucleoprotein gene and the other in the PEST domain after the introduced stop codon (and that was therefore irrelevant).

Having verified that the NPEST and N*PEST viruses retained their respective modifications (at least in all 32 of the clones sequenced for each), we tested their ability to spread transsynaptically *in vivo* in the absence of TEVP, using corticostriatal neurons as the starting cells (Figure 4, panel B). We injected an AAV2-retro (11) expressing FLPo into the dorsolateral striatum of three mice each of the tdTomato reporter line Ai14(46), with a FLP-dependent cocktail of two helper viruses (AAV1) injected into the primary somatosensory cortex (barrel field) in the same surgery. The helper virus cocktail was a FLP-dependent version of a combination we have published elsewhere (47–49), with the sole modifications being the substitution of orthogonal FRT sites for orthogonal lox ones, and the incorporation of a 2-bp frameshift to prevent the production of a premature stop codon within the first FRT site, in the first helper virus (Supplementary Figure S5 shows the results of pilot testing of this FLP-dependent helper virus). Corticostriatal neurons coinfected with all three AAVs were therefore intended to express, from the first helper virus, TVA (to allow infection by EnvA-enveloped RV), the tetracycline transactivator (tTA), and EGFP (to mark expression of TVA and tTA), as well as, from the second helper virus, G and the blue fluorophore mTagBFP2 (to mark G expression). One week after AAV injection, we injected either the NPEST or the N*PEST virus at equalized titers, then perfused the mice either 7 days or 3 weeks after the rabies virus injections.

Figure 4 and Supplementary Figure S6 show the results of these injections. The “revertant” N*PEST virus, which had only an additional two amino acids on the C-terminus of its nucleoprotein, spread efficiently (bottom rows of images in panels C and D, and rightmost bar in each pair in the charts in panel E). We counted tdTomato-labeled neurons in ipsilateral cortex (regardless of cortical area) as well as in thalamus and in contralateral cortex (regardless of cortical area). Note that, because we counted neurons only in every sixth 50 µm section (see Methods), the numbers of labeled neurons in the entire brain of each mouse would be approximately six times the numbers given below and in the figures. At 7 days after rabies virus injection (Supplementary Figure S6), we found an average of 4751 tdTomato-labeled neurons in ipsilateral cortex (all areas); we also found an average of 73 labeled neurons in contralateral cortex and 321 in thalamus. At 3 weeks after rabies virus injection (Figure 4), a time point which is unusually long for a first-generation virus (21) but which we included to match the duration used by Ciabatti et al. (25), the N*PEST virus had spread to many more neurons: on average 20759 in ipsilateral cortex, 472 in contralateral cortex, and 244 in thalamus. In order to ensure that all this label was not due simply to leaky TVA expression or residual RVG-enveloped virus, we performed control experiments in which we omitted the AAV expressing G and replaced it by an AAV expressing TEVP. This virus, AAV1-TREtight-H2b-emiRFP670-TEVP, was of the same design as the G-expressing AAV (AAV1-TREtight-mTagBFP2-B19G) but encoded TEVP (S219V mutant) instead of G and an H2b-fused near-infrared fluorescent protein (emiRFP670 (50)) instead of mTagBFP2. These control injections resulted in very little labeling in these input regions (Supplementary Figure S8), suggesting that the large amount of label seen when G was included indicated *bona fide* and efficient transsynaptic spread of the N*PEST virus, consistent with our lab’s prior experience with the parent virus RVΔG-4Cre(EnvA) as well as with other first-generation RVΔG vectors.

In contrast, matched experiments using the NPEST virus, which had the intact PEST domain fused to its nucleoprotein, showed much less transsynaptic spread without TEVP at either time point examined (top rows of images in panels C and D, and leftmost bar in each pair in the charts in panel E). At 7 days (Supplementary Figure S6), we found averages of 1024 labeled cells in ipsilateral cortex (all areas), 2 in contralateral cortex, and 4 in thalamus (although these numbers were much lower than the comparable numbers for the N*PEST virus, the differences at this timepoint were not statistically significant due to high variance and low n (3 mice for each condition). See Supplementary File S8 for all cell counts and results of statistical analyses, and see Supplementary File S9 for a summary table of mean cell counts for all conditions. Single-factor ANOVAs were used for all comparisons). At 3 weeks, we found averages of 1366 labeled cells at in ipsilateral cortex, 6 in contralateral cortex, and 11 in thalamus; these numbers were significantly lower than those obtained with the “revertant” N*PEST virus. Here again, this label in input regions did not appear to be due simply to direct retrograde infection by residual RVG- enveloped virus, because it was almost completely absent in control animals in which the G-expressing AAV was omitted and replaced with one expressing TEVP (Supplementary Figure S8).

Finally, we tested the ability of the two viruses to spread when both TEVP and G were supplied (Figure 5 and Supplementary Figure S7). These experiments were done in exactly the same way as the experiments described above but with both the G-expressing and TEVP- expressing AAVs included in the helper virus mixture.

**Figure 5:**
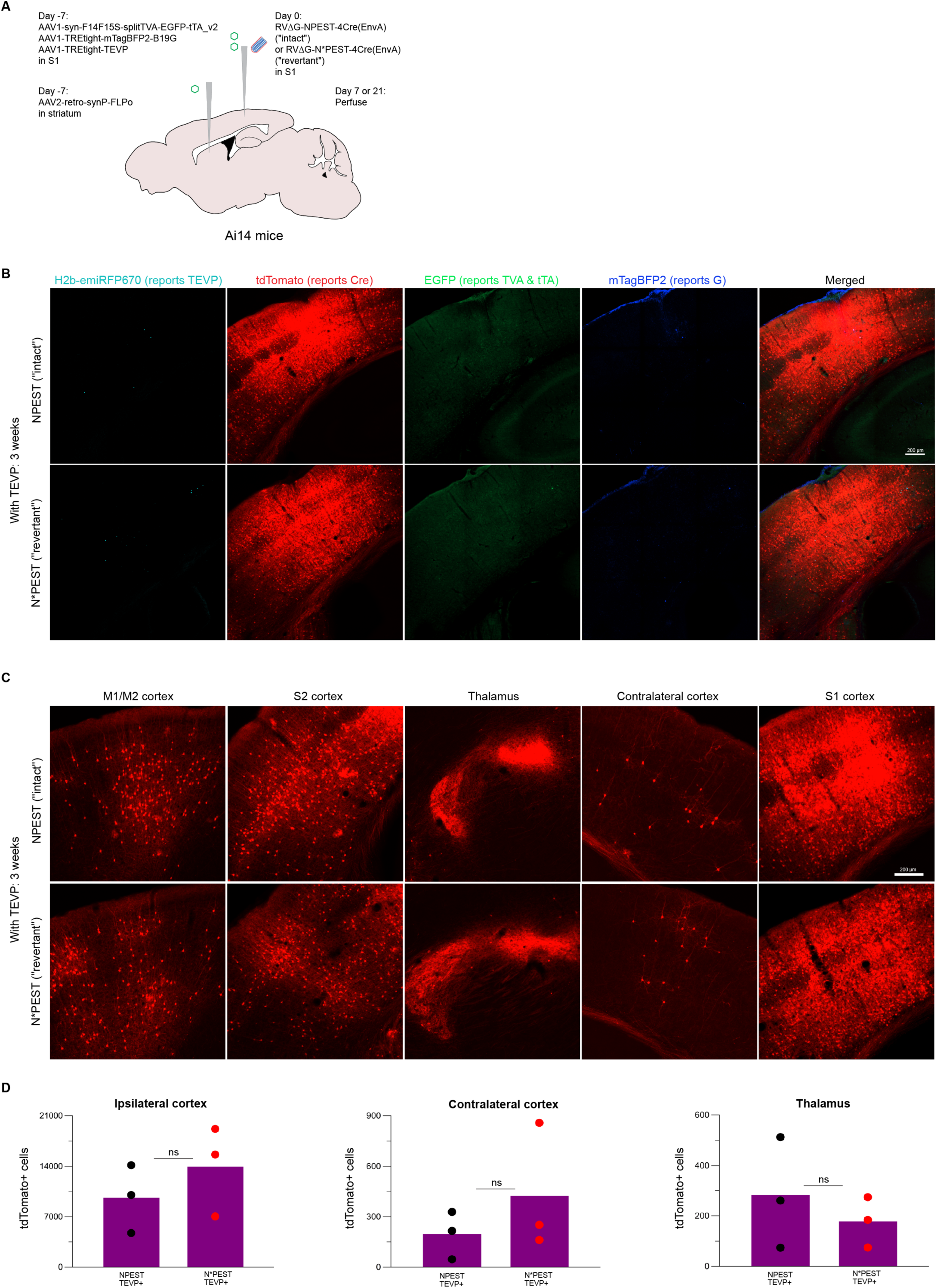
In the presence of TEVP, rabies virus with an apparently intact PEST domain spreads to a similar degree as one without a PEST domain. With the experimental design exactly the same as shown in Figure 4, Panel B but with a TEVP- expressing AAV included in the helper virus mixture (A), the NPEST virus did spread transsynaptically, to a similar degree as the N*PEST one. B-D) Results at 21 days after injection (see Supplementary Fig. S7 for similar results at 7 days). 21 days after injection, both NPEST (top row) and N*PEST (bottom row) viruses have labeled many cells in ipsilateral cortex (B). Both viruses have also spread to contralateral cortex and thalamus (C). D) Counts of tdTomato+ cells in ipsilateral cortex, contralateral cortex, and thalamus. Each dot indicates the total number of labeled cells found in a given mouse brain when examining every sixth 50 µm section, so that the corresponding numbers in the entire brain would be roughly six times as high (see Methods). Differences between numbers of cells labeled by the NPEST and N*PEST viruses are not significant (see Supplementary File S8 for all counts and statistical analyses). Note that the inclusion of TEVP reduced the spread of the N*PEST virus (cf. Figure 4; all counts and statistical comparisons are in Supplementary File S8), possibly because of competition for receptors or expression machinery in source cells.

Whereas the inclusion of TEVP did not much change the results using the “revertant” N*PEST virus, it greatly increased the spread of the NPEST virus, with means at 7 days of 8802 labeled cells in ipsilateral cortex, 82 in contralateral cortex, and 191 in thalamus; at 3 weeks, there were on average 9652 in ipsilateral cortex, 197 in contralateral cortex, and 283 in thalamus. These numbers were all not statistically significantly different from the corresponding ones for the N*PEST virus. These results suggest that the inclusion of TEVP restored the ability of the NPEST virus to replicate and spread, as we assume that the original authors had initially intended.

Importantly, we also found indications that both viruses eventually killed most of the “source cells” in which they replicated (Supplementary Figure S9). For example, using the NPEST virus with TEVP provided, the average number of remaining “source cells” (defined in this case as cells that were labeled by the virus and expressed both G and TEVP) dropped from 29 at 7 days to 2 at 3 weeks (although here again the comparison was not quite statistically significant (p = 0.071820149) due to the high variance and low number (3) of mice per group).

## DISCUSSION

Our transsynaptic tracing results using virus with an intact PEST domain contradict the earlier claim that such a virus can efficiently spread between neurons in the absence of TEVP(25). Our findings from sequencing the SiR viruses from the originating laboratory suggest that the reason for that claim was that the viruses used in the original study had lost the intended modification.

While it is possible that the escape of the two SiR virus samples from the modification intended to attenuate them was a fluke due to bad luck with those two batches, we view this as unlikely, for three reasons. First, the reported finding that viral replication and spread occurred in the absence of TEVP is difficult to understand in the absence of mutations but is easily explained if the viral preparations used for those experiments harbored the kind of mutations that we found in the two preparations to which we had access. Second, the two SiR virus samples that we analyzed had independently developed mutations causing loss of the intended C-terminal addition to the nucleoprotein; we know that the mutations were independent because the two samples were of different viruses so did not both derive from a single compromised parental stock. Third, the mutation profiles of the two viruses were very different: whereas the SiR-CRE sample had the same point mutation in nearly 100% of its viral particles, only a minority of the SiR-FLPo particles had that particular mutation, with the majority having a different point mutation three codons away that had the same result. This suggests that any of the many opportunities for removing the C-terminal addition — creation of a premature stop codon at any one of a number of sites, or a frameshift mutation anywhere in the vicinity — can be exploited by a given batch of virus, greatly increasing the probability of such mutants arising.

While it is clearly possible to make virus with the intended modification to the nucleoprotein in at least the majority of virions, because we have done so here (and the authors of the original paper also report doing so, in a recent preprint (51) in response to our own preprint of an earlier version of this paper), our findings suggest that the approach in its current form is vulnerable to being undermined by viral mutation. Consistent with this, the original authors’ recent preprint reports loss of the 3’ addition in a significant minority of virions within as few as four passages(51). Although a number of groups have made recombinant rabies viruses — as well as other rhabdoviruses and other nonsegmented negative-strand RNA viruses — encoding fusions of exogenous proteins to viral proteins (52–62), most of these groups have found that the additions significantly impaired function, and some have found that the viruses rapidly lost C-terminal additions to viral proteins. For example, an attempt to make SAD B19 rabies virus with EGFP fused to the C-terminus of the nucleoprotein was unsuccessful, suggesting that large C-terminal additions make the nucleoprotein dysfunctional; the authors of that paper resorted instead to making virus encoding the fusion protein in addition to the wild-type nucleoprotein (55). A vesicular stomatitis virus with GFP fused to the C-terminus of the glycoprotein gene lost the modification within a single passage of the virus because of a point mutation creating a premature stop codon (52). Relatedly, a VSV with its glycoprotein gene replaced with that of a different virus was found to quickly develop a premature stop codon causing loss of the last 21 amino acids of the exogenous glycoprotein, conferring a marked replication advantage to the mutants bearing the truncated version (63). Generalizing from these prior examples as well as our findings here, we suggest that any attempt to attenuate a virus by addition to the C-terminus of a viral protein will be vulnerable to loss of the modification, and that any such virus will therefore need to be monitored very carefully.

It is unclear whether further improvements to the design of the viruses can be made to make loss of the PEST domain less likely. Although the synonymous mutations that we made to five codons near the junction of N and the 3’ addition are a good start, there are numerous other codons in the immediate vicinity that are also one point mutation away from being stop codons but that do not have synonyms that are not. Furthermore, no such changes would protect against frameshifts.

If the viruses used for the transsynaptic tracing experiments in the original paper were actually *de facto* first-generation, ΔG viruses like the SiR samples that we analyzed, how could the authors have found, in postmortem tissue, cells labeled by SiR-CRE that had survived for weeks? The answer may simply be that, as we have shown in Chatterjee et al. (21) and again here (Figure 3), and as the original authors also report in their new preprint (51), a preparation of first-generation rabies viral vector expressing Cre can leave a large fraction of labeled cells alive for at least months, in contrast to similar ones encoding tdTomato (21) or EGFP (4). Similarly, Gomme et al. found long-term survival of some neurons following infection by a replication-competent rabies virus expressing Cre (64)). Our results with the “revertant” control virus RVΔG- N*PEST-4Cre (Figures 4 and 5, and Supplementary Figures S5-S7), showing thousands of labeled neurons three weeks after rabies virus injection, are consistent with this as well.

One reason, in turn, why a preparation of a simple ΔG rabies virus encoding Cre can leave many cells alive may be that not all the virions are in fact first-generation viral particles, because of the high mutation rate that we have highlighted in this paper. We have shown in Chatterjee et al. (21) that a second-generation (ΔGL) rabies virus, which has both its glycoprotein gene G and its polymerase gene L deleted, leaves cells alive for the entire four months that we followed them. However, any first-generation (ΔG) virus that contains a frameshift or point mutation knocking out L will in practice be a ΔGL virus. In fact, a stop codon or frameshift mutation in any of several other viral genes is likely to have a similar effect as one in L (and it might be that the Ser419Thr mutation that we found in 9.49% of our RVΔG-4Cre virions is just such a knockout mutation of N). Together with the high mutation rate of rabies virus, this means that, within every preparation of first-generation rabies virus there is almost guaranteed to be a population of de facto second-generation variants mixed in with the intended first-generation population and propagated in the producer cells by complementation by the first-generation virions. Any rabies virus preparation (whether made in the laboratory or occurring naturally) can be expected to contain a population of such knockout (whether by substitution, frameshift, or deletion) mutants (related to the classic phenomenon of “defective interfering particles”, or mutants with a marked replication advantage (65–67), and the higher the multiplicity of infection when passaging the virus, the higher the proportion of such freeloading viral particles typically will be. This would not necessarily be noticed in the case of a virus encoding a more common transgene product such as a fluorophore, because the expression levels of these by the knockout mutants would be too low to label cells clearly (see Figure 1 in Chatterjee et al. (21)). However, with Cre as the payload, any “second-generation” particles would be able to label neurons but not kill them, because second-generation rabies viral vectors do not kill cells for at least months (21). This explanation would predict that the percentage of neurons surviving infection with a rabies virus encoding Cre will depend on the particular viral preparation that is injected, with some having a greater fraction of knockout particles than others.

This could explain why the SiR-CRE virus sample killed cells faster than our own RVΔG-4Cre (Figure 3). This analysis would also presumably apply to first-generation (ΔG) viruses expressing FLPo: while we found that the FLPo-expressing version that we made did not leave as many cells alive as the Cre-expressing version did (Supplementary Figure S10), that preparation may simply have had fewer mutants with knockout of genes essential for replication. Most interestingly, we found that virus with an intact PEST domain does spread efficiently between neurons when TEVP is supplied. This suggests that such viruses could become the basis for monosynaptic tracing systems with reduced toxicity after all. Our finding is complementary to those of the original authors in their recent preprint (51) that an intact SiR virus did not kill labeled neurons for five months: that study was of neurons only infected by the rabies virus (i.e., without expression of G to complement the ΔG virus or expression of TEVP to remove the C-terminal addition to the nucleoprotein); they did not show that their intact virus could spread between neurons or that it is nontoxic as it does so. Conversely, we showed here that an intact NPEST virus can spread between neurons, but we did not specifically investigate toxicity during this process. It remains to be seen to what degree an NPEST or SiR virus that spreads transsynaptically is cytotoxic, not only to transsynaptically-labeled cells but also to the starting cells, which need to express both G and TEVP to allow replication and spread of the virus. This is not a trivial point, as G is toxic when overexpressed (68), and the original authors’ finding that cultured cells rapidly lost TEVP activity (51) suggests that TEVP expressed at sufficient levels may be toxic as well. Indeed, the much lower average numbers of source cells that we found at 3 weeks than at 7 days (Supplementary Figure S9) suggest that the NPEST virus in the presence of TEVP may be just as cytotoxic to source cells as ordinary ΔG virus is.

However, there may be room for improvement: the TEVP expression levels that we provided here are unlikely to happen to be optimal. Given that we have used the tTA/TRE system to provide both TEVP and G in the source cells, the expression of both could be controlled with doxycycline to result in a system that is less toxic to source cells (mirroring an approach we have used with ΔGL viruses (24). Another consideration (perhaps minor) is the nine additional amino acids that are unavoidably left on the C-terminus of the nucleoprotein after the rest of the addition is removed by TEVP: while these additional amino acids may not much impair the function of the protein, they are unlikely to help it.

It is also unclear to what degree a minority population of revertant mutants that do arise in stocks of otherwise-intact virus would pose a problem for monosynaptic tracing studies. If, for example, ∼5% of the virions in a given preparation were revertant mutants (e.g., as the original authors report obtaining after six passages in cells highly expressing TEVP (51)), one might expect the same percentage of labeled presynaptic neurons to be labeled by those mutants and therefore to experience the toxicity of infection by a first-generation, ΔG virus. However, because the process of infecting the source cells, replicating within them (if provided with G), and spreading to other cells is comparable to an additional passage in cell culture, the percentage of presynaptic cells labeled by the revertant mutants could be higher, and would presumably depend on the level of TEVP expression in each source cell.

Such a minority population of revertant mutants in our “intact” NPEST virus preparation may explain the limited spread of this virus that we observed in the absence of TEVP. Although all 32 of the clones that we checked using Sanger sequencing had the intended 3’ modification to the nucleoprotein gene (Supplementary Figure S7), it is entirely possible (perhaps even inevitable) that revertant virions were present in lower numbers, possibly at a level sufficient to support some replication of the mixed population of virions in cells expressing G without TEVP. While we view this as the most likely explanation, showing this convincingly would be difficult, even assuming that high-throughput sequencing (such as the SMRT sequencing we used to analyze the originating lab’s stocks) would reveal the presence of revertant mutants at some level. Even *in situ* sequencing sequencing(69–71) of viral genomes in pre- and postsynaptic neurons in mice injected with the “intact” NPEST virus might not be conclusive: because functional nucleoprotein produced in postsynaptic cells by revertant virus would complement and package “intact” NPEST-encoding virions, allowing them to spread, it would not be particularly surprising to find that the majority of virions in presynaptic neurons were “intact”, even if a minority population of revertant mutants were necessary to allow any spread at all in the absence of TEVP. This is speculation, however.

What seems clear from the results we have presented here is that reliance on addition to a viral protein of a C-terminal domain that could be lost by mutation at any replication event is a tenuous basis for rendering a virus nontoxic. At the very least, we would suggest that any such attenuating addition be placed on the N terminus of the viral protein (if this turns out to be possible without impairing the function or trafficking of the protein) so that it could not easily be mutated away.

In summary, our results suggest that rabies virus with a PEST domain added to the C terminus of its nucleoprotein is vulnerable to the loss of that modification and only spreads efficiently between neurons if it is removed, whether by unintended mutation or by expression of TEVP. Our finding that virus with a PEST domain that is intact in the majority of virions can spread efficiently when TEVP is provided, as the designers had presumably originally intended, raises the possibility that further optimization and validation could make the SiR approach a viable option for monosynaptic tracing with reduced toxicity.

## SUMMARY OF METHODS (see Supplementary Methods for details)

### Cloning

The following novel plasmids were made using standard cloning techniques (see Supplementary Methods): pLV-CAG-FLEX-BFP-(mCherry)’ (Addgene 115234), pLV-CAG-F14F15S-BFP-(mCherry)’ (Addgene 115235), pLV-U-TVA950 (Addgene 115236), pRVΔG-4FLPo (Addgene 122050), pAAV-syn-F14F15S-splitTVA-EGFP-tTA_v2 (Addgene 183352), pB-CAG-TEVP-IRES-mCherry (Addgene 174377), pAAV-syn-FLPo (Addgene 174378), pAAV-TREtight-H2b- emiRFP670-TEVP (Addgene 174379), pRVΔG-NPEST-4Cre (Addgene 174380), pRVΔG- N*PEST-4Cre (Addgene 174381), pCAG-hypBase. All of the above novel plasmids have been deposited with Addgene and can be obtained from there, except for pCAG-hypBase, the distribution of which is not permitted due to intellectual property constraints.

### Production of lentiviral and adeno-associated viral vectors

Lentiviral vectors were made as described (72) but using a vesicular stomatitis virus envelope expression plasmid pMD2.G for most vectors except for LV-U-TVA950(B19G), which was made using the rabies virus envelope expression plasmid pCAG-B19GVSVGCD (72).

AAV1-syn-F14F15S-splitTVA-EGFP-tTA_v2, AAV1-TREtight-H2b-emiRFP670-P2A-TEVP, and AAV2-retro-syn-FLPo were made in-house by standard techniques (see Supplementary Methods).

AAV1-TREtight-mTagBFP2-B19G (which we have described previously(47, 48)), was packaged as serotype 1 by Addgene (catalog # 100798-AAV1).

### Production of titering cell lines

Reporter cell lines 293T-FLEX-BC and 293T-F14F15S-BC were made using lentiviral vectors made from pLV-CAG-FLEX-BFP-(mCherry)’ and pLV-CAG-F14F15S-BFP-(mCherry)’, described above. TVA-expressing versions, 293T-FLEX-BC-TVA and 293T-F14F15S-BC-TVA, were made by infecting the above lines with LV-U-TVA950(VSVG) (described above).

### Production of TEVP-expressing cell line

293T-TEVP was made by transfecting HEK 293T/17 cells with pCAG-hypBase and pB-CAG- TEVP-IRES-mCherry, then sorting.

### Production and titering of rabies viruses

RVΔG-4Cre, RVΔGL-4Cre, RVΔG-NPEST-4Cre, and RVΔG-N*PEST-4Cre were produced mostly as described (21, 35) (see Supplementary Methods); titering was as described (33) but using the 293T-FLEX-BC and 293T-F14F15S-BC lines used for B19G-enveloped viruses and the 293T-FLEX-BC-TVA and 293T-F14F15S-BC-TVA used for the EnvA-enveloped viruses.

### Extraction of viral genomic RNA and preparation for Sanger sequencing

RNA viral genomes were extracted from virus samples using a Nucleospin RNA kit (Macherey-Nagel, Germany), then converted to cDNA by RT-PCR (Agilent Technologies, USA) with a barcoded primer. cDNA sequences were amplified using Platinum SuperFi Green Master Mix (Invitrogen (Thermo Fisher), USA) and cloned into pEX-A (Eurofins Genomics, USA) using an In- Fusion HD Cloning Kit (Takara Bio, Japan). Sequencing data was collected for over fifty clones per sample.

### Single-molecule, real-time (SMRT) sequencing

Double-stranded DNA samples for SMRT sequencing were prepared similarly to the above, except that that the clones generated from each of the three virus samples were tagged with one of the standard PacBio barcode sequences to allow identification of each clone’s sample of origin following multiplex sequencing. This was in addition to the random index (10 nucleotides in this case) that was again included in the RT primers in order to uniquely tag each individual genome.

### Surgeries and virus injections for two-photon imaging

All experimental procedures using mice were conducted according to NIH guidelines and were approved by the MIT Committee for Animal Care (CAC). Mice were housed 1-4 per cage under a normal light/dark cycle for all experiments.

Adult mice of Cre-dependent reporter strains Ai14 (73) (Jackson Laboratory #007908) or Ai35D (41) (Jackson Laboratory # 012735) mice were injected in V1 with LV-U-TVA950(B19G), then implanted with a glass window. Seven days later, windows were removed and one of the three EnvA-enveloped rabies viral vectors (with equalized titers) was injected at the same coordinates, then coverslips were reapplied.

### *In vivo* two-photon imaging and image analysis

Beginning seven days after injection of each rabies virus and continuing every seven days up to a maximum of four weeks following rabies virus injection, the injection sites were imaged on a two-photon microscope. One field of view was chosen in each mouse in the area of maximal fluorescent labelling. Cell counting was performed with the ImageJ Cell Counter plugin.

### Monosynaptic tracing experiments: surgeries and virus injections

The three helper AAV1s were combined at final titers of 3.6E10 gc/ml for AAV1-syn-F14F15S- splitTVA-EGFP-tTA_v2 and 6.60E11 gc/ml for AAV1-TREtight-mTagBFP2-B19G and/or AAV1- TREtight-H2b-emiRFP670-P2A-TEVP. 250 nl of helper virus mixture was injected into layer 5 of barrel cortex of Ai14 mice; in the same surgery, 300 nl of AAV2-retro-syn-FLPo (1.16E13 gc/ml) was injected into dorsolateral striatum. 7 days after AAV injection, 300nl of RVΔG-NPEST- 4Cre(EnvA) (1.86E9 iu/ml) or RVΔG-N*PEST-4Cre(EnvA) (diluted to 1.86E9 iu/ml) was injected in barrel cortex at the same site as the helper AAV mixtures.

### Monosynaptic tracing experiments: perfusions and histology

12 days or 3 weeks after injection of rabies virus, mice were perfused; Brains were postfixed overnight and cut into 50 µm coronal sections on a vibrating microtome. Sections were immunostained as described(48) with a chicken anti-GFP (Aves Labs GFP-1020) 1:500 and donkey anti-chicken Alexa Fluor 488 (Jackson Immuno 703-545-155) 1:200.

### Monosynaptic tracing experiments: cell counts and microscopy

tdTomato-labeled neurons in contralateral cortex and thalamus were counted manually with the Cell Counter plugin in ImageJ. Cells at the injection site were counted either manually or (when dense) using the Analyze Particle function in ImageJ. Only one of the six series of sections (i.e., every sixth section: see above) was counted for each mouse. Images for figures were taken on a confocal microscope (Zeiss, LSM 900).

## ACKNOWLEDGEMENTS

We thank Ernesto Ciabatti and Marco Tripodi for sharing samples of EnvA/SiR-CRE and EnvA/SiR-FLPo and for comments on the manuscript. We thank Ed Callaway, Sean Whelan, Ayano Matsushima, and Kim Ritola for helpful discussion and Jun Zhuang, Soumya Chatterjee, and Ali Cetin for helpful discussion and sharing their own results with SiR viruses. We thank Stuart Levine, Noelani Kamelamela, and Huiming Ding of the MIT BioMicro Center for assistance with SMRT sequencing and bioinformatic data analysis and Sara Beach for helpful feedback on the manuscript. We thank Xin Fu for noticing the stop codon that I.R.W. had inadvertently introduced into the FLP-dependent helper virus plasmid pAAV-F14F15S-splitTVA-EGFP-tTA. Research reported in this publication was supported by the following BRAIN Initiative awards from the National Institute of Mental Health: U01MH106018 (Wickersham) RF1MH120017 (Wickersham), U01MH114829 (Dong), and U19MH114830 (Zeng).

## SUPPLEMENTARY METHODS

### Cloning

Lentiviral transfer plasmids were made by cloning, into pCSC-SP-PW-GFP (1) (Addgene #12337), the following components:

the CAG promoter (2) and a Cre-dependent “FLEX” (3) construct consisting of pairs of orthogonal lox sites flanking a back-to-back fusion of the gene for mTagBFP2 (4) immediately followed by the reverse-complemented gene for mCherry (5), to make the Cre reporter construct pLV-CAG-FLEX-BFP-(mCherry)’ (Addgene 115234);

the CAG promoter (2) and a Flp-dependent “FLEX” (3) construct consisting of pairs of orthogonal FRT sites (6) flanking a back-to-back fusion of the gene for mTagBFP2 (4) immediately followed by the reverse-complemented gene for mCherry (5), to make the Flp reporter construct pLV-CAG-F14F15S-BFP-(mCherry)’ (Addgene 115235);

the ubiquitin C promoter from pUB-GFP (7) (Addgene 11155) and the long isoform of TVA (8) to make the TVA expression vector pLV-U-TVA950 (Addgene 115236).

The first-generation vector genome plasmid pRVΔG-4FLPo (Addgene 122050) was made by cloning the FLPo gene (9) into pRVΔG-4Cre.

pAAV-syn-F14F15S-splitTVA-EGFP-tTA_v2 (Addgene 183352) is a FLP-dependent version of the Cre-dependent helper virus genome plasmid pAAV-syn-FLEX-splitTVA-EGFP-tTA (Addgene 52473) with orthogonal FRT sites(6) instead of orthogonal lox sites, and with a 2-bp frameshift to avoid creation of a premature stop codon in the first FRT site. Note that this plasmid is a replacement for pAAV-syn-F14F15S-splitTVA-EGFP-tTA, which was used in an earlier preprint of this manuscript.

pCAG-hypBase was made by synthesizing the 1785-bp gene for an improved version of piggyBac transposase (10) and cloning it into the EcoRI and NotI sites of pCAG-GFP (7) (Addgene 11150).

pB-CAG-TEVP-IRES-mCherry (Addgene 174377) was made by cloning the CAG promoter (2), a mammalian codon-optimized version (11, 12) of the TEVP gene (S219V mutant) (13), the EMCV IRES (14), and the mCherry gene (5) into pB-CMV-MCS-EF1-Puro (System Biosciences #PB510B-1).

pAAV-syn-FLPo (Addgene 174378) was made by cloning the FLPo gene(9) into the EcoRI and AccIII sites of pAAV-syn-FLEX-EGFP-B19G (Addgene 59333).

pAAV-TREtight-H2b-emiRFP670-TEVP (Addgene 174379) was made by cloning an H2b-emiRFP670 fusion gene (15) (sequence from Addgene 136571 but with the internal kozak sequence replaced by a short GSG linker to prevent translation of fluorophore unfused to H2b), followed by a P2A sequence and the above-described TEVP gene, into the EcoRI and NheI sites of pAAV-TREtight-mTagBFP2-B19G (16) (Addgene 100799).

pRVΔG-NPEST-4Cre (Addgene 174380) was made by cloning a synthesized fragment containing the 3’ addition from Ciabatti et al. ’17, with synonymous changes to 5 codons (17) in the immediate vicinity of the junction between the nucleoprotein gene and the 3’ addition so that more than a single point mutation would be required to convert them to stop codons, into the PmlI and BstI sites of pRVΔG-4Cre(18) (Addgene 98034).

pRVΔG-N*PEST-4Cre (Addgene 174381) was made identically to the above except that the glycine codon in position 453 was replaced with a stop codon (TGA).

All of the above novel plasmids have been deposited with Addgene, with the accession numbers given above, and can be purchased from there except for pCAG-hypBase, the distribution of which is not permitted due to intellectual property constraints.

### Production of lentiviral and adeno-associated viral vectors

Lentiviral vectors were made by transfection of HEK-293T/17 cells (ATCC 11268) as described (19) but using the vesicular stomatitis virus envelope expression plasmid pMD2.G (Addgene 12259) for all vectors except for LV-U-TVA950(B19G), which was made using the rabies virus envelope expression plasmid pCAG- B19GVSVGCD (19). Lentiviral vectors expressing fluorophores were titered as described (20); titers of LV-U- TVA950(VSVG) and LV-U-TVA950(B19G) were assumed to be approximately the same as those of the fluorophore-expressing lentiviral vectors produced in parallel.

AAV1-TREtight-mTagBFP2-B19G (which we have described previously(16, 21)), was packaged as serotype 1 by Addgene (catalog # 100798-AAV1).

AAV1-syn-F14F15S-splitTVA-EGFP-tTA_v2, AAV1-TREtight-H2b-emiRFP670-P2A-TEVP and AAV2-retro-syn-FLPo were made by transfecting HEK 293T/17 cells with the respective genome plasmid along with pHelper (Cellbiolabs VPK-421) and either pAAV-RC1 (Cellbiolabs VPK-421) or “rAAV2-retro helper” (Addgene 81070)(22), using Xfect Transfection Reagent (Takara 631318) according to the manufacturer’s protocol, with collection of supernatant (and replacement with fresh media) at 3 days after transfection and collection of supernatant as well as the transfected cells at 5 days after transfection. Virus was pelleted from supernatants using PEG 8000; cells were lysed by four freeze-thaw cycles. Pelleted virus and cell lysate were pooled and treated with benzonase, purified on an iodixanol gradient, then concentrated in an Amicon Ultra-15 centrifugal filter unit (Millipore Sigma UFC9100).

### Production of titering cell lines

To make reporter cell lines, HEK-293T/17 cells were infected with either pLV-CAG-FLEX-BFP-(mCherry)’ or pLV-CAG-F14F15S-BFP-(mCherry)’ at a multiplicity of infection of 100 in one 24-well plate well each. Cells were expanded to 2x 15cm plates each, then sorted on a FACS Aria to retain the top 10% most brightly blue fluorescent cells. After sorting, cells were expanded again to produce the cell lines 293T-FLEX-BC and 293T- F14F15S-BC, reporter lines for Cre and FLPo activity, respectively. TVA-expressing versions of these two cell lines were made by infecting one 24-well plate well each with LV-U-TVA950(VSVG) at an MOI of approximately 100; these cells were expanded to produce the cell lines 293T-FLEX-BC-TVA and 293T-F14F15S-BC-TVA.

### Production of TEVP-expressing cell line

The 293T-TEVP cell line was made by transfecting HEK 293T/17 (ATCC CRL-11268) cells with pCAG-hypBase and pB-CAG-TEVP-IRES-mCherry in a 15 cm plate (293T/17) or one well each of a 24 well plate (BHK-B19G2 and BHK-EnvA2) using Lipofectamine 2000 (Thermo Fisher 11668019) according to the manufacturer’s instructions, then expanding and sorting the cells on a FACSAria (BD Biosciences) for the brightest 20% of red fluorescent cells, then expanding and freezing the sorted cells.

### Production and titering of rabies viruses

RVΔG-4Cre and RVΔGL-4Cre were produced as described (18, 23), with EnvA-enveloped viruses made by using cells expressing EnvA instead of G for the last passage. Titering and infection of cell lines with serial dilutions of viruses was as described (24), with the 293T-TVA-FLEX-BC and 293T-TVA-F14F15S-BC lines used for B19G-enveloped viruses and the 293T-TVA-FLEX-BC and 293T-TVA-F14F15S-BC used for the EnvA-enveloped viruses. For the *in vivo* injections, the three EnvA-enveloped, Cre-encoding viruses were titered side by side, and the two higher-titer viruses were diluted so that the final titer of the injected stocks of all three viruses were approximately equal at 1.39E9 infectious units (i.u.) per milliliter.

RVΔG-4mCherry(EnvA) was produced and titered as described (18, 23, 24), with final titer of 1.7E10 i.u./mL on 293T-TVA800 cells and 1.60E6 i.u./mL on HEK-293T cells. RVΔG-4Cre(EnvA) used for Supplementary Figure S5 was produced and titered as described (18, 23, 24), with final titers of 8.58E9 i.u./mL on 293T-TVA800 cells (and 3.14E5 i.u./mL on HEK-293T cells) and was diluted to a titer of 4.49E9 i.u./mL before injection.

The two new rabies viruses RVΔG-NPEST-4Cre and RVΔG-N*PEST-4Cre were produced as described(18, 23) but with only one amplification passage (P1) between the rescue transfection step and the final passage on EnvA-expressing cells. For the NPEST version, the following additional modifications were made: for the rescue transfection, the new cell line 293T-TEVP (see above) was used instead of HEK 293T-17, and pB-CAG-TEVP-IRES-mCherry (10 µg per 15 cm plate) was included in the plasmid mix. Because pilot testing showed no clear advantage of the 293T-TEVP line over transfected 293Ts for passaging, the P1 passage was on HEK 293T/17 cells transfected with equal amounts of pCAG-B19G and pB-CAG-TEVP-IRES-mCherry (32 µg of each plasmid per 15 cm plate). The final passage was on BHK-EnvA2 cells(24) transfected with pB-CAG-TEVP-IRES-mCherry. Lipofectamine 2000 (Thermo Fisher 11668019) was used for all transfections, following the manufacturer’s protocol, except for the final passage of RVΔG-NPEST-4Cre, for which Lipofectamine 3000 (Thermo Fisher L3000-015) was used.

The final stocks of RVΔG-NPEST-4Cre(EnvA) and RVΔG-N*PEST-Cre(EnvA) were titered side by side as described(24) on 293T-FLEX-BC-TVA cells (see above) in two-fold dilution series and on 293T-FLEX-BC cells (see above) undiluted (for determination of the titer of contaminating non-EnvA-enveloped virus). Final titers of the viruses on the TVA-expressing cells were determined to be 1.18E9 i.u./mL for RVΔG-NPEST-4Cre(EnvA) and 2.41E10 i.u./mL (normalized) for RVΔG-N*PEST-4Cre(EnvA). Prior to injection *in vivo* (see below), the N*PEST virus was diluted 20.42-fold in DPBS to match the titer of the NPEST version. Final titers of the viruses on the non-TVA-expressing cells were determined to be 3.79E5 i.u./mL for RVΔG-NPEST-4Cre(EnvA) and 8.28E5 i.u./mL (normalized) for RVΔG-N*PEST-4Cre(EnvA).

### Immunostaining and imaging of cultured cells

Reporter cells (see above) plated on coverslips coated in poly-L-lysine (Sigma) were infected with serial dilutions of RVΔG-4Cre and RVΔGL-4Cre as described (24). Three days after infection, cells were fixed with 2% paraformaldehyde, washed repeatedly with blocking/permeabilization buffer (0.1% Triton-X (Sigma) and 1% bovine serum albumin (Sigma) in PBS), then labeled with a blend of three FITC-conjugated anti-nucleoprotein monoclonal antibodies (Light Diagnostics Rabies DFA Reagent, EMD Millipore 5100) diluted 1:100 in blocking buffer for 30 minutes, followed by further washes in blocking buffer, then finally briefly rinsed with distilled water and air-dried before mounting the coverslips onto microscope slides with Prolong Diamond Antifade (Thermo P36970) mounting medium. Images of wells at comparable multiplicities of infection (∼0.1) were collected on a Zeiss 710 confocal microscope.

### Extraction of viral genomic RNA and preparation for Sanger sequencing

RNA viral genomes were extracted from two Tripodi lab (EnvA/SiR-CRE and EnvA/Sir-FLPo) and three Wickersham lab (RVΔG-4mCherry, RVΔG-NPEST-4Cre(EnvA), and RVΔG-N*PEST-Cre(EnvA)) rabies virus samples using a Nucleospin RNA kit (Macherey-Nagel, Germany) and treated with DNase I (37°C for 1 hour, followed by 70°C for 5 minutes). Extracted RNA genomes were converted to complementary DNA using an AccuScript PfuUltra II RT-PCR kit (Agilent Technologies, USA) at 42°C for 2 hours with the following barcoded (so that individual viral particles’ genomes would be marked with distinct barcodes) primer annealing to the rabies virus leader sequence:

Adapter_N8_leader_fp: TCAGACGATGCGTCATGCNNNNNNNNACGCTTAACAACCAGATC

cDNA sequences from the leader through the first half of the rabies virus P gene were amplified using Platinum SuperFi Green Master Mix (Invitrogen (Thermo Fisher), USA) with cycling conditions as follows: denaturation at 98°C for 30 seconds, followed by 25 cycles of amplification (denaturation at 98°C for 5 seconds and extension at 72°C for 75 seconds), with a final extension at 72°C for 5 minutes, using the following primers: pEX_adapter_fp: CAGCTCAGACGATGCGTCATGC

Barcode2_P_rp: GCAGAGTCATGTATAGCTTCTTGAGCTCTCGGCCAG

The ∼2kb PCR amplicons were extracted from an agarose gel, purified with Nucleospin Gel and PCR Clean-up (Macherey-Nagel, Germany), and cloned into pEX-A (Eurofins Genomics, USA) using an In-Fusion HD Cloning Kit (Takara Bio, Japan). The cloned plasmids were transformed into Stellar competent cells (Takara Bio, Japan), and 200 clones per rabies virus sample were isolated and purified for sequencing. For each clone, the index and the 3’ end of the N gene were sequenced until sequencing data was collected for over fifty clones per sample: 51 clones from SiR-CRE(EnvA), 50 from SiR-FLPo(EnvA), and 51 from RVΔG-4mCherry(EnvA). Although viral samples may contain plasmid DNA, viral mRNA, and positive-sense anti-genomic RNA, this RT-PCR procedure can amplify only the negative-sense RNA genome: the reverse transcription primer is designed to anneal to the leader sequence of the negative-strand genome so that cDNA synthesis can start from the negative-sense RNA genome, with no other possible templates. Additionally, the PCR amplifies the cDNA, not any plasmids which were transfected into producer cell lines during viral vector production, because the forward PCR primer anneals to the primer used in the reverse transcription, rather than any viral sequence. This RT-PCR protocol ensures that only negative-sense RNA rabies viral genomes can be sequenced.

### Sanger sequencing of transgenes in SiR viruses

The procedure for sequencing the transgene inserts was the same as above, but with the RT primer being Adaptor_N8_M_fp (see below), annealing to the M gene and again with a random 8-nucleotide index to tag each clone, and with PCR primers pEX_adaptor_fp (see above) and Barcode2_L_rp (see below), to amplify the sequences from the 3’ end of the M gene to the 5’ end of the L gene, covering the iCre-P2A-mCherryPEST (or FLPo-P2A-mCherryPEST) sequence.

Primers for RT and PCR for Sanger sequencing were as follows:

Adaptor_N8_M_fp:

TCAGACGATGCGTCATGCNNNNNNNNCAACTCCAACCCTTGGGAGCA

Barcode2_L_rp:

GCAGAGTCATGTATAGTTGGGGACAATGGGGGTTCC

### Sanger sequencing analysis of PEST region in SiR and control viruses

Alignment and mutation detection were performed using SnapGene 4.1.9 (GSL Biotech LLC, USA). Reference sequences of the viral samples used in this study were based on deposited plasmids in Addgene: pSAD-F3- NPEST-iCRE-2A-mCherryPEST (Addgene #99608), pSAD-F3-NPEST-FLPo-2A-mCherryPEST (Addgene #99609), and pRVΔG-4mCherry (Addgene #52488). Traces corresponding to indices and mutations listed in Figure 1 and Supplementary File S1 were also manually inspected and confirmed.

### Single-molecule, real-time (SMRT) sequencing

Double-stranded DNA samples for SMRT sequencing were prepared similarly to the above, except that that the clones generated from each of the three virus samples were tagged with one of the standard PacBio barcode sequences to allow identification of each clone’s sample of origin following multiplex sequencing (see https://www.pacb.com/wp-content/uploads/multiplex-target-enrichment-barcoded-multi-kilobase-fragments-probe-based-capture-technologies.pdf and https://github.com/PacificBiosciences/Bioinformatics-Training/wiki/Barcoding-with-SMRT-Analysis-2.3). This was in addition to the random index (10 nucleotides in this case) that was again included in the RT primers in order to uniquely tag each individual genome.

RNA viral genomes were extracted from two Tripodi lab (SiR-CRE and Sir-FLPo) and one Wickersham lab (RVΔG-4Cre (18); see Addgene #98034 for reference sequence) virus samples using a Nucleospin RNA kit (Macherey-Nagel, Germany) and treated with DNase I (37°C for 1 hour, followed by 70° for 5 minutes). Primers for RT and PCR are listed below. PCR cycling conditions were as follows: denaturation at 98°C for 30 seconds, followed by 20 cycles of amplification (denaturation at 98°C for 5 seconds and extension at 72°C for 75 seconds), with a final extension at 72°C for 5 minutes. This left each amplicon with a 16bp barcode at each of its ends that indicated which virus sample it was derived from, in addition to a 10-nt index sequence that was unique to each genome molecule.

Primers for RT and PCR for SMRT sequencing were as follows:

RVΔG-4Cre:

RT:

PCR:

Barcode1_cagc_N10_leader_fp: TCAGACGATGCGTCATCAGCNNNNNNNNNNACGCTTAACAACCAGATC

PCR:

Barcode1_cagc_fp: TCAGACGATGCGTCAT-CAGC

Barcode2_P_rp (see above)

SiR-CRE:

RT:

Barcode5_cagc_N10_leader_fp: ACACGCATGACACACTCAGCNNNNNNNNNNACGCTTAACAACCAGATC

PCR:

Barcode5_cagc_fp: ACACGCATGACACACT-CAGC

Barcode3_P_rp: GAGTGCTACTCTAGTACTTCTTGAGCTCTCGGCCAG

SiR-FLPo:

RT:

Barcode9_cagc_N10_leader_fp: CTGCGTGCTCTACGACCAGCNNNNNNNNNNACGCTTAACAACCAGATC

PCR:

Barcode9_cagc_fp: CTGCGTGCTCTACGAC-CAGC

Barcode4_P_rp: CATGTACTGATACACACTTCTTGAGCTCTCGGCCAG

After the amplicons were extracted and purified from an agarose gel, the three were mixed together at 1:1:1 molar ratio. The amplicons’ sizes were confirmed on the Fragment Analyzer (Agilent Technologies, USA), then hairpin loops were ligated to both ends of the mixed amplicons to make circular SMRTbell templates for Pacbio Sequel sequencing. SMRTbell library preparation used the PacBio Template Preparation Kit v1.0 and Sequel Chemistry v3. Samples were then sequenced on a PacBio Sequel system running Sequel System v.6.0 (Pacific Biosciences, USA), with a 10-hour movie time.

### Bioinformatics for PacBio sequence analysis

For the ∼2kb template, the DNA polymerase with a strand displacement function can circle around the template and hairpins multiple times; the consensus sequence of multiple passes yields a CCS (circular consensus sequence) read for each molecule. Raw sequences were initially processed using SMRT Link v.6.0 (Pacific Biosciences, USA). Sequences were filtered for a minimum of read length 10 bp, pass 3, and read score 65. 127,178 CCS reads were filtered through passes 3 and Q10; 89,188 CCS reads through passes 5 and Q20; 29,924 CCS reads through passes 8 and Q30. Downstream bioinformatics analysis was performed using BLASR V5.3.2 for the alignment, bcftools v.1.6 for variant calling. Mutations listed in Figure 2 and Supplementary File S2 were also manually inspected and confirmed using Integrative Genomics Viewer 2.3.32 (software.broadinstitute.org/software/igv/). Analysis steps included the following: 1. Exclude CCS reads under 1000 bases, which may have been derived from non-specific reverse transcription or PCR reactions. 2. Classify the CCS reads to the three samples, according to the PacBio barcodes on the 5’ ends. 3. For any CCS reads that contain the same 10-nucleotide random index, select only one of them, to avoid double-counting of clones derived from the same cDNA molecule. 4. Align the reads to the corresponding reference sequence (see Supplementary Files S3-S5). 5. Count the number of mutations at each nucleotide position of the reference sequences.

### Surgeries and virus injections for two-photon imaging

Adult (>9 weeks, male and female) Cre-dependent tdTomato reporter Ai14 (25) (Jackson Laboratory #007908) or Arch-EGFP reporter Ai35D (26) (Jackson Laboratory # 012735) mice were anesthetized with isoflurane (4% in oxygen) and ketamine/xylazine (100mg/kg and 10mg/kg respectively, i.p.). Mice were given buprenorphine (0.1 mg/kg s.q.) and meloxicam (2 mg/kg s.q.) as preemptive analgesics, as well as eye ointment (Puralube); the scalp was then shaved, depilated with Nair, and thoroughly rinsed before the mice were mounted on a stereotaxic instrument (Stoelting Co.) with a hand warmer (Heat Factory) underneath the animal to maintain body temperature. The scalp was then disinfected with povidone-iodine, and an incision was made at the appropriate location (see below).

A 3 mm craniotomy was opened over primary visual cortex (V1). 300 nl of LV-U-TVA950(B19G) (see above) was injected into V1 (-2.70 mm AP, 2.50 mm LM, -0.26 mm DV; AP and LM stereotaxic coordinates are with respect to bregma; DV coordinate is with respect to brain surface) using a custom injection apparatus comprised of a hydraulic manipulator (MO-10, Narishige) with headstage coupled via custom adaptors to a wire plunger advanced through pulled glass capillaries (Wiretrol II, Drummond) back-filled with mineral oil and front-filled with virus solution. Glass windows composed of a 3mm-diameter glass coverslip (Warner Instruments CS-3R) glued (Optical Adhesive 61, Norland Products) to a 5mm-diameter glass coverslip (Warner Instruments CS-5R) were then affixed over the craniotomy with Metabond (Parkell). Seven days after injection of the lentiviral vector, the coverslips were removed and 300 nl of one of the three EnvA-enveloped rabies viral vectors (with equalized titers as described above) was injected at the same stereotaxic coordinates. Coverslips were reapplied and custom stainless steel headplates (eMachineShop) were affixed to the skulls around the windows.

### *In vivo* two-photon imaging and image analysis

Beginning seven days after injection of each rabies virus and continuing every seven days up to a maximum of four weeks following rabies virus injection, the injection sites were imaged on a Prairie/Bruker Ultima IV In Vivo two-photon microscope driven by a Spectra Physics Mai-Tai Deep See laser with a mode locked Ti:sapphire laser emitting at a wavelength of 1020 nm for tdTomato and mCherry or 920 nm for EGFP. Mice were reanesthetized and mounted via their headplates to a custom frame, again with ointment applied to protect their eyes and with a handwarmer maintaining body temperature. One field of view was chosen in each mouse in the area of maximal fluorescent labelling. The imaging parameters were as follows: image size 512 X 512 pixels (282.6 μm x 282.6 μm), 0.782 Hz frame rate, dwell time 4.0 μs, 2x optical zoom, Z-stack step size 1 μm. Image acquisition was controlled with Prairie View 5.4 software. Laser power exiting the 20x water-immersion objective (Zeiss, W plan-apochromat, NA 1.0) varied between 20 and 65 mW depending on focal plane depth (Pockel cell value was automatically increased from 450 at the top section of each stack to 750 at the bottom section). For the example images of labeled cells, maximum intensity projections (stacks of 150-400 μm) were made with Fiji software. Cell counting was performed with the ImageJ Cell Counter plugin. When doing cell counting, week 1 tdTomato labelled cells were defined as a reference; remaining week 1 cells were the same cells at later time point that align with week 1 reference cells but the not-visible cells at week 1 (the dead cells). Plots of cell counts were made with Origin 7.0 software (OriginLab, Northampton, MA). For the thresholded version of this analysis (Supplementary Figure S4), in order to exclude cells that could possibly have been labeled only with mCherry in the SiR-CRE group, only cells with fluorescence intensity greater than the average of the mean red fluorescence intensities of cells imaged in Ai35 versus Ai14 mice at the same laser power at 1020 nm at 7 days postinjection (32.33 a.u.) were included in the population of cells tracked from 7 days onward.

### Monosynaptic tracing experiments: Sanger sequencing of viral genomic RNA

Sanger sequencing of RVΔG-NPEST-4Cre(EnvA) and RVΔG-N*PEST-4Cre(EnvA) was as described above but with the following modifications. The following barcoded primer was used for RT-PCR.

Adaptor-UMI-N_fp_57:

ACACTCTTTCCCTACACGACGCTCTTCCGATCTNNNNNNNNNNNNNNNNNNNNAGAAGTCCGGAGGCTG

TTTAT

cDNA sequences from the nucleoprotein gene through the first half of the rabies virus P gene were amplified using Platinum SuperFi Green Master Mix (Invitrogen (Thermo Fisher), USA) with cycling conditions as follows: denaturation at 98°C for 30 seconds, followed by 21 cycles of amplification (denaturation at 98°C for 5 seconds, annealing at 60°C for 10 seconds and extension at 72°C for 21 seconds), with a final extension at 72°C for 5 minutes, using the following primers:

i5-anchor_CTAGCGCT_fp_56: AATGATACGGCGACCACCGAGATCTACACCTAGCGCTACACTCTTTCCCTACACGAC

P_Sanger_rp_56: CAAGCAGAAGACGGCACGATTTTCCATCATCCAGGTG

The 700bp PCR amplicons for RVdG-NPEST-4Cre (EnvA) or RVΔG-N*PEST-4Cre (EnvA) virus were cloned into pEX-A (Eurofins Genomics, USA) using an In-Fusion HD Cloning Kit (Takara Bio, Japan) as above. The plasmids were transformed into Stellar competent cells (Takara Bio, Japan), and 32 clones from each rabies virus sample were isolated and purified for sequencing.

### Monosynaptic tracing experiments: surgeries and virus injections

For pilot testing of AAV1-syn-F14F15S-splitTVA-EGFP-tTA_v2 virus (Supplementary Fig. S5), 200 nl of AAV2- retro-syn-FLPo (1.16E13 g.c/mL) was injected into dorsolateral striatum (AP = +0.74 mm w.r.t. bregma, LM = 2.25 mm w.r.t. bregma, DV = -2.20 mm w.r.t the brain surface) (except when omitted for no-Flpo controls); in the same surgery, 250 nl of helper virus mixture (AAV1-syn-F14F15S-sTpEptTA_v2 (diluted to 7.88E10 g.c./mL) mixed with AAV1-TREtight-mTagBFP2-B19G (diluted to 6.50E11) in a 50/50 ratio by volume) was injected into layer 5 of barrel cortex (AP -1.55 mm w.r.t. bregma, LM 3.00 mm w.r.t. bregma, DV -0.75 mm w.r.t the brain surface) in Ai14 (het) mice. 7 days after AAV injection, 250nl of RVΔG-Cre (EnvA) (8.58E9 i.u./mL, diluted to 4.49E09 i.u./mL) was injected at the same cortical injection site as the helper viruses.

For comparisons of NPEST and N*PEST viruses, the three helper AAVs were combined at final titers of 3.6E10 genome count (“g.c.”)/mL for AAV1-syn-F14F15S-splitTVA-EGFP-tTA_v2, 6.60E11 g.c./mL for AAV1-TREtight- mTagBFP2-B19G (when included) and AAV1-TREtight-H2b-emiRFP670-P2A-TEVP (when included) in DPBS (Fisher, 14-190-250). 250 nl of helper virus mixture was injected into layer 5 of barrel cortex (AP -1.55 mm w.r.t. bregma, LM 3.00 mm w.r.t. bregma, -DV 0.75 mm w.r.t. brain surface) of Ai14 mice; in the same surgery, 300 nl of AAV2-retro-syn-FLPo (1.16E13 g.c./mL) was injected into dorsolateral striatum (AP 0.74 mm w.r.t. bregma, LM 2.25 mm w.r.t. bregma, DV -2.30 mm w.r.t the brain surface). 7 days after AAV injection, 300nl of RVΔG- NPEST-4Cre(EnvA) (1.86E9 i.u./mL) or RVΔG-N*PEST-4Cre(EnvA) (diluted 12.96-fold in DPBS from 2.41E10 i.u./mL to 1.86E9 i.u./mL) was injected in barrel cortex at the same site as the helper AAV mixtures.

### Monosynaptic tracing experiments: perfusions and histology

12 days or 3 weeks (depending on experiment; see main text) after injection of rabies virus, mice were transcardially perfused with 4% paraformaldehyde in phosphate-buffered saline. Brains were postfixed overnight in 4% paraformaldehyde in PBS on a shaker at 4°C and cut into 50 µm coronal sections on a vibrating microtome (Leica, VT-1000S). Sections were collected anterior to posteriorly into 6 tubes containing cryoprotectant. Collection goes on for 15 rounds so that each tube contains a sixth of the collected tissue (15 sections in each tube). Sections were immunostained as described(21) with a chicken anti-GFP primary antibody (Aves Labs GFP-1020) 1:500 and donkey anti-chicken Alexa Fluor 488 secondary antibody (Jackson Immuno 703-545-155) 1:200. Sections were mounted with Prolong Diamond Antifade mounting medium (Thermo Fisher P36970).

### Monosynaptic tracing experiments: cell counts and microscopy

Coronal sections between 1.2mm and -3.3mm relative to bregma were examined under an epifluorescence microscope (Zeiss, Imager.Z2). When necessary due to high density of labeled cells, images were taken with the same microscope for cell counting. tdTomato-labeled neurons in contralateral cortex and thalamus were counted manually with the Cell Counter plugin in ImageJ. For cells at the injection site, when tdTomato expressing cells were few and sparse (usually less than 100 per section), cells coexpressing mTagBFP2 or H2b- emiRFP670 alongside were counted manually adding separate labels to each and then looking for overlapping cells. When tdTomato expressing cells were dense, tdTomato labeled cells were first counted using the Analyze Particle function in ImageJ (size in micron^2: 20-400; circularity: 0.20-1.00). The outline of these cells was then merged on top of images of mTagBFP2 and H2b-emiRFP670 labeled cells for the counting of the overlapping cells. Only one of the six series of sections (i.e., every sixth section: see above) was counted for each mouse.

Images for figures were taken on a confocal microscope (Zeiss, LSM 900). So that the confocal images of brain tissue included in the figures in this paper are representative of each group, the images were taken after the counts were conducted (see above), in each case using the mouse with the middle number of labeled neurons in that group (i.e., neither the highest nor the lowest in the group of 3 mice used for each condition).

**Supplementary Video S1 *(separate file)*: Video of 95% of SiR-CRE-labeled neurons in an Arch-EGFP-ER2 reporter mouse disappearing between 11 days and 28 days postinjection.** Two-photon image stacks of a single FOV of visual cortical neurons in an Ai35 mouse imaged at four different time points; time in the video represents depth of focus. Large blobs are glia. 18 out of the 19 neurons visibly labeled with Arch-EGFP-ER2 at 11 days following injection of SiR-Cre are no longer visible 17 days later. White circles indicate cells present at both 11 days and all subsequent imaging sessions; red circles indicate cells present at 11 days but gone by 28 days.

---

**Supplementary Figure S1:**
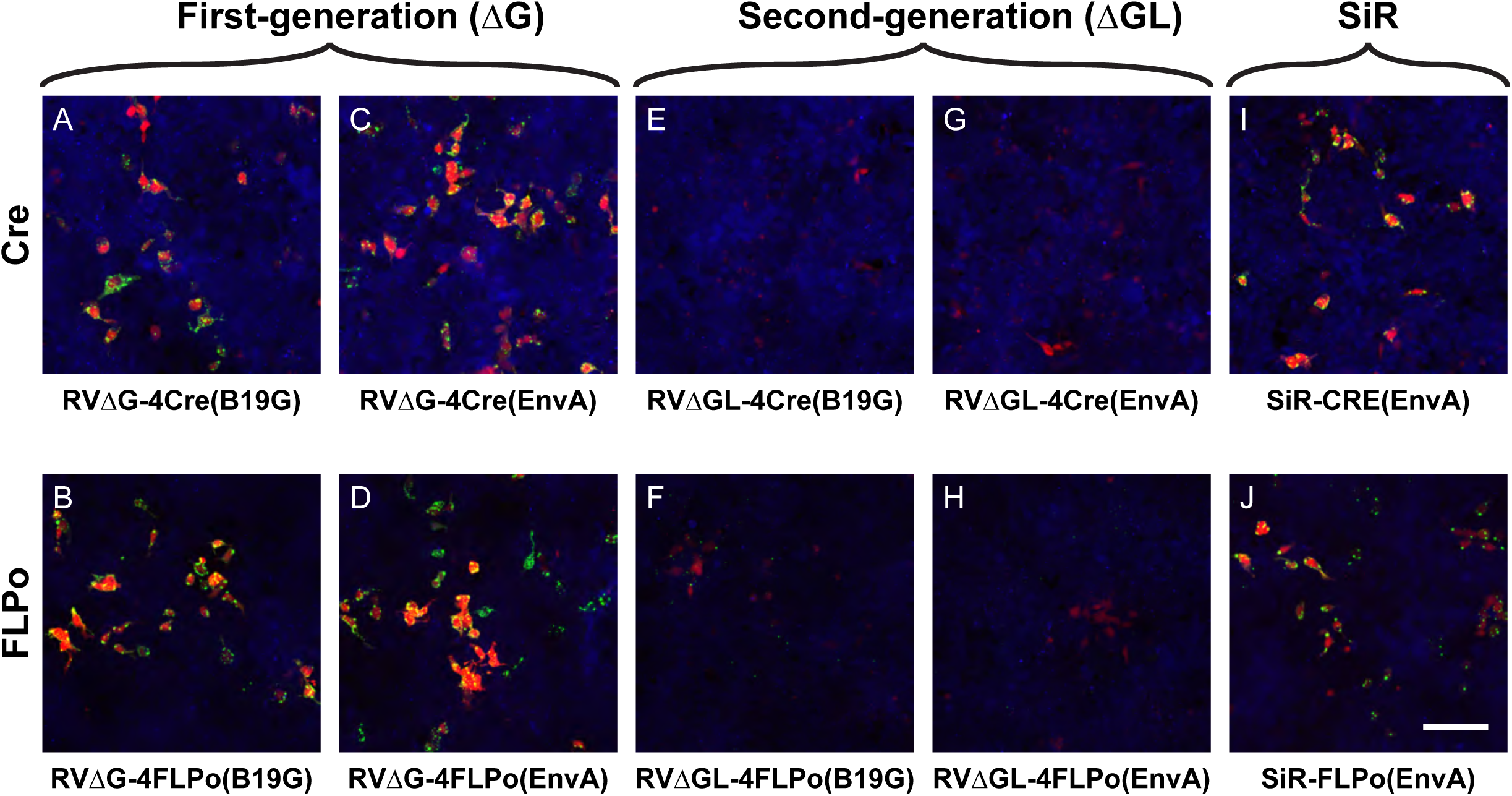
SiR viruses appeared to cause expression of viral nucleoprotein at levels similar to those of first-generation ΔG viruses. (A-D) Reporter cells infected with first-generation, ΔG viruses show characteristic bright, clumpy anti-nucleoprotein staining (green), indicating high nucleoprotein expression and active viral replication. Red is mCherry expression, reporting expression of Cre or FLPo; blue is mTagBFP2, constitutively expressed by these reporter cell lines. (E-H) Reporter cells infected with second-generation, ΔGL viruses show only punctate staining for nucleoprotein, indicating isolated individual viral particles or ribonucleoprotein complexes; these viruses do not replicate intracellularly (21). Reporter cassette activation takes longer from the lower recombinase expression levels of these viruses, so mCherry expression is dimmer than in cells infected with ΔG viruses at the same time point. (I-J) Reporter cells infected with SiR viruses show clumps of nucleoprotein and rapid reporter expression indicating high expression of recombinases, similarly to cells infected with ΔG viruses. Scale bar: 100 µm, applies to all panels.

**Supplementary Figure S2:**
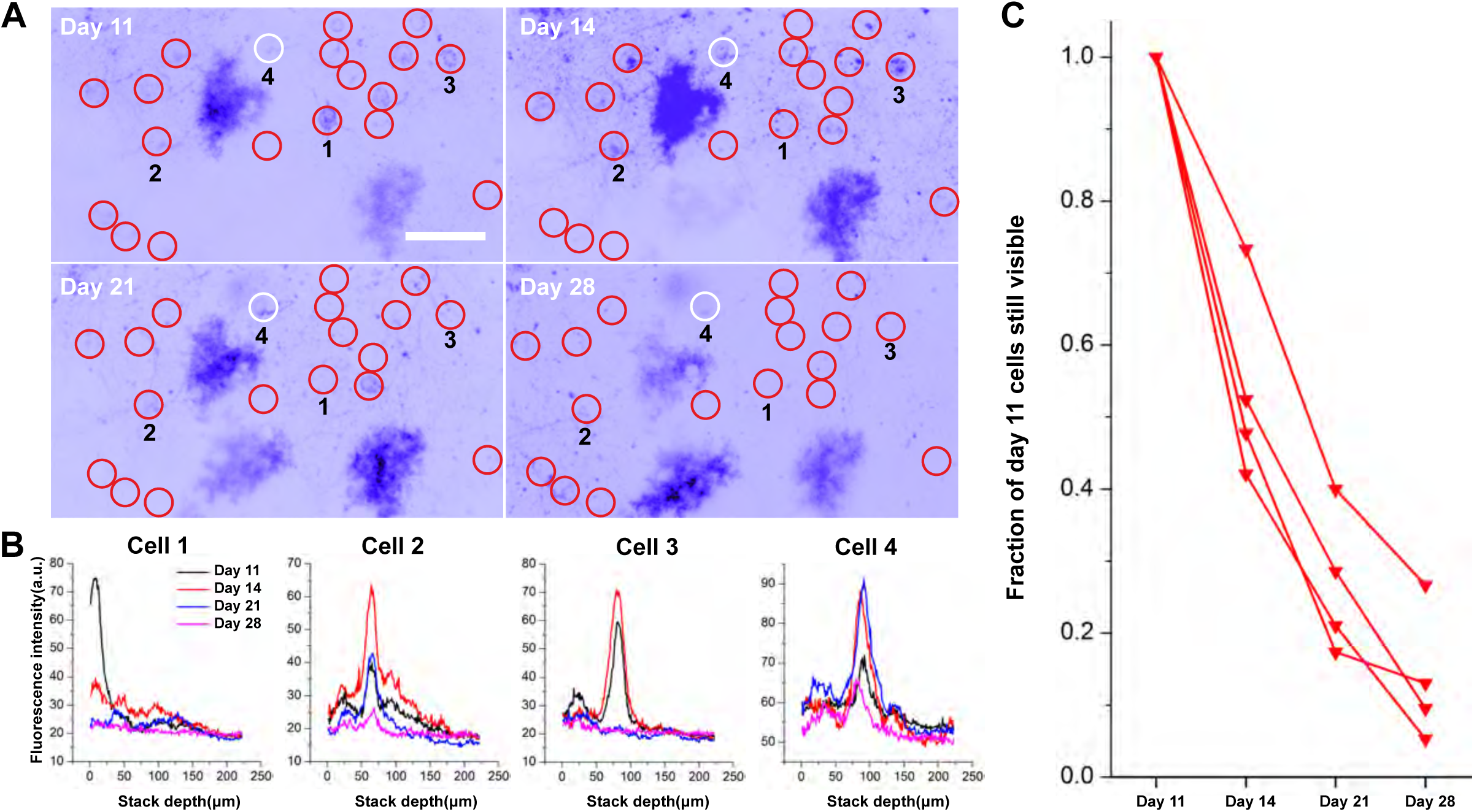
86% of SiR-CRE-labeled neurons in Arch-EGFP-ER2 reporter mice disappeared between 11 days and 28 days after injection. A) Maximum intensity projections of the two-photon FOV shown in Supplementary Video S1 of visual cortical neurons labeled with SiR-CRE in an Ai35 mouse, 11-28 days postinjection. Images are from the same FOV at four different time points. All cells clearly visible on day 11 are circled. In this example, 18 out of 19 cells (red circles) disappeared by a subsequent imaging session. Only one cell (white circle) is still visible on day 28. Numbers below four of the circles mark the cells for which intensity profiles are shown in panel B. Scale bar: 50 µm, applies to all images. See Supplementary Video S1 for a clearer view of the same cells. B) Green fluorescence intensity versus depth for the four representative neurons numbered in panel A at the four different time points, showing disappearance of three of them over time. C) Fraction of cells visibly EGFP-labeled at day 11 still visible at later time points, from four different FOVs in two Ai35 mice. Connected sets of markers indicate cells from the same FOV. 86% of SiR-CRE-labeled neurons had disappeared by 4 weeks postinjection.

**Supplementary Figure S3:**
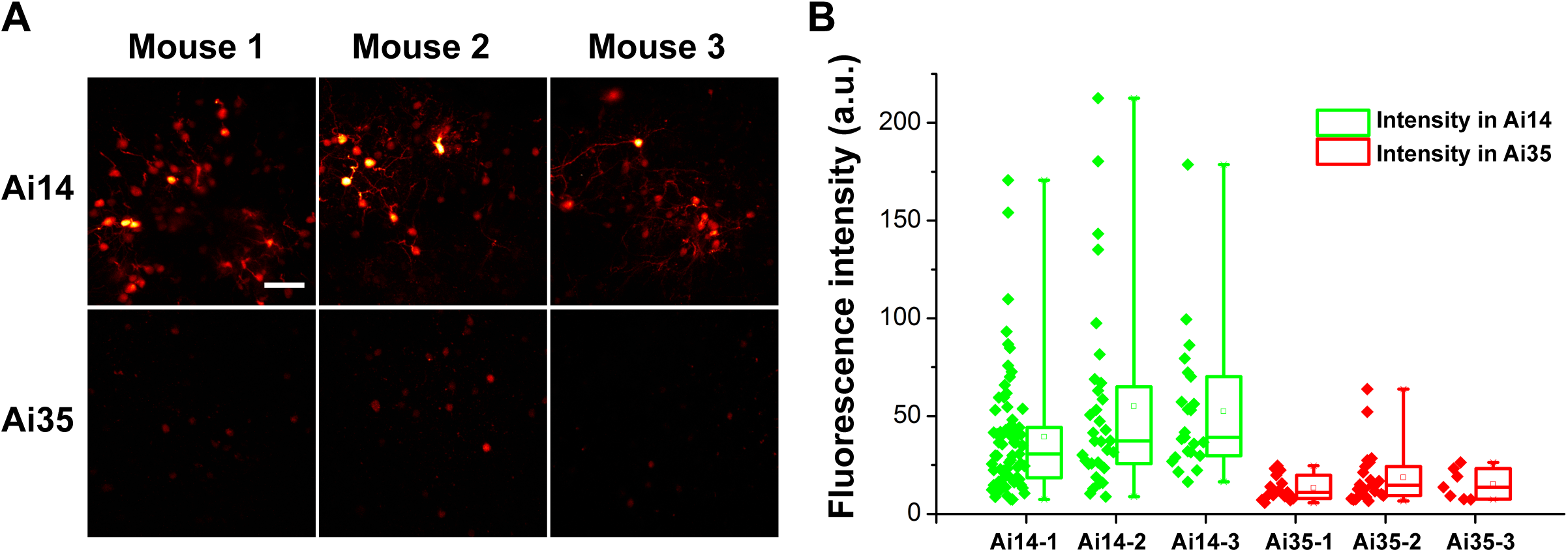
mCherry fluorescence from SiR-CRE is much dimmer than tdTomato fluorescence in Ai14 mice, suggesting that the disappearance of the brighter cells in SiR-CRE-injected Ai14 mice indicates their death. A) Representative images of red fluorescence in SiR-CRE-labeled cells in Ai14 (Cre-dependent expression of tdTomato, top row) and Ai35 (Cre-dependent expression of Arch-EGFP-ER2, bottom row). The three images for each mouse line are from 3 different mice of each line, imaged 7 days following SiR-CRE injection (see Methods), all with the same laser intensity and wavelength (1020 nm). Red fluorescence due only to mCherry (i.e., in Ai35 mice) is obviously much dimmer than that due to tdTomato (i.e., in Ai14 mice). Scale bar: 50 µm, applies to all images. B) Intensity of red fluorescence of SiR-CRE-labeled cells in Ai14 (left) and Ai35 (right) mice. Data point indicate intensity of individual cells in arbitrary units at the same laser and microscope settings (see Methods). Box plots indicate median, 25th–75th percentiles (boxes), and full range (whiskers) of intensities for each mouse. The average of the mean red fluorescent intensity in each mouse was 48.97 in Ai14 and 15.69 in Ai35 (p=0.00283 < 0.01, one-way ANOVA); the midpoint of these means, 32.33, was used as the cutoff for the reanalysis of the data in Ai14 mice to exclude neurons that could have been labeled with mCherry alone.

**Supplementary Figure S4:**
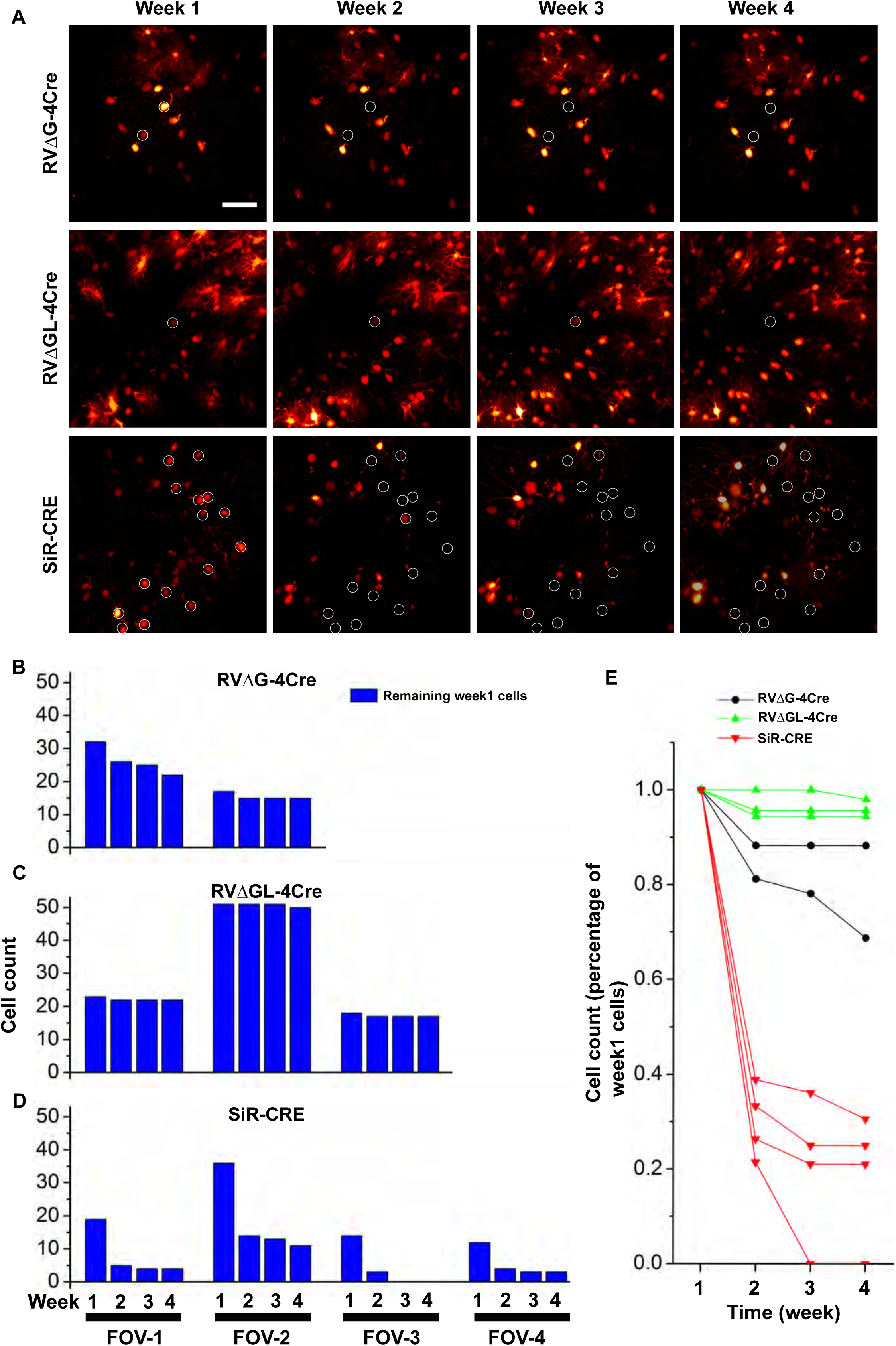
81% of SiR-CRE-labeled neurons in tdTomato reporter mice disappear within 2-4 weeks, even excluding dimmer cells that might have only been labeled with mCherry. A) Same representative fields of view as in Figure 3 but with circles now marking only cells with intensity at 7 days of greater than 32.33 a.u. (see text and Supplementary Figure S3) that are no longer visible at a subsequent time point. Scale bar: 50 µm, applies to all images. B-D) Numbers of cells above threshold fluorescence intensity at week 1 that were still present in subsequent weeks. The conclusions from Figure 3 are unchanged: few cells labeled with RVΔGL- 4Cre were lost, RVΔG-4Cre killed a significant minority of cells, and SiR-CRE killed the majority of labeled neurons within two weeks following injection. F) Percentages of cells above threshold at week 1 that were still present in subsequent imaging sessions. By 28 days postinjection, an average of only 19.2% of suprathreshold SiR-CRE-labeled cells remained.

**Supplementary Figure S5:**
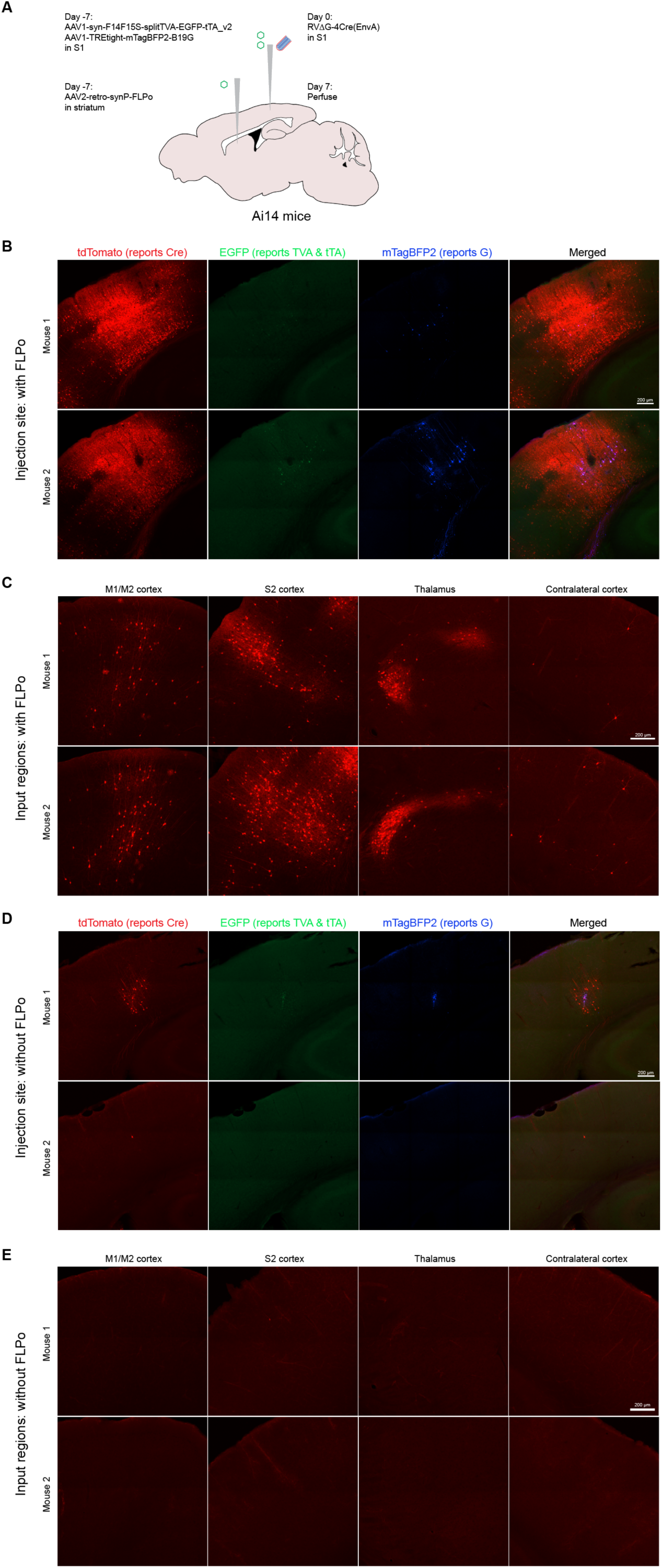
Pilot testing of Flp-dependent helper virus AAV1-syn-F14F15S- splitTVA-EGFP-tTA_v2. A) Schematic of experimental design to test new Flp-dependent helper virus AAV1-syn-F14F15S- splitTVA-EGFP-tTA_v2, which was designed without the error discovered in AAV1-syn-F14F15S- splitTVA-EGFP-tTA, used in the initial preprint of this paper. The new helper virus as well as AAV1-TREtight-BFP2-B19G were injected in S1, with AAV2-retro-syn-Flpo injected in striatum (or not, for no-Flpo controls), of two Ai14 mice, in order to target corticostriatal neurons. The first- generation rabies virus RVΔG-Cre(EnvA) was injected one week after helper virus injection, and mice were perfused one week later. B-C) Representative confocal images of injection site (B) and input regions (C) with FLPo virus in each of the two mice. D-E) Representative confocal images of injection site (D) and input regions (E) without FLPo virus in each of the two mice. Far fewer cells are labeled at the injection sites; very cells are present in input regions.

**Supplementary Figure S6:**
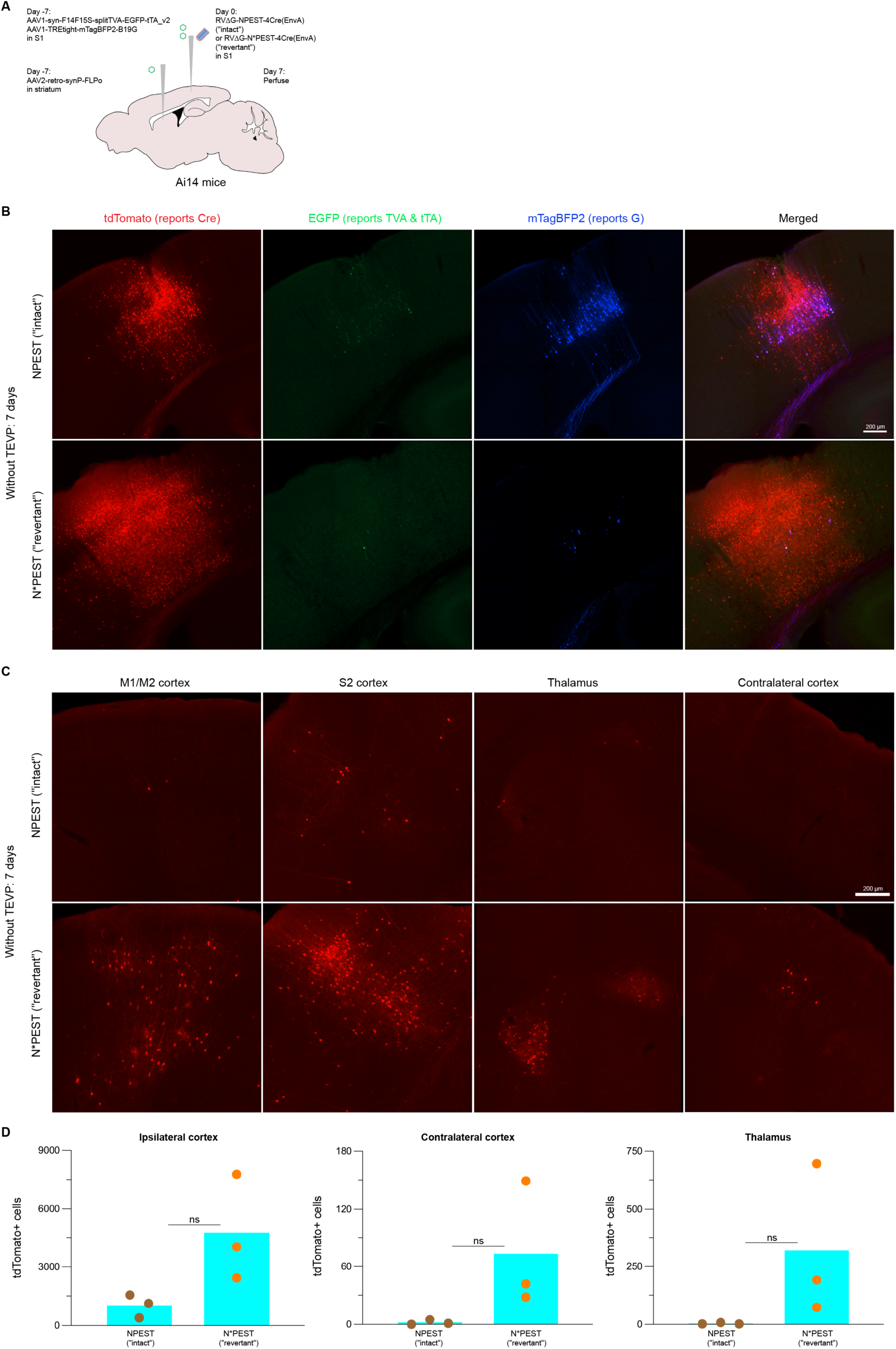
*In vivo* comparison of NPEST and N*PEST viruses without TEVP: results at 7 days (cf. Figure 4). A) Schematic of experiment. B-D) Results at 7 days after injection are similar to those at 21 days (see Fig. 4) but not statistically significant due to high variance. B) Representative images of injection sites and surrounding ipsilateral cortex. C) Representative images of input regions. D) Counts of labeled neurons in ipsilateral cortex (left), contralateral cortex (center), and thalamus (right) at 7 days for the two viruses. Each dot indicates the total number of labeled cells found in a given mouse brain when examining every sixth 50 µm section (see Methods). Although the numbers of labeled cells are much higher in every case for the N*PEST virus than for the NPEST one, the comparisons are not statistically significant (see Supplementary File S8 for all counts and statistical analyses).

**Supplementary Figure S7:**
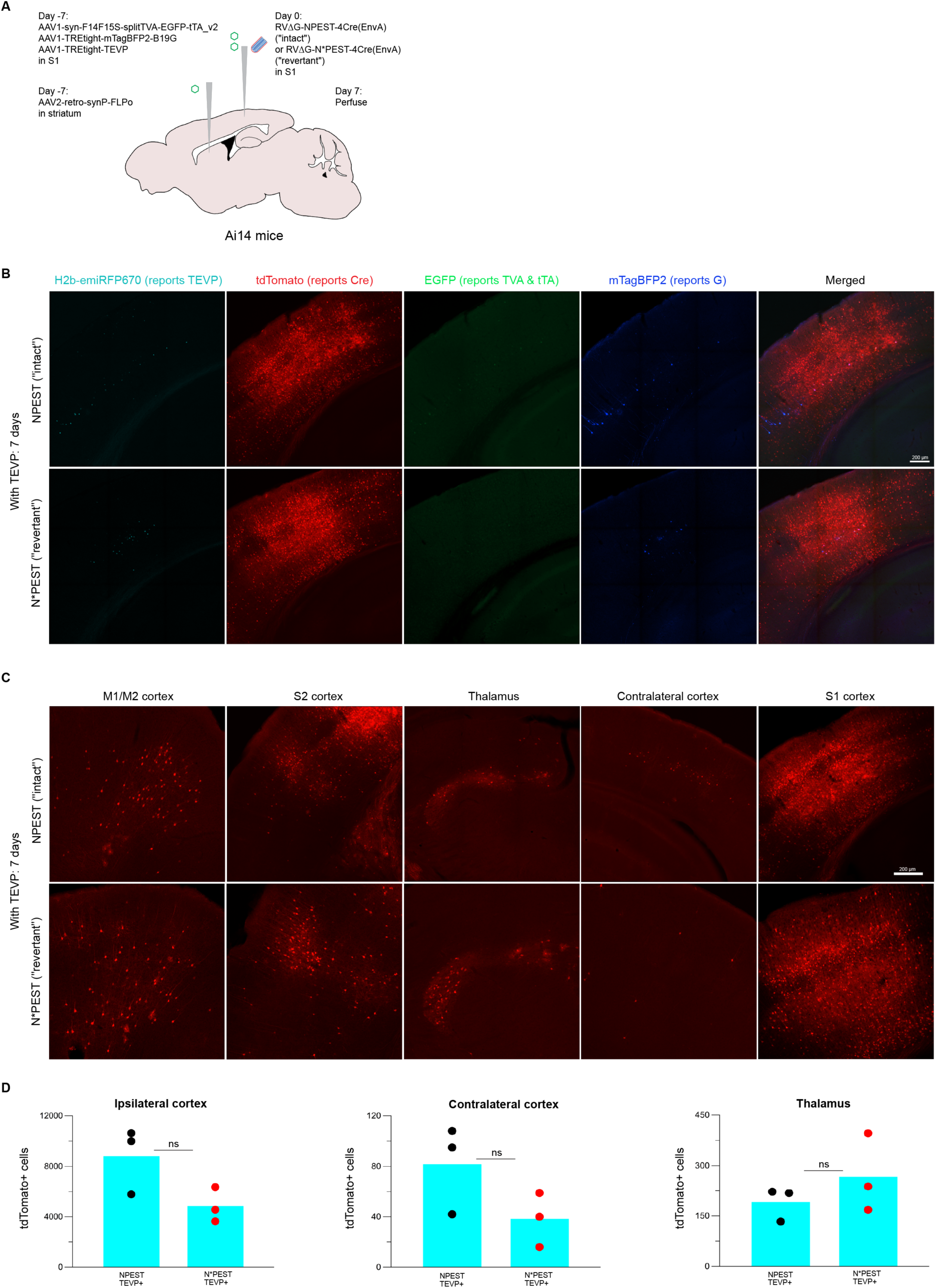
*In vivo* comparison of NPEST and N*PEST viruses with TEVP: results at 7 days (cf. Figure 5). A) Schematic of experiment. B-D) Results at 7 days after injection are similar to those at 21 days (see Fig. 5). B) Representative images of injection sites and surrounding ipsilateral cortex. C) Representative images of input regions. D) Counts of labeled neurons in ipsilateral cortex (left), contralateral cortex (center), and thalamus (right) at 7 days for the two viruses. Each dot indicates the total number of labeled cells found in a given mouse brain when examining every sixth 50 µm section (see Methods). At this timepoint, the average numbers of labeled neurons in ipsilateral and contralateral cortex are higher for the NPEST virus, whereas the average number of labeled thalamic neurons is higher for the N*PEST virus, although again the comparisons are not statistically significant (see Supplementary File S8 for all counts and statistical analyses).

**Supplementary Figure S8:**
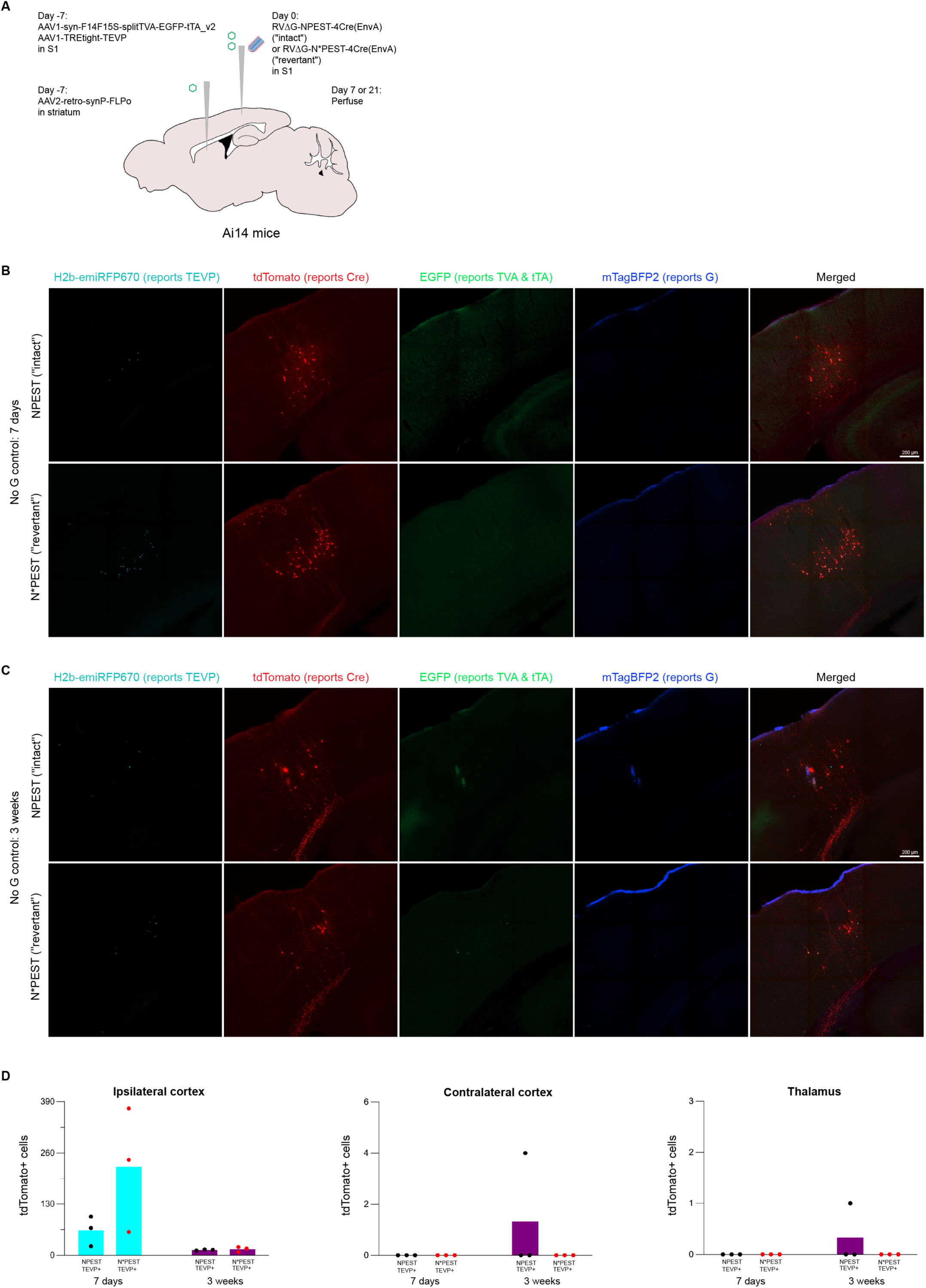
Results for control experiments without G. A) Schematic of experiment. Design is the same as for the transsynaptic tracing experiments with the helper virus expressing TEVP included, except that in these experiments the G-encoding helper virus, AAV1-TREtight-BFP2-B19G, was omitted. B-C) Images of injection sites using NPEST (top) and N*PEST (bottom) viruses, with survival times of 7 days (B) or 3 weeks (C). Scale bar: 200 µm, applies to all images in each panel. D) Counts of tdTomato-labeled neurons in ipsilateral cortex, contralateral cortex, and thalamus. Few cells were found in input regions with either virus at either survival time, suggesting that the extensive label found there in experiments in which G was included was primarily due to *bona fide* transneuronal spread rather than direct retrograde infection by injected rabies virus..

**Supplementary Figure S9:**
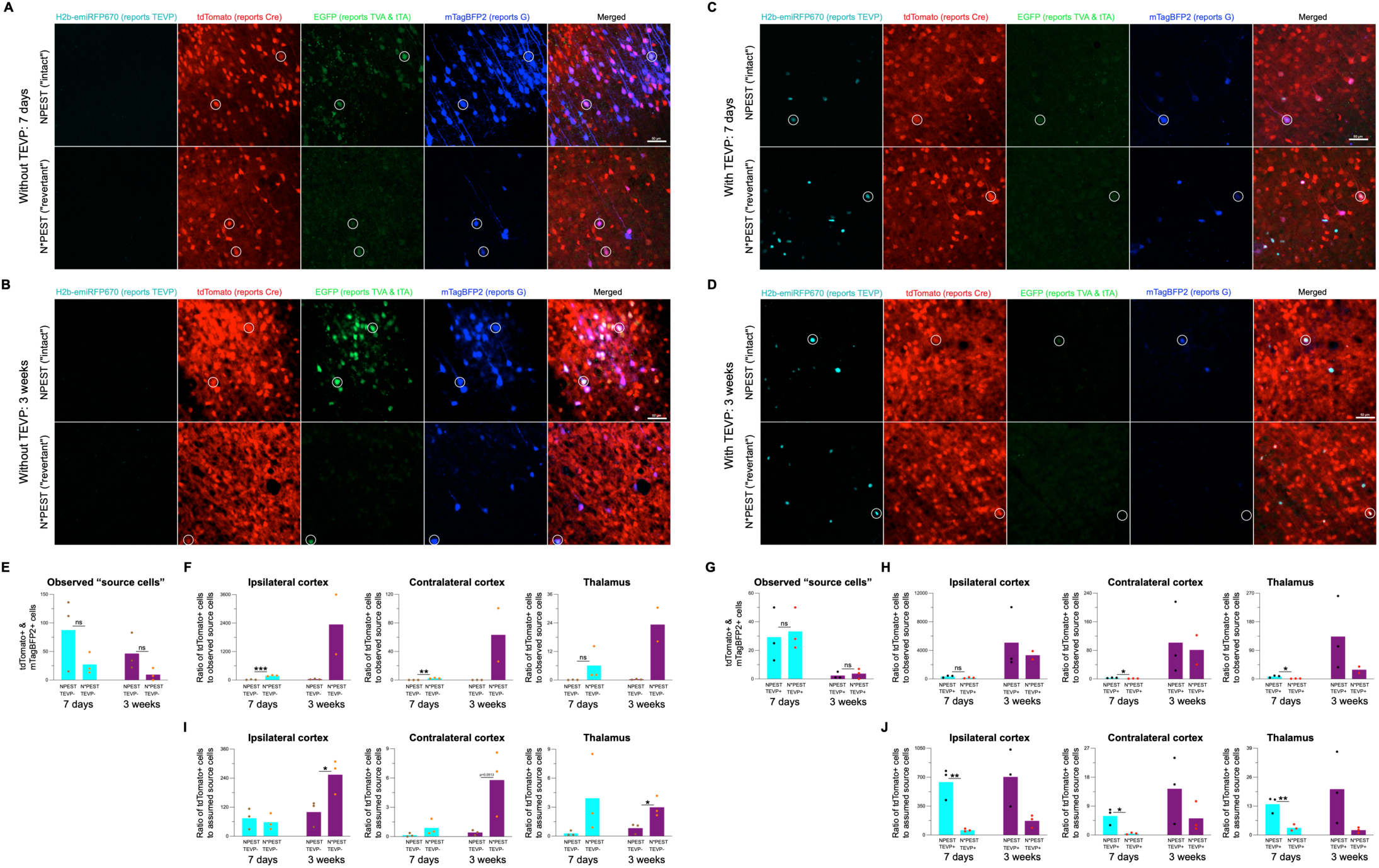
Images and counts of source cells for all conditions. A-D) Representative images of injection sites without TEVP (A-B) or with TEVP (C-D), with 7-day (A & C) or 3-week (B & D) survival time following rabies virus injection. Top rows: NPEST (“intact”); bottom rows: N*PEST (“revertant”). Scale bars: 50 µm, apply to all images in each panel. E-H) Counts of “source cells” and ratios of input cells to source cells in the absence (E-F) or presence (G-H) of TEVP. Source cells are here defined as cells coexpressing tdTomato and mTagBFP2 when TEVP was omitted, and cells coexpressing tdTomato, mTagBFP2, and emiRFP670 when TEVP was supplied. Note that the average numbers of source cells at 3 weeks are in most cases much lower than at 7 days, for both viruses, with or without TEVP, possibly suggesting toxicity to source cells that results in the death of the majority. Because of this apparent toxicity to source cells, the ratios of input cells to source cells may be artificially high and therefore arguably not particularly meaningful, particularly at the later timepoint: for example, for N*PEST at 3 weeks both with and without TEVP, no source cells were found (E & G), so the ratios for that mouse would be infinite and are omitted from the graphs (F & H). We have therefore omitted significance comparisons for the 3-week timepoint from the graphs, although Supplementary File S8 contains all counts and statistical analyses. I-J) Same as graphs in panels F & G but using the mean number of tdTomato+/emiRFP670+ cells in the 7d G-/TEVP+ control condition for each virus (i.e., 225 for N*PEST, 63.3 for NPEST; cf. Supplementary Figure S8 and Supplementary File S8) as proxy for the unknown number of original source cells in each case.

**Supplementary Figure S10:**
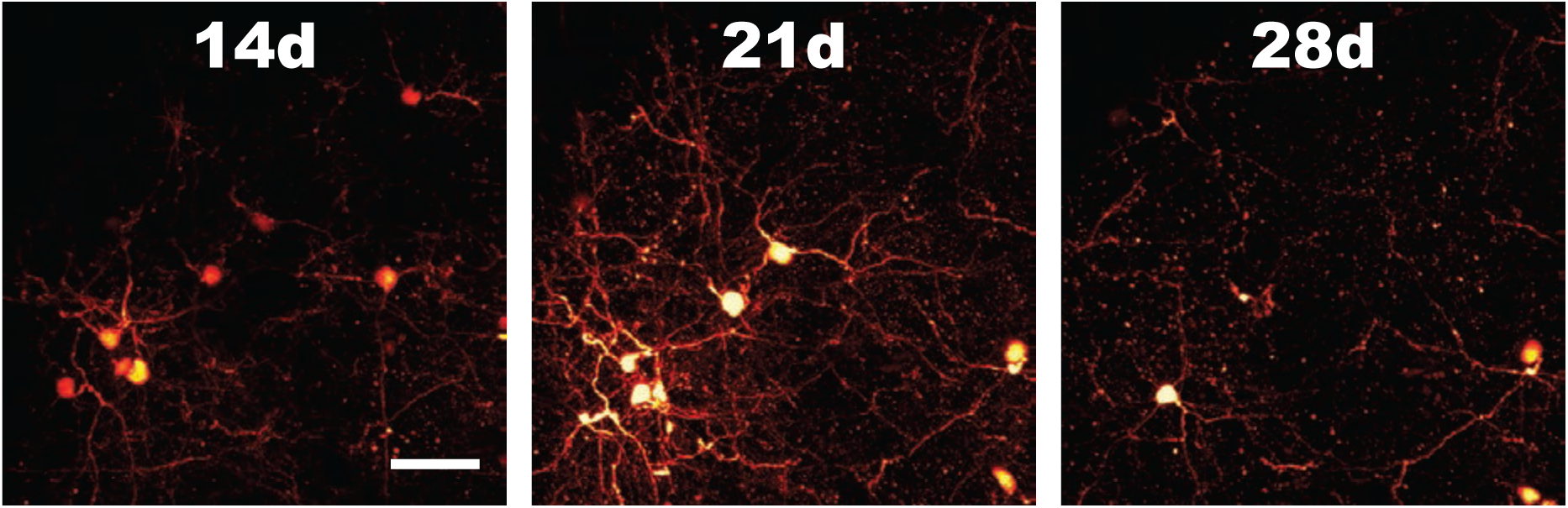
First-generation vector RVΔG-4FLPo appears to be toxic to most cells, unlike the comparable first-generation vector RVΔG-Cre. Although we did not rigorously quantify the effect, our FLPo-encoding RVΔG-4Flpo appears to kill neurons more quickly than does RVΔG-4Cre (cf. Figure 3 and Chatterjee et al. (21)). In this example field of view, most neurons clearly visible at earlier time points have disappeared by 28 days postinjection, leaving degenerating cellular debris. See Discussion for possible reasons why a preparation of a first-generation vector encoding a recombinase may or may not preserve a large percentage of infected neurons. Scale bar: 50 µm, applies to all panels.

**Supplementary File S1:**
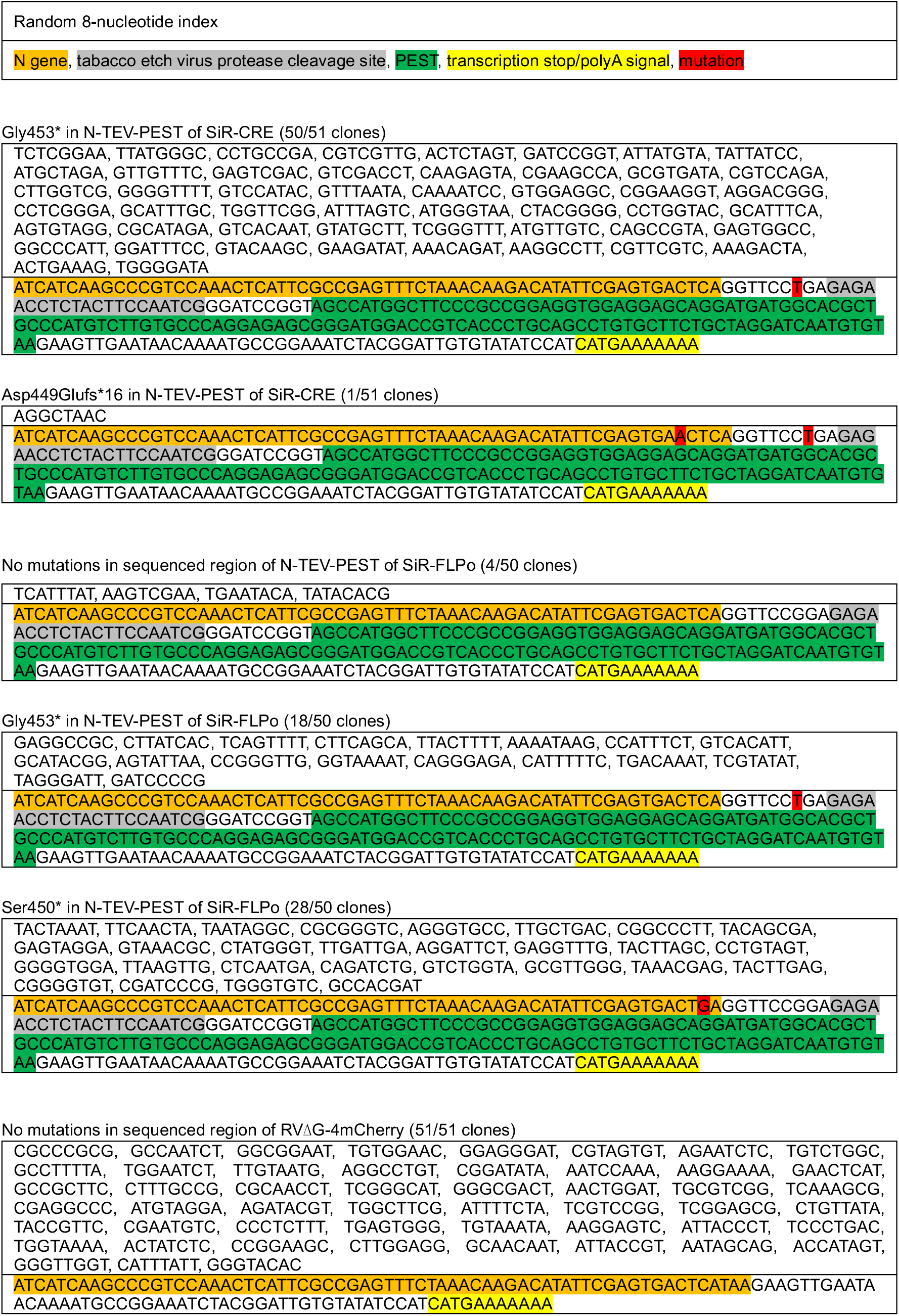
Sanger sequencing data of all clones shown in Figure 1. 51 clones derived from SiR-CRE, 50 from SiR-FLPo, and 51 from RVΔG-4mCherry are identified by their unique indices. All of the indices as well as the sequences corresponding to the 3’ end of the nucleoprotein gene are shown.

**Supplementary File S2: Summary tables of SMRT sequencing data.**

These tables show all mutations occurring at positions mutated at greater than 2% frequency in the three virus samples analyzed. Position numbers in these tables refer to the sequences in the three Genbank files below (Supplementary Files S3-S5).

**SINGLE-MOLECULE, REAL-TIME SEQUENCING RESULTS**

Frameshifts and insertions

Position numbers in this file refer to the reference sequences included as Supplementary Files S4-S5. A “frameshift” is included in Tables 1a-2b if the number of deleted bases in positions 1439-1492 (the vicinity of the junction of the end of the N gene and the intended 3’ addition) is not an integer multiple of 3, with insertions ignored. “Any error” includes either the apparent frameshifts, or the new TAA/TAG/TGA stop codons, or both, with insertions ignored. The number of “frameshifts” increases considerably if insertion mutations are included in the calculation, indicating that there is a much higher insertion rate as compared to that of deletion; however, previous studies have found that spurious insertions are high with SMRT (see main text), so we ignore insertions in this paper apart from summarizing the data below.

**Table 1a.**
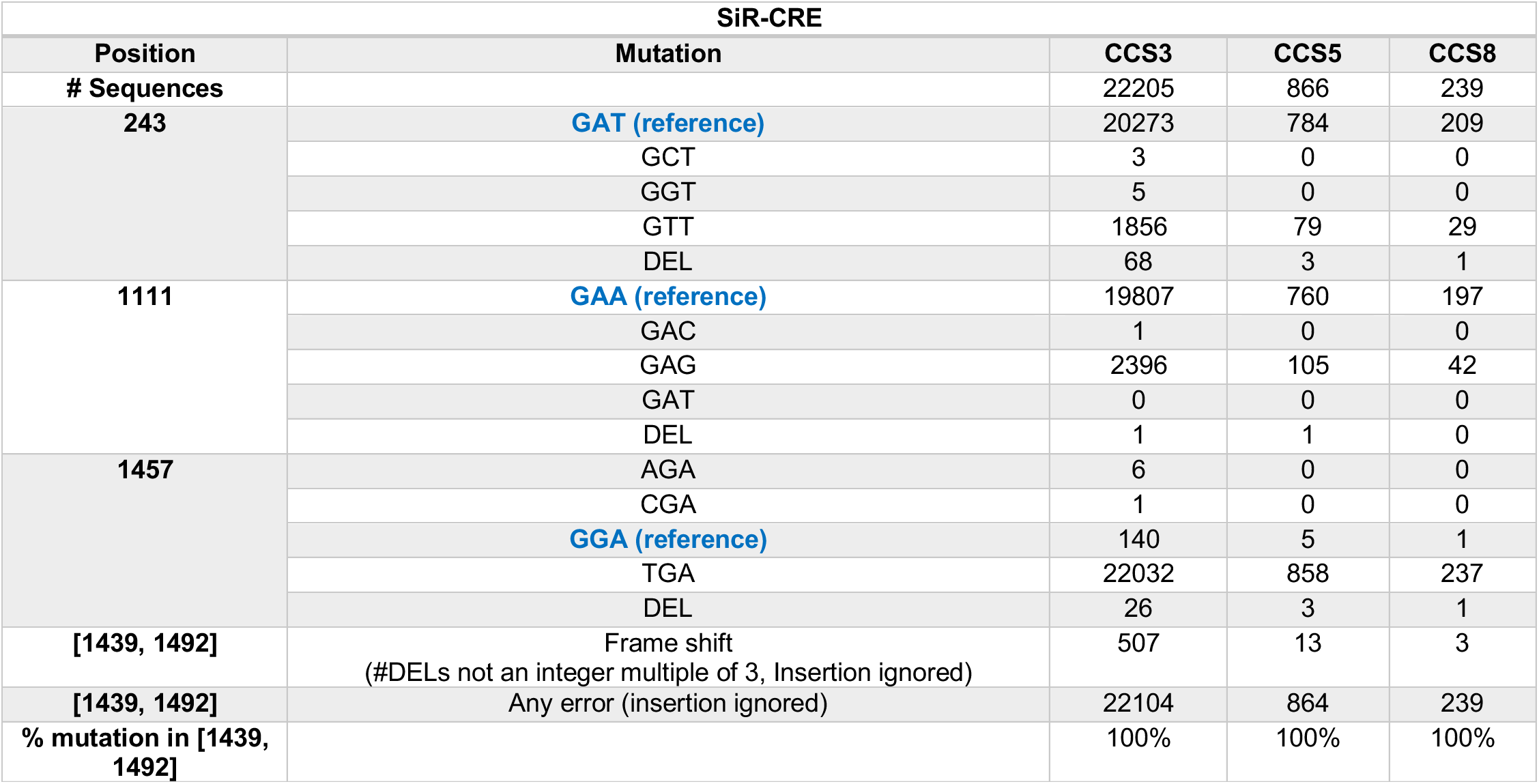
All mutations in the SiR-CRE sample at positions mutated at >2% frequency at all stringencies (CCS3, CCS5, CCS8), as well as frameshift mutations found in the C-terminal region of N.

**Table 1b.**
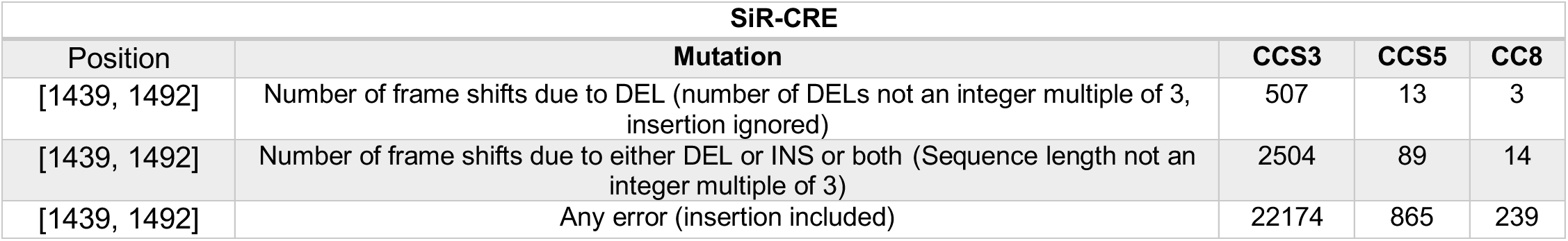
Frameshift mutations in the C-terminal region of N in the SiR-CRE sample at positions mutated at >2% frequency at all stringencies (CCS3, CCS5, CCS8).

**Table 2a.**
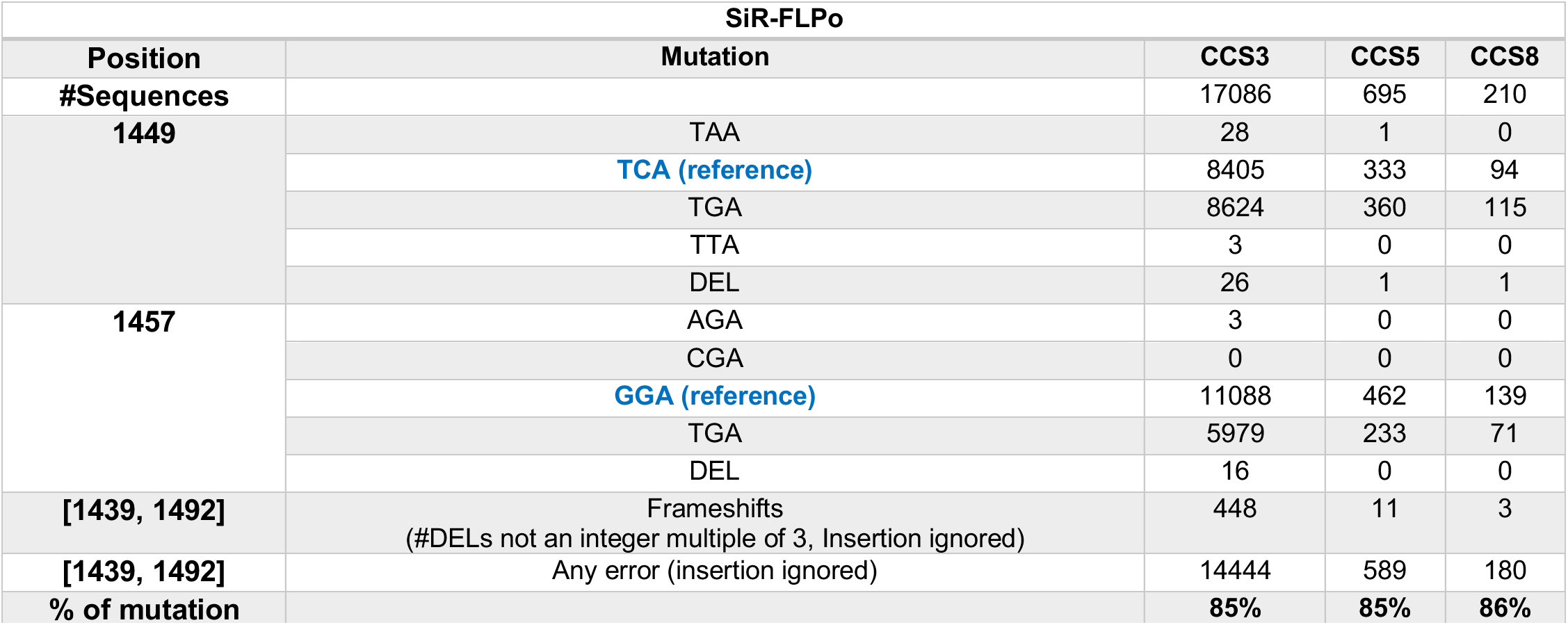
All mutations in the SiR-FLPo sample at positions mutated at >2% frequency at all stringencies (CCS3, CCS5, CCS8), as well as frameshift mutations found in the C-terminal region of N.

**Table 2b.**
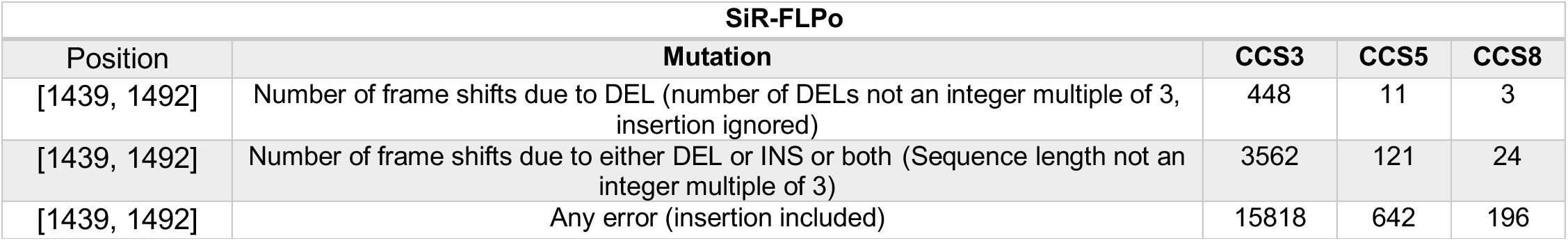
Frameshift mutations in the C-terminal region of N in the SiR-FLPo sample at positions mutated at >2% frequency at all stringencies (CCS3, CCS5, CCS8).

**Table 3.**
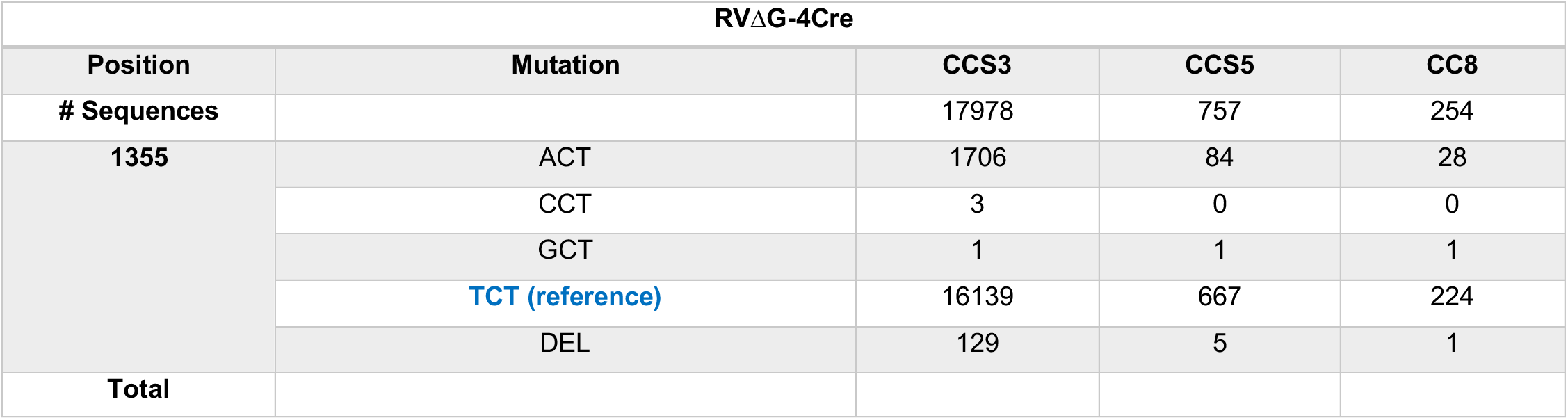
All mutations in the SiR-FLPo sample at positions mutated at >2% frequency at all stringencies (CCS3, CCS5, CCS8).

Tables of all mutations above 2% frequency threshold

Table 4a to 4c list all single-nucleotide substitutions and deletions at positions mutated at >2% threshold frequency. The percentage of mutations is calculated based on the total number of single nucleotide and deletion mutations divided by the total number of reads aligned, when insertion mutations are ignored. Deletion mutations dominate in the medium-frequency range between 2% and 5%.

**Table 4a.**
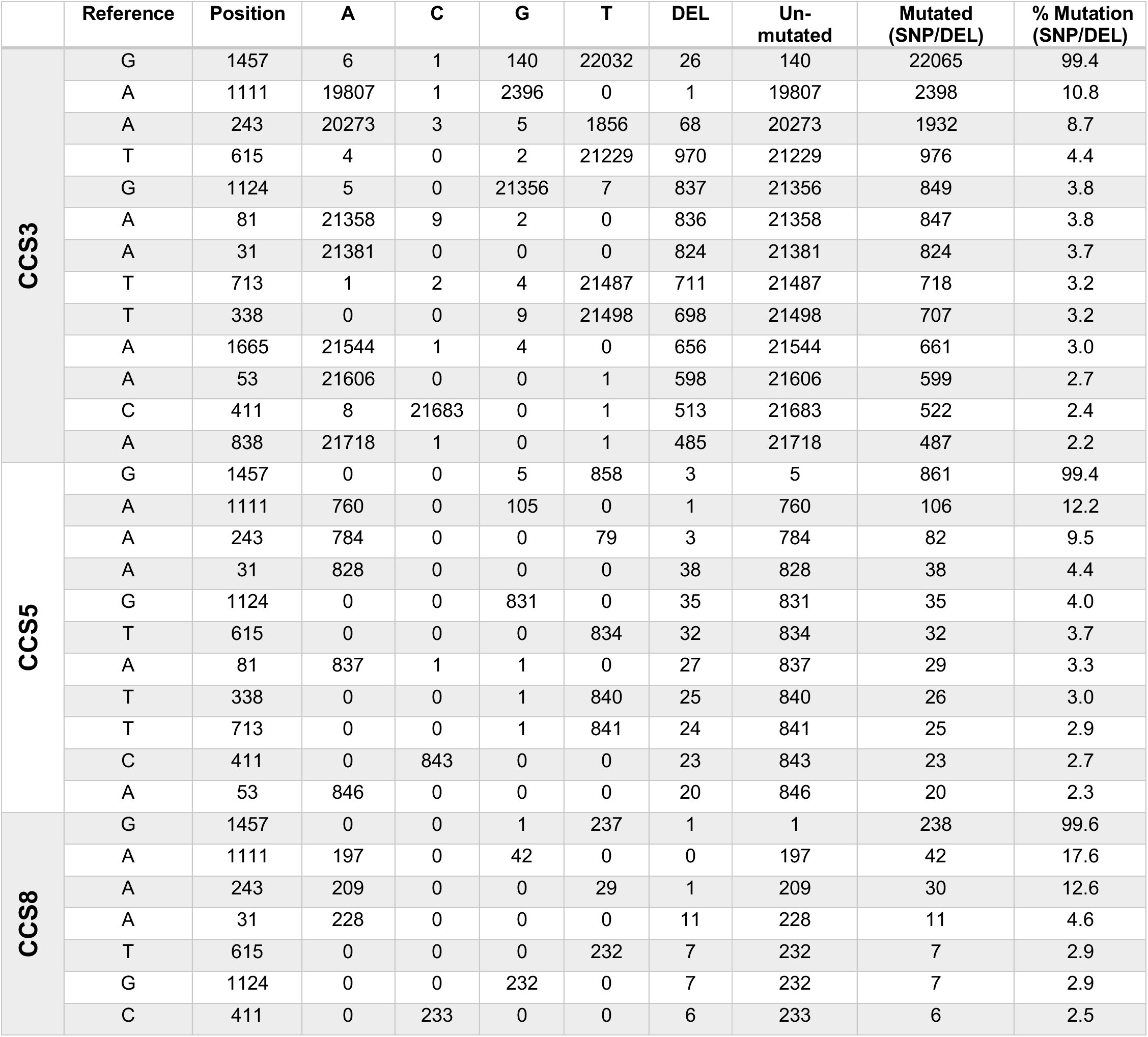
SiR-CRE: substitutions and deletions at positions mutated at >2% frequency at all stringencies (CCS3, CCS5, CCS8).

**Table 4b.**
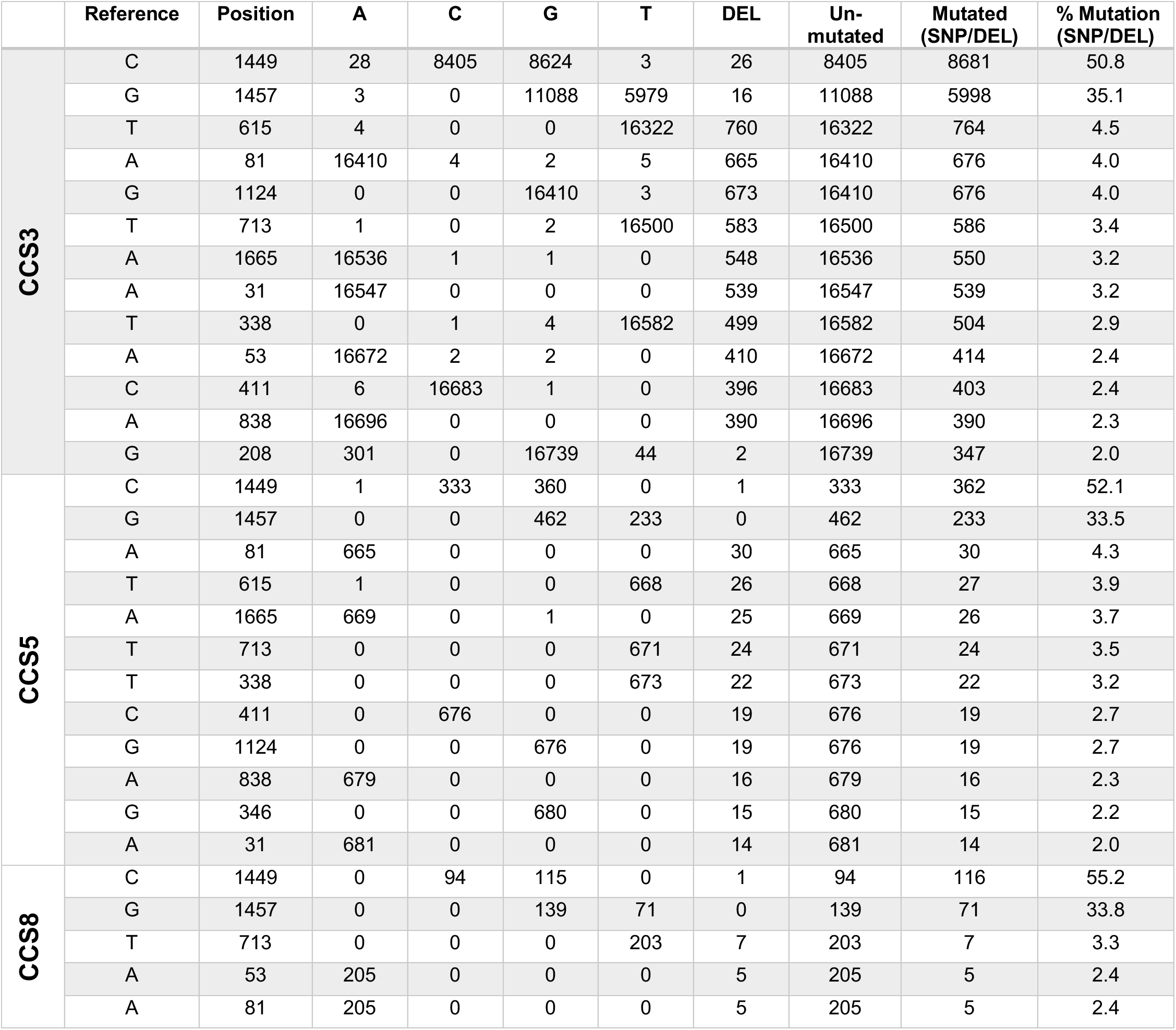
SiR-FLPo: substitutions and deletions at positions mutated at >2% frequency at all stringencies (CCS3, CCS5, CCS8).

**Table 4c.**
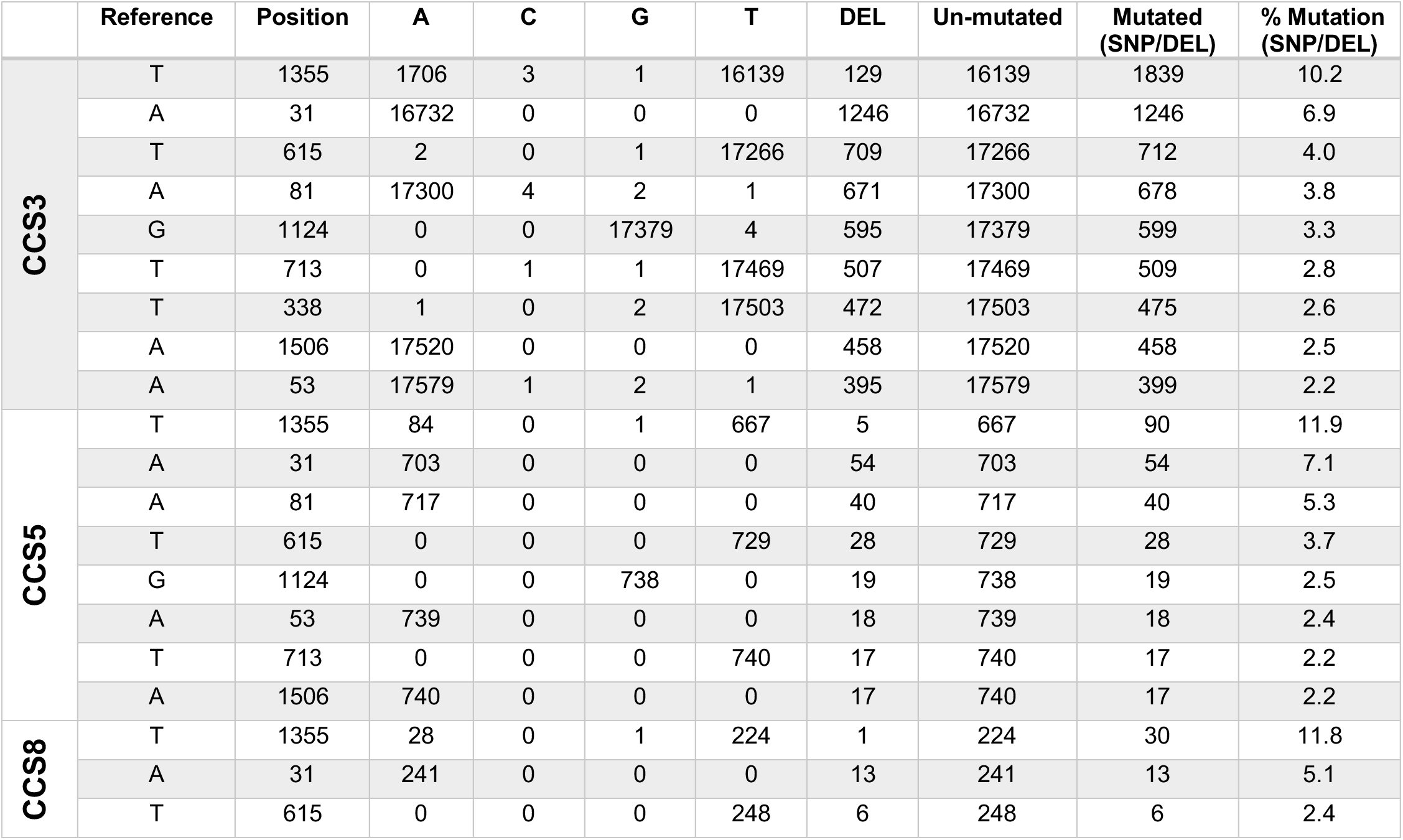
RVΔG-4Cre: substitutions and deletions at positions mutated at >2% frequency at all stringencies (CCS3, CCS5, CCS8).

**Supplementary File S3:**
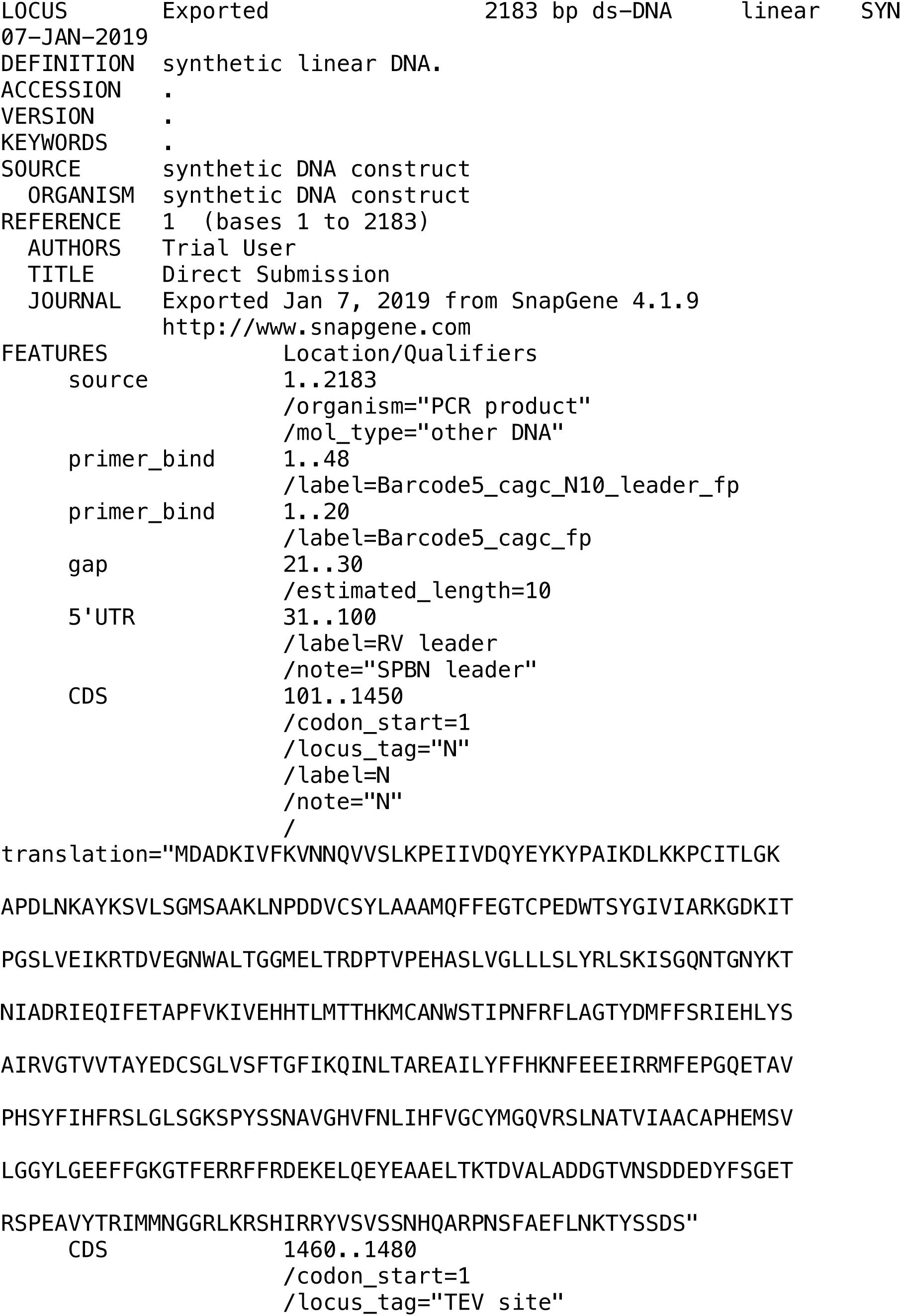
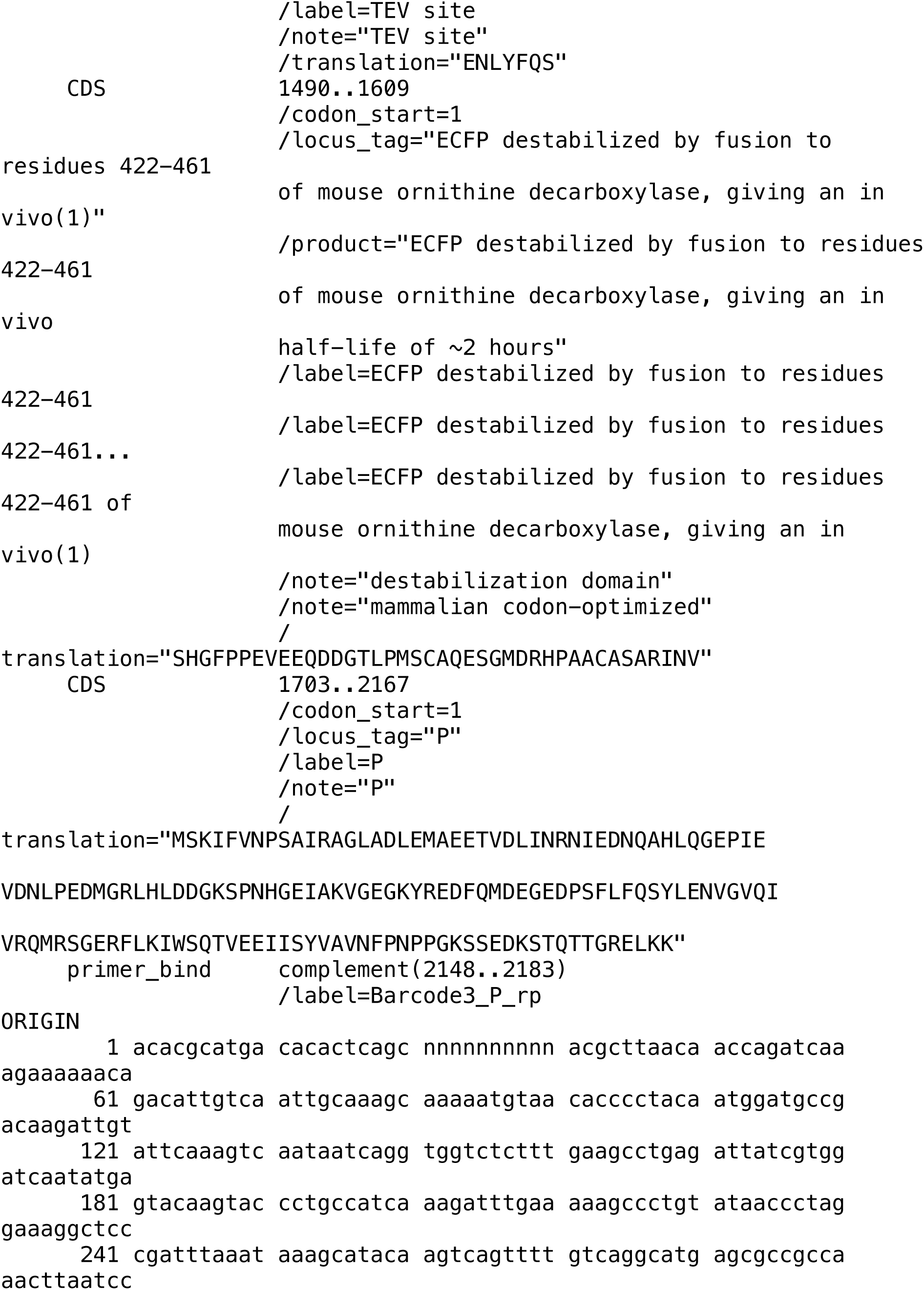
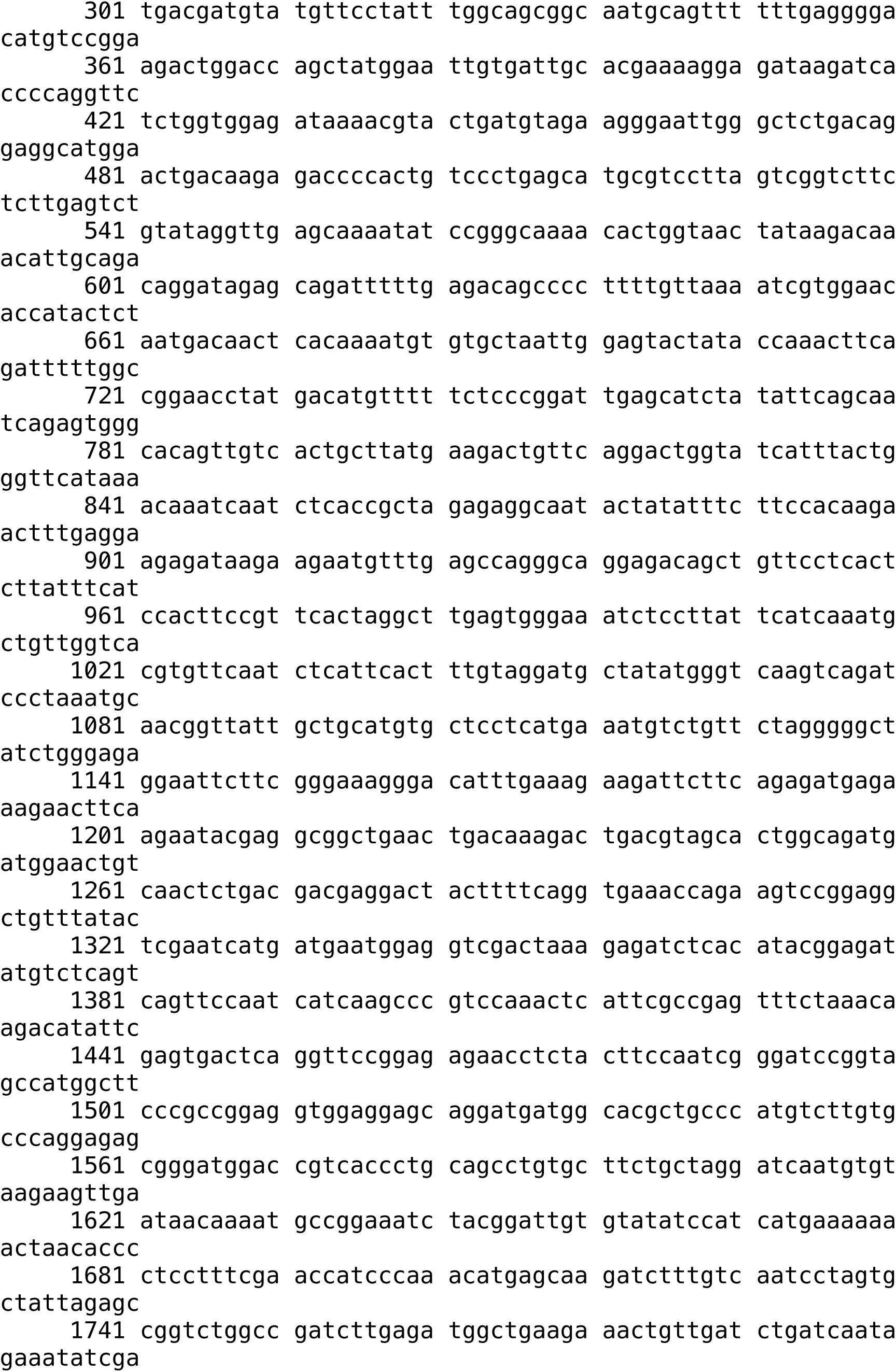
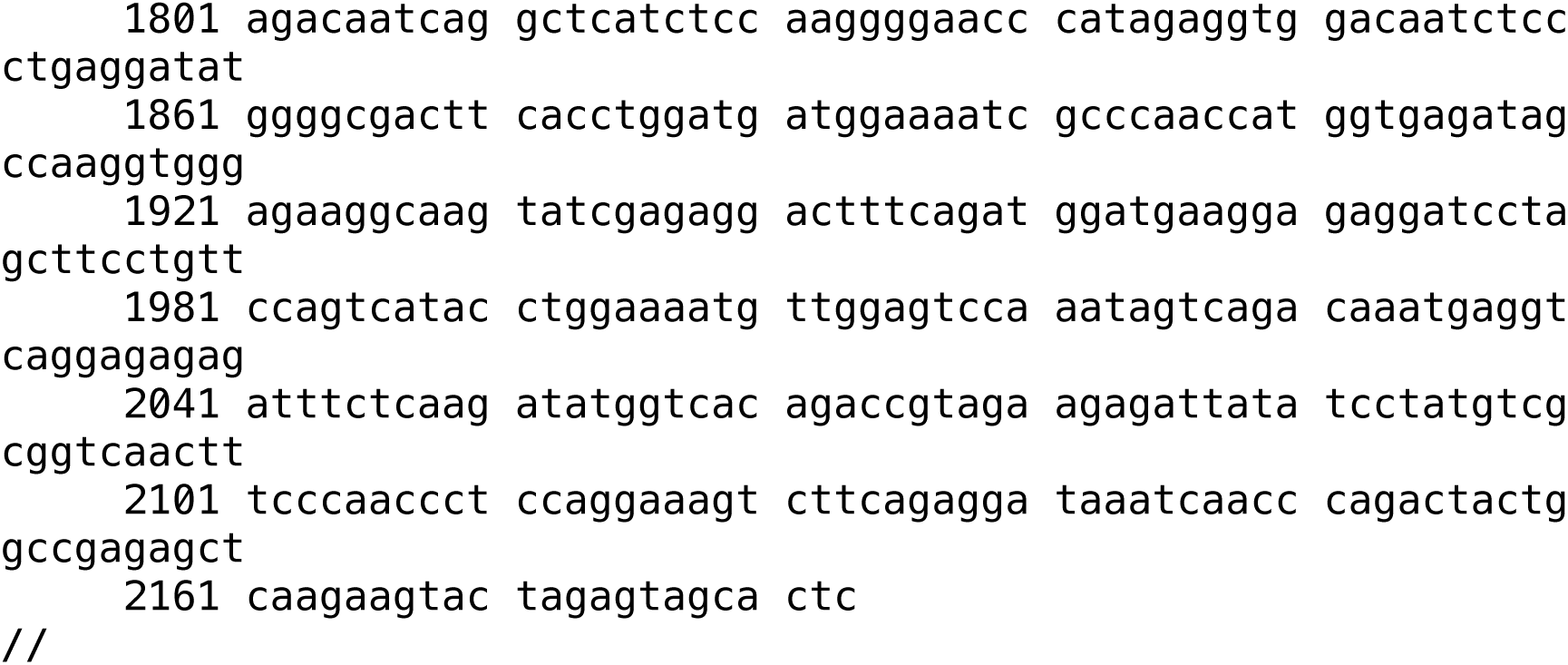
SiR-CRE amplicon reference sequence. This Genbank-format file contains the expected (i.e., based on the published sequence: Addgene #99608) sequence of amplicons obtained from SiR-CRE for SMRT sequencing.

**Supplementary File S4:**
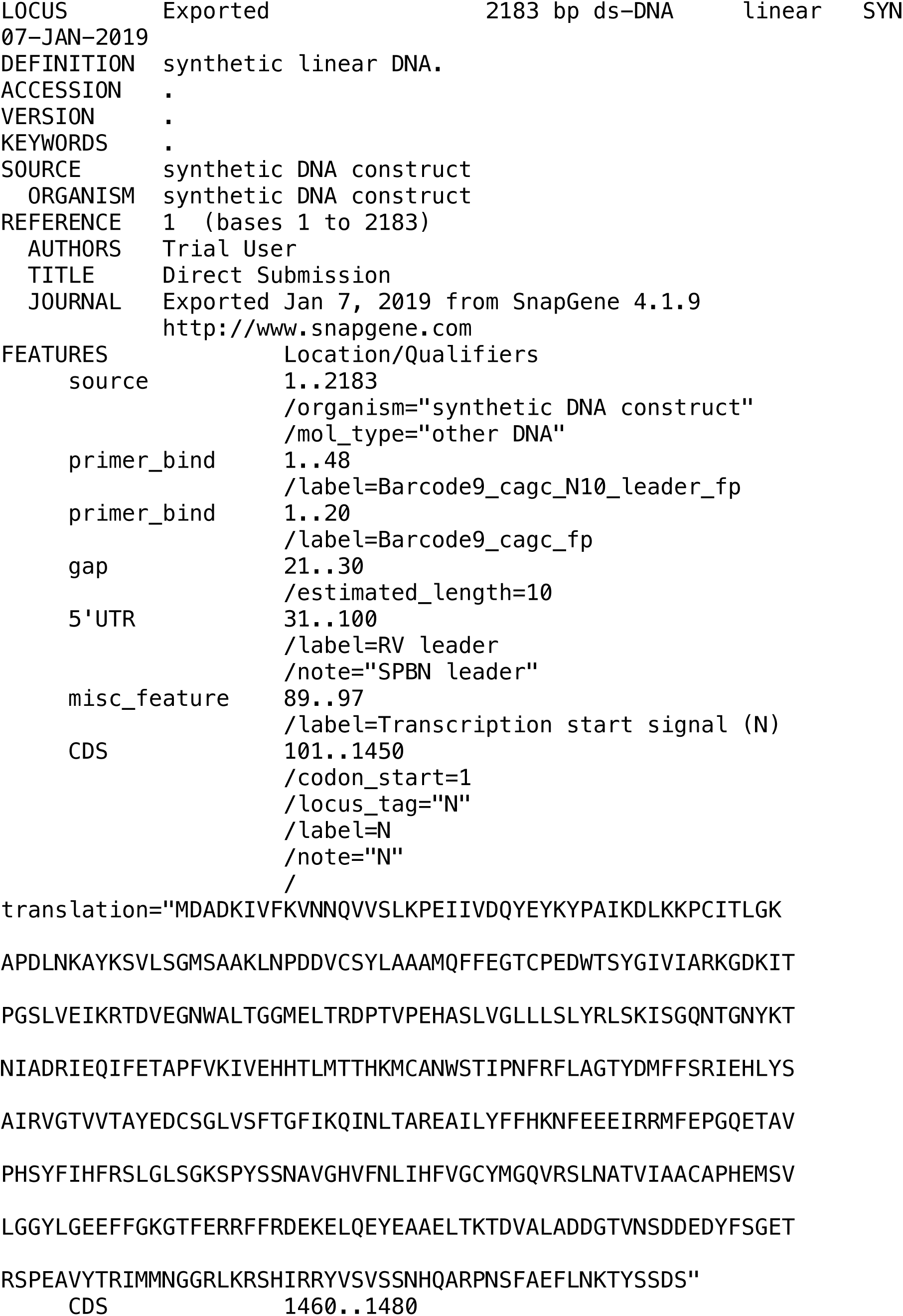
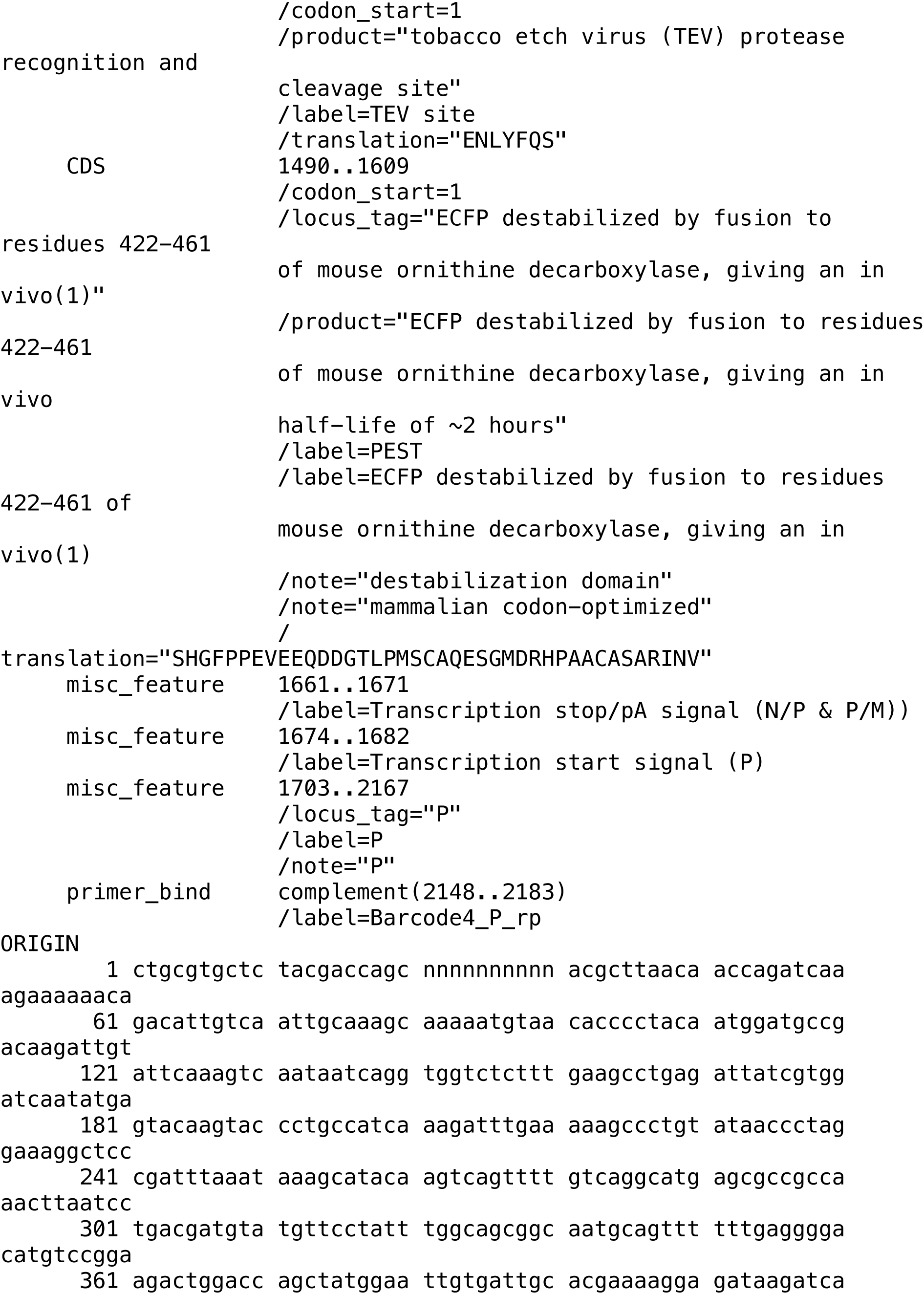
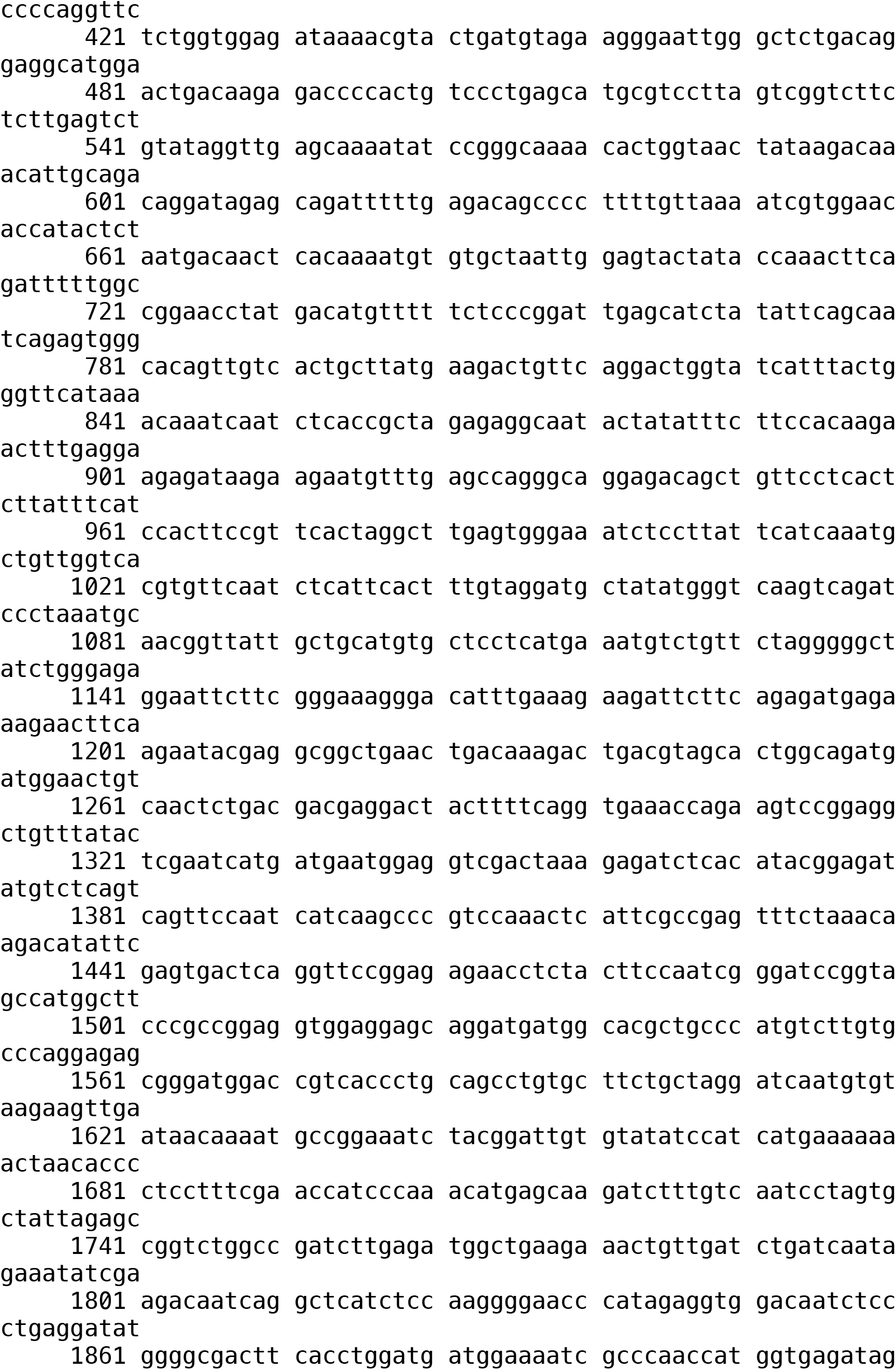
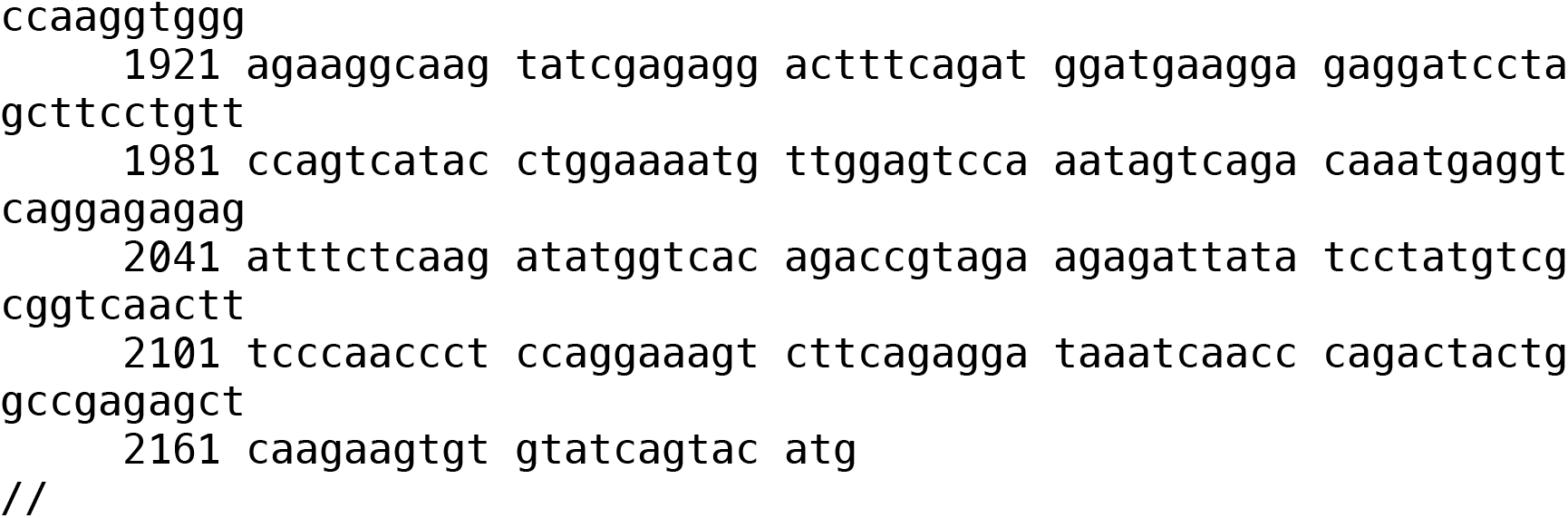
SiR-FLPo amplicon reference sequence. This Genbank-format file contains the expected (i.e., based on the published sequence: Addgene # 99609) sequence of amplicons obtained from SiR-FLPo for SMRT sequencing.

**Supplementary File S5:**
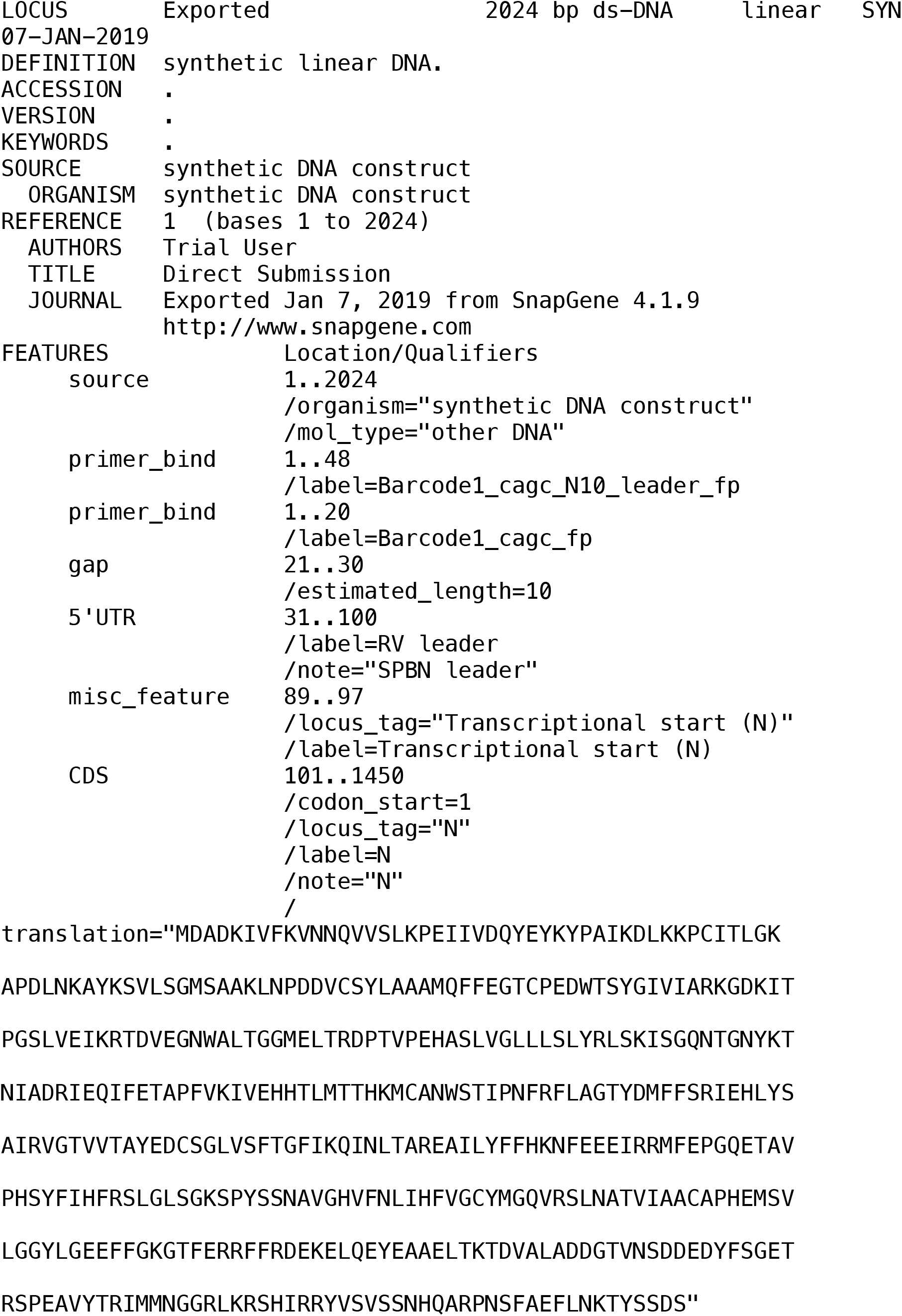
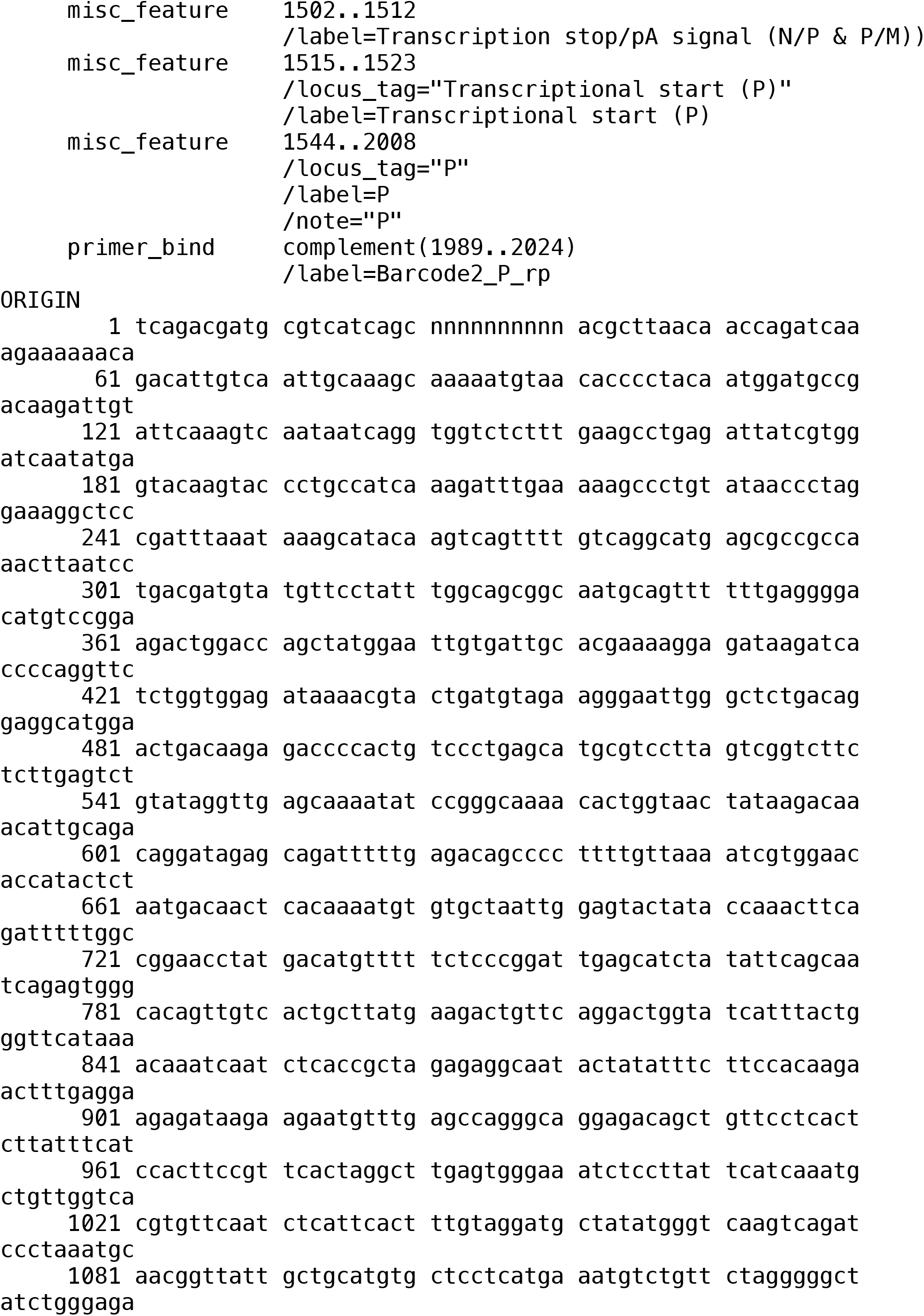
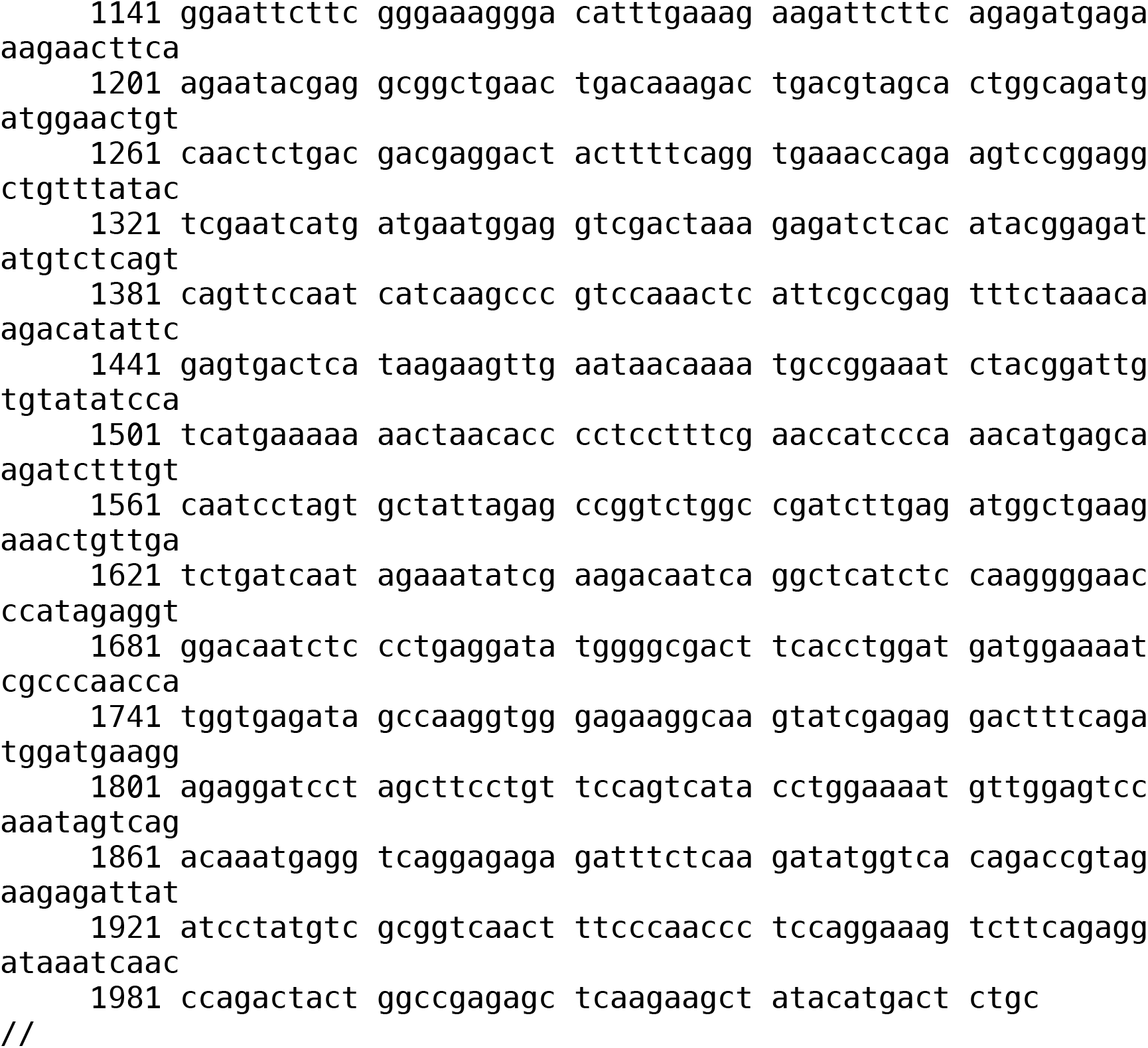
RVΔG-4Cre amplicon reference sequence. This Genbank-format file contains the expected (i.e., based on the published sequence: see Addgene #98034) sequence of amplicons obtained from RVΔG-4Cre for SMRT sequencing.

**Supplementary File S6:**
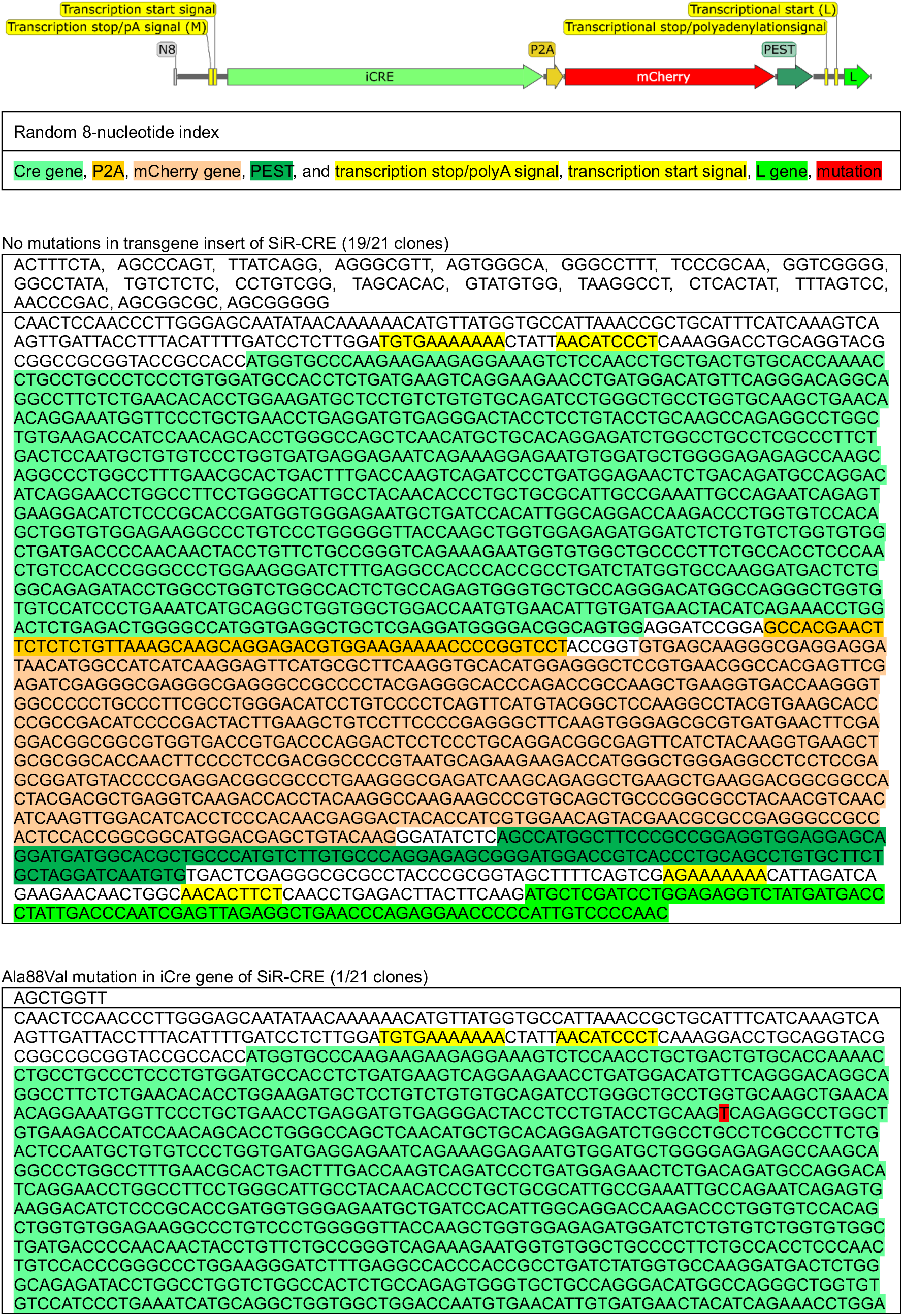
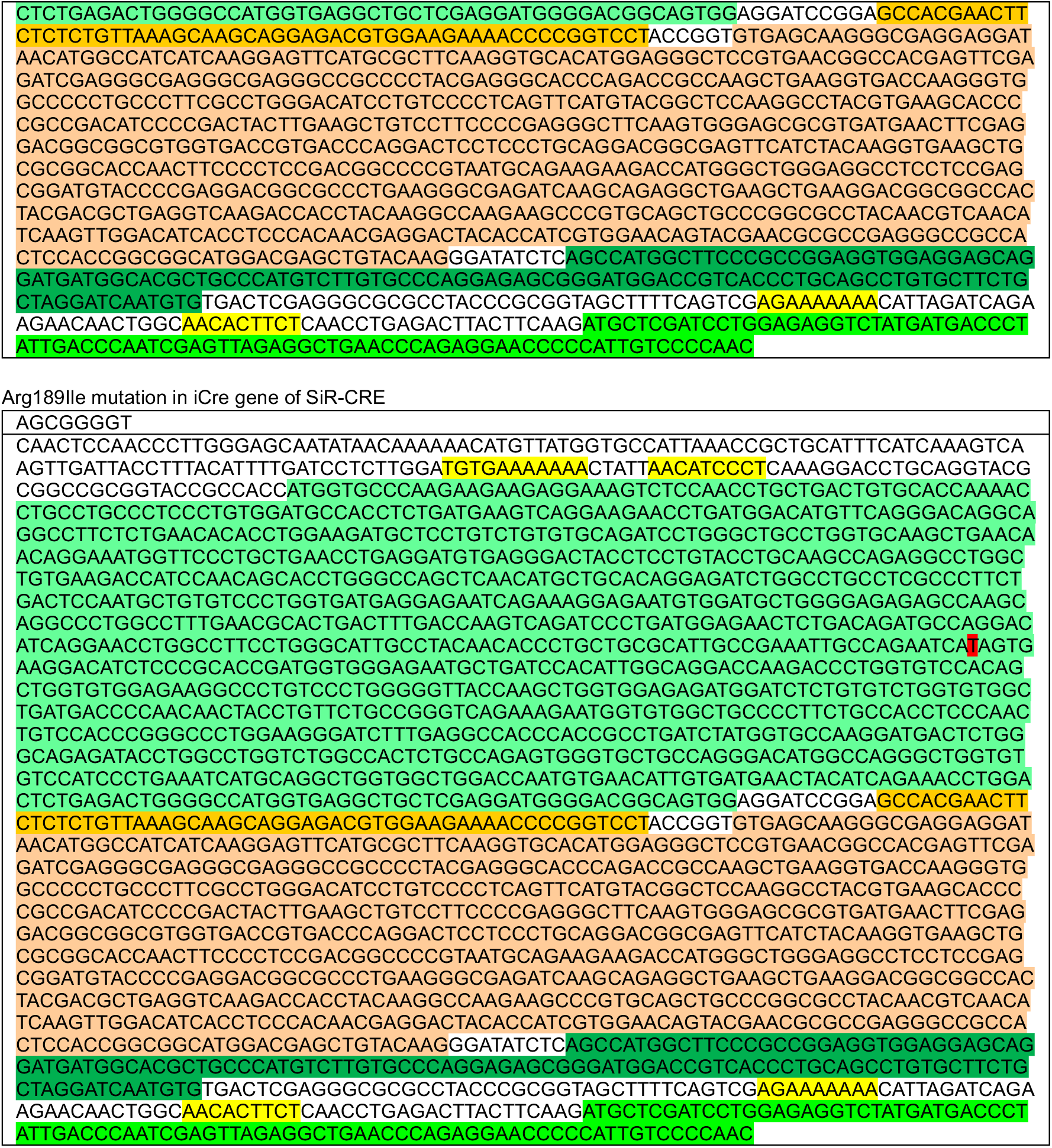
>90% of SiR-CRE viral particles with the mCherry gene intact also have the Cre gene intact, suggesting that most of the SiR-CRE-infected cells that disappear over time in tdTomato reporter mice are dying rather than simply stopping expression of mCherry. We sequenced the transgene inserts for 21 individual SiR-CRE clones (see Methods). 19 out of 21 had no mutations in the Cre gene, and two had one point mutation each (Ala88Val and Arg189Ile). All 21 had an intact mCherry gene. The lack of a large proportion of Cre-knockout mutants is one indication that the majority of red fluorescent neurons in SiR-CRE-injected Ai14 (tdTomato reporter) mice are not labeled only with mCherry, providing evidence that their disappearance is equivalent to their death.

**Supplementary File S7:**
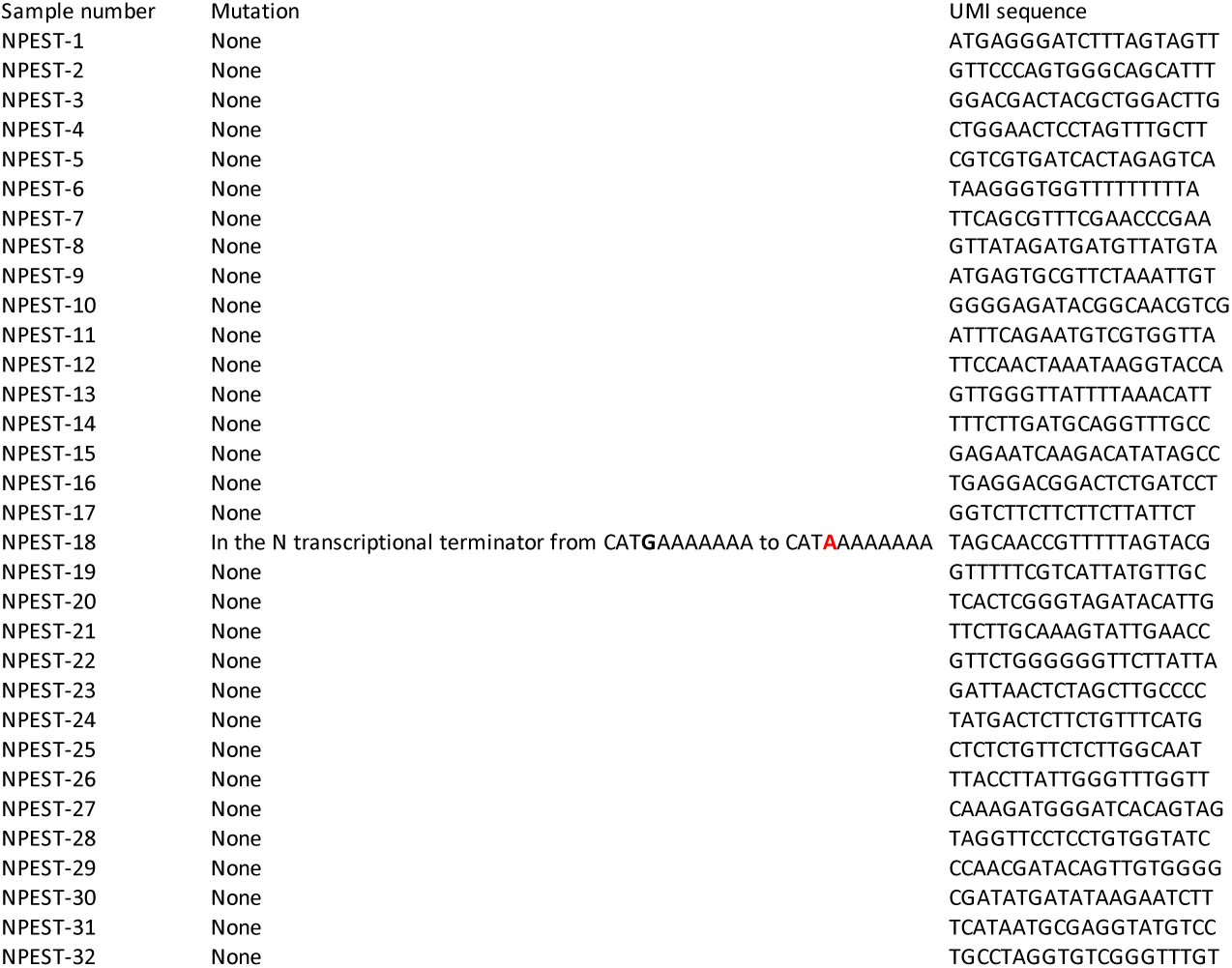
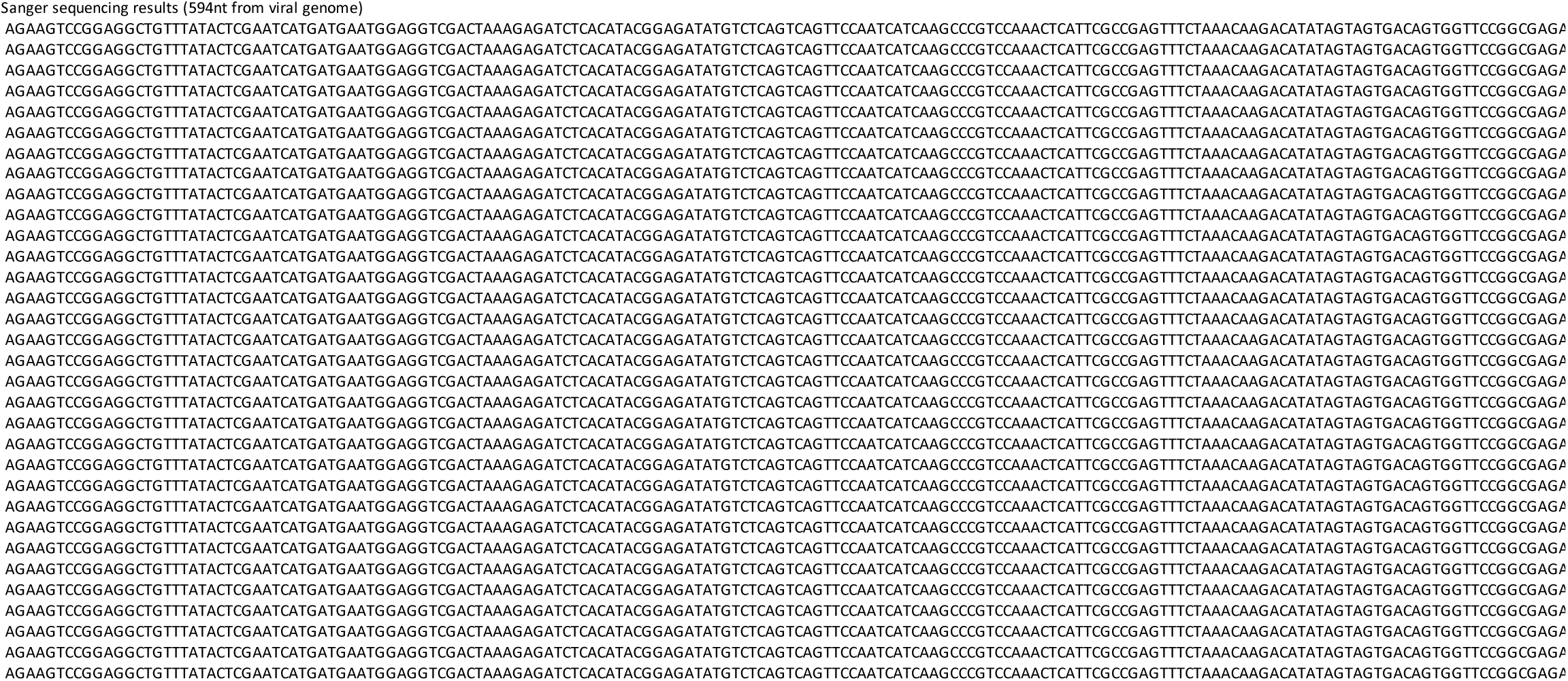
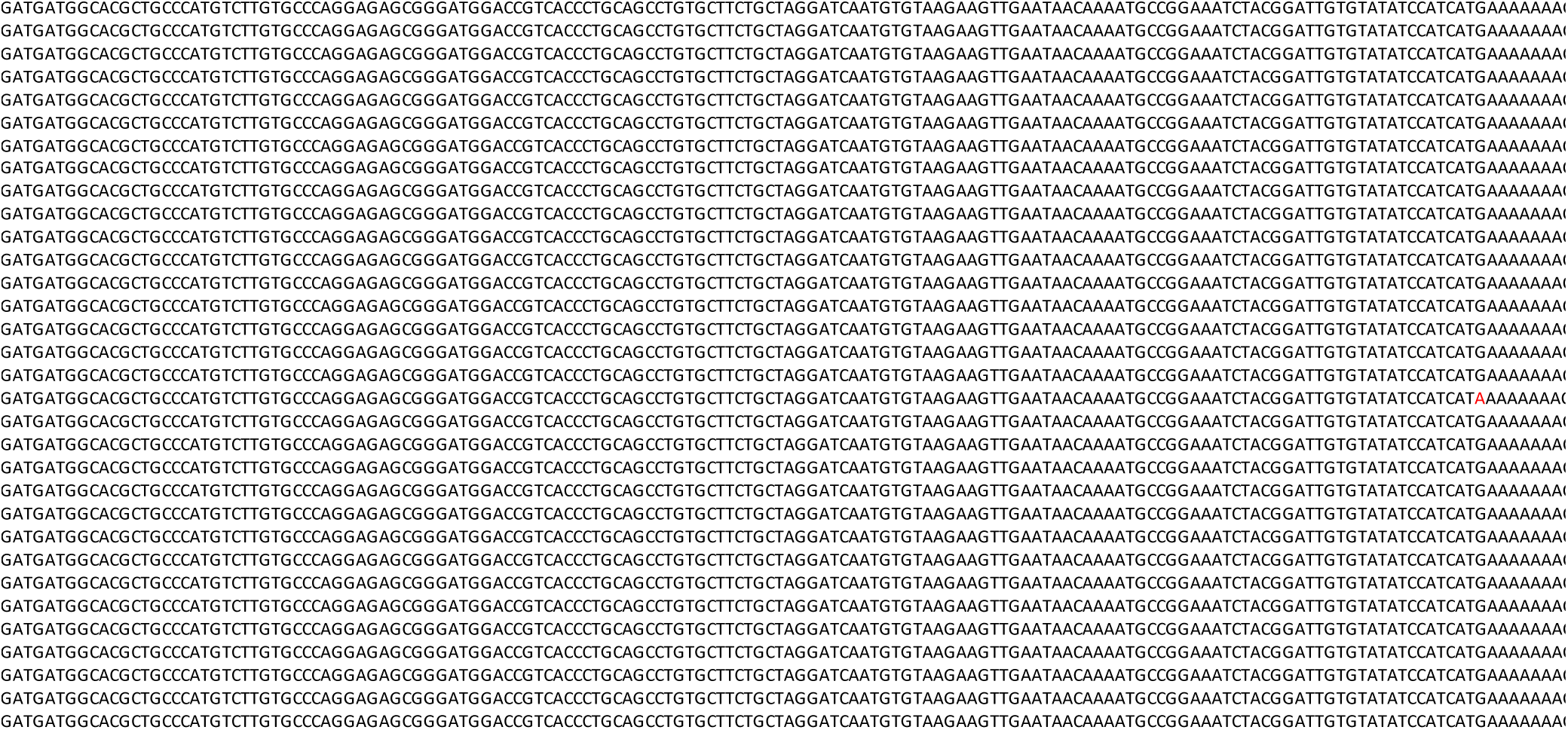
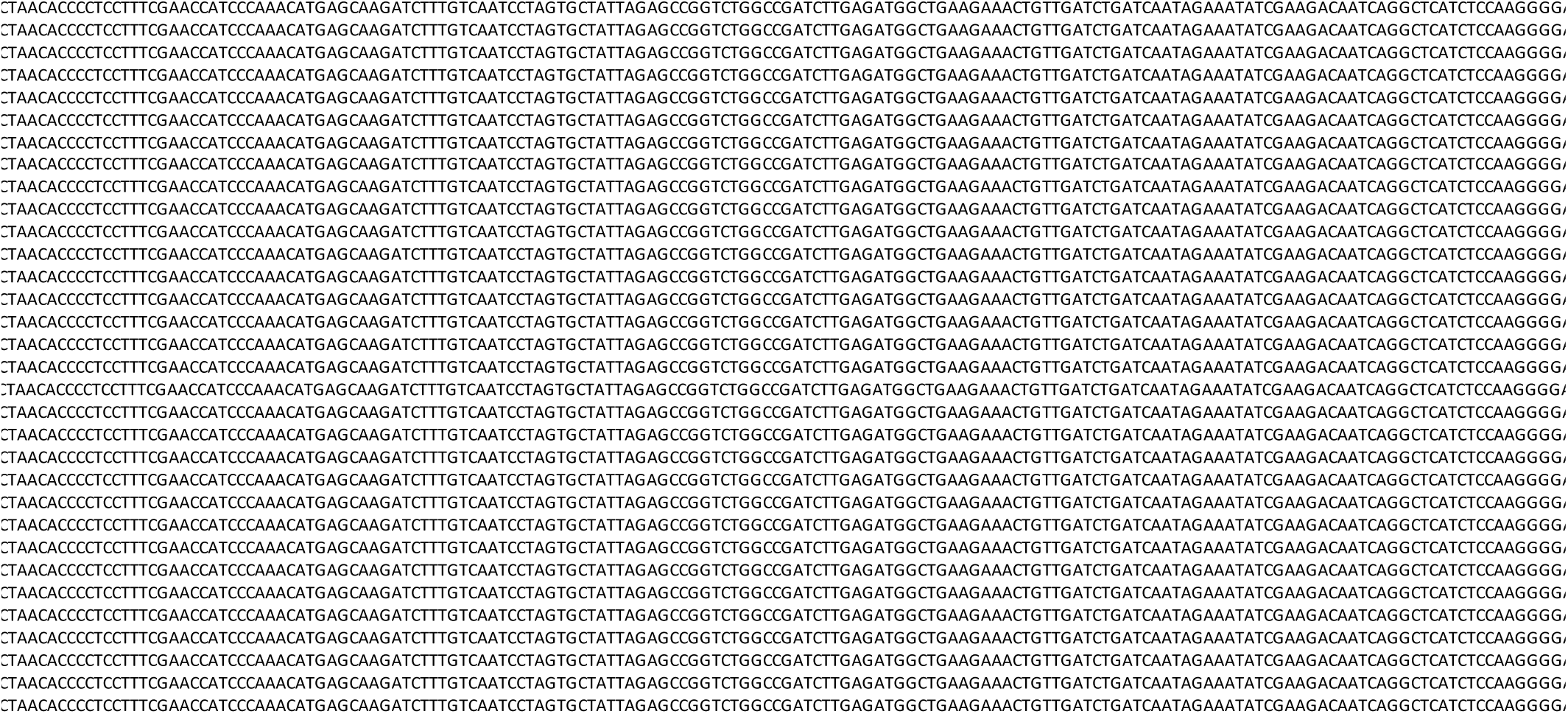
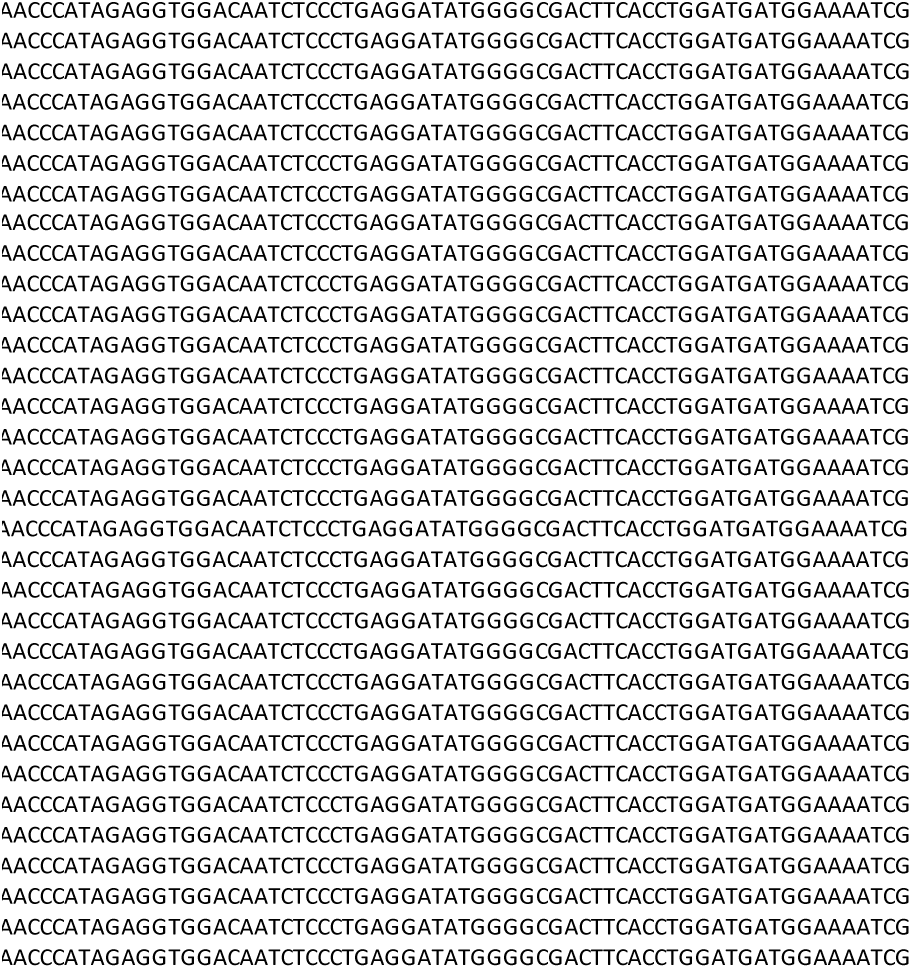
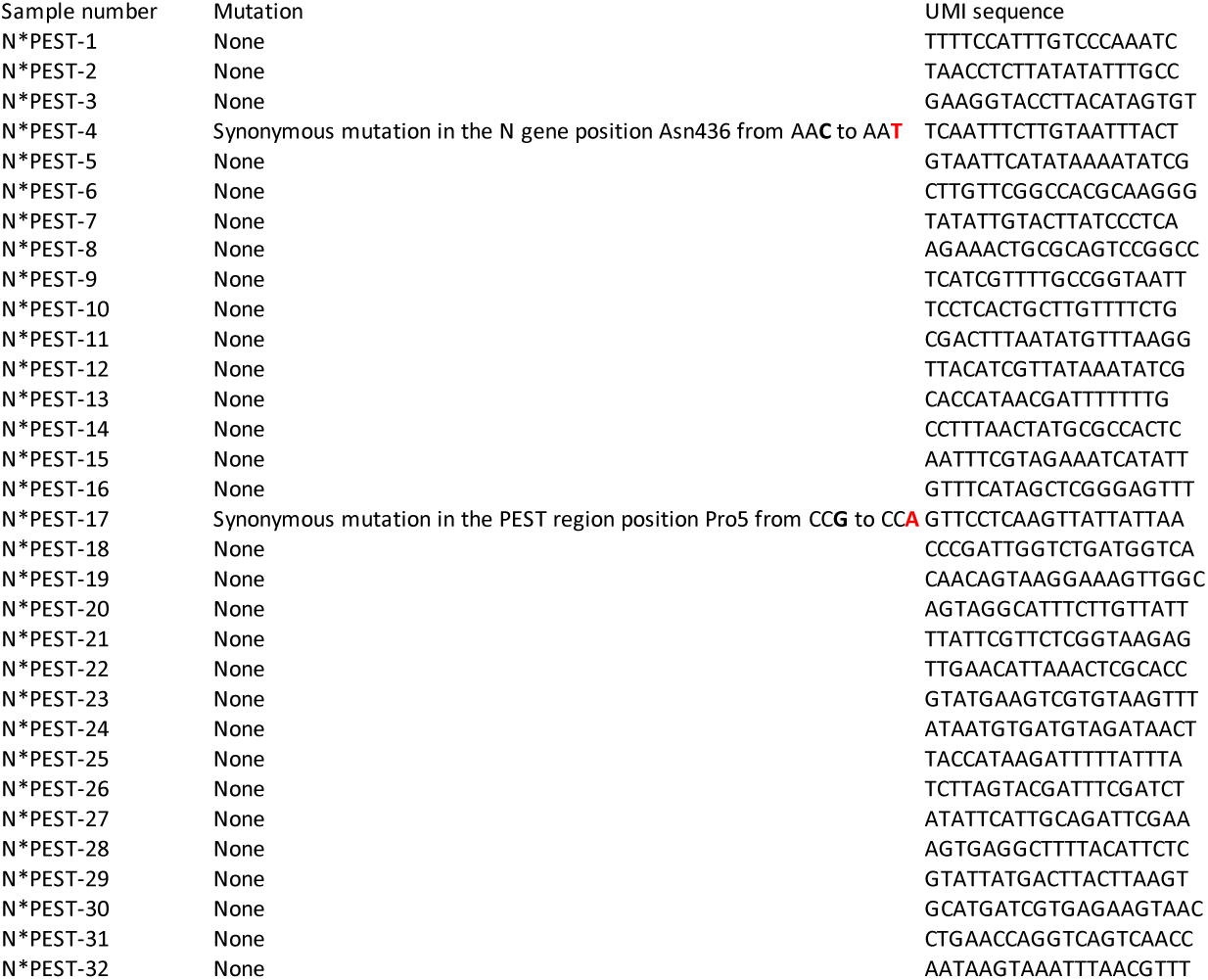
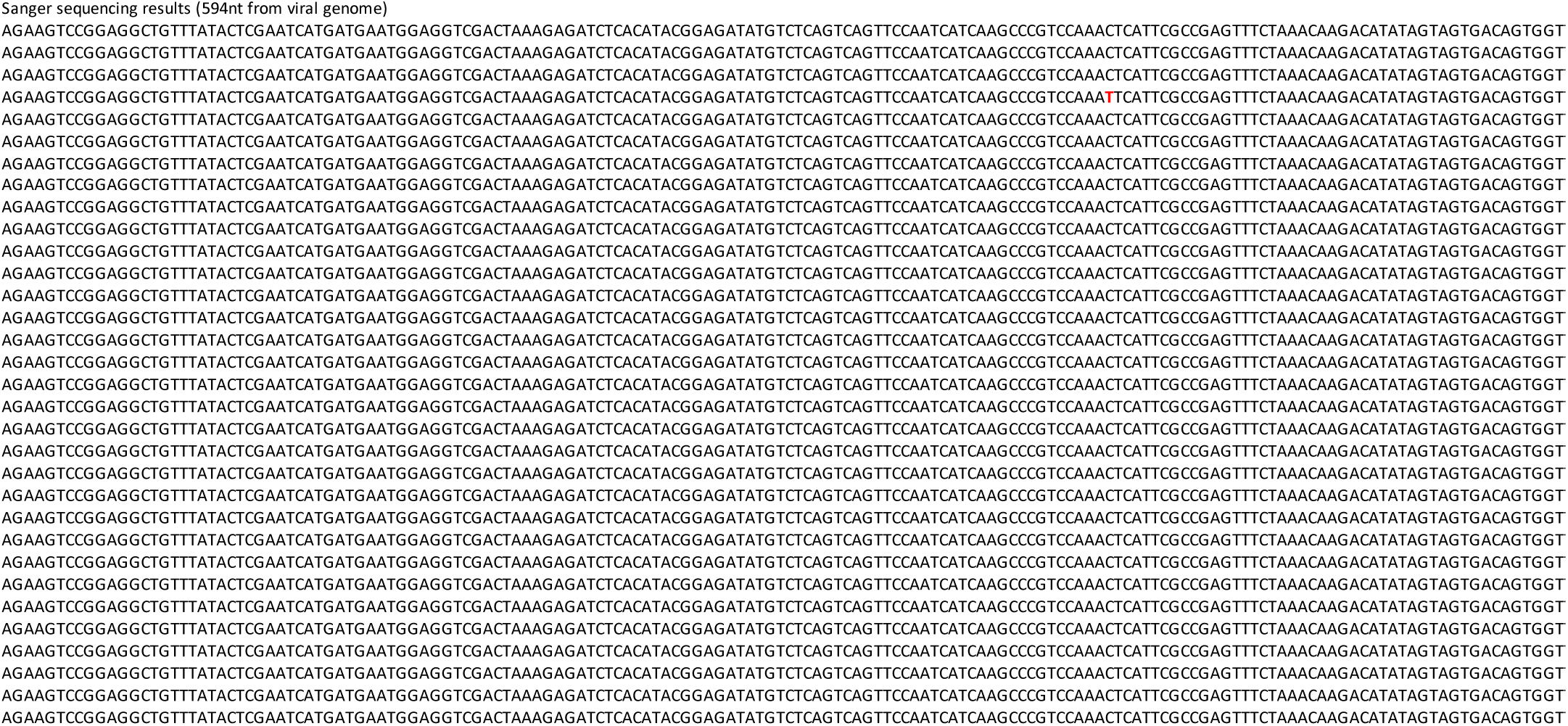
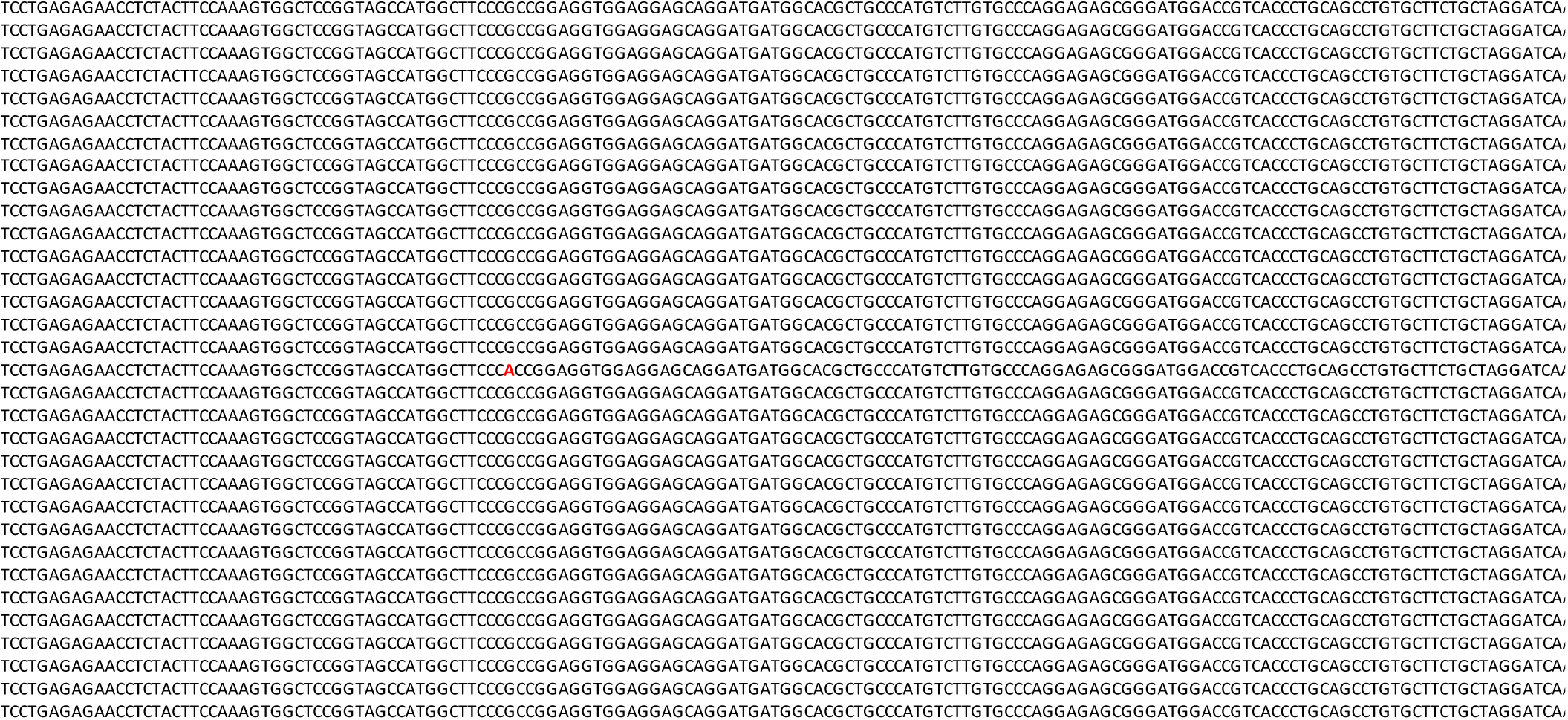
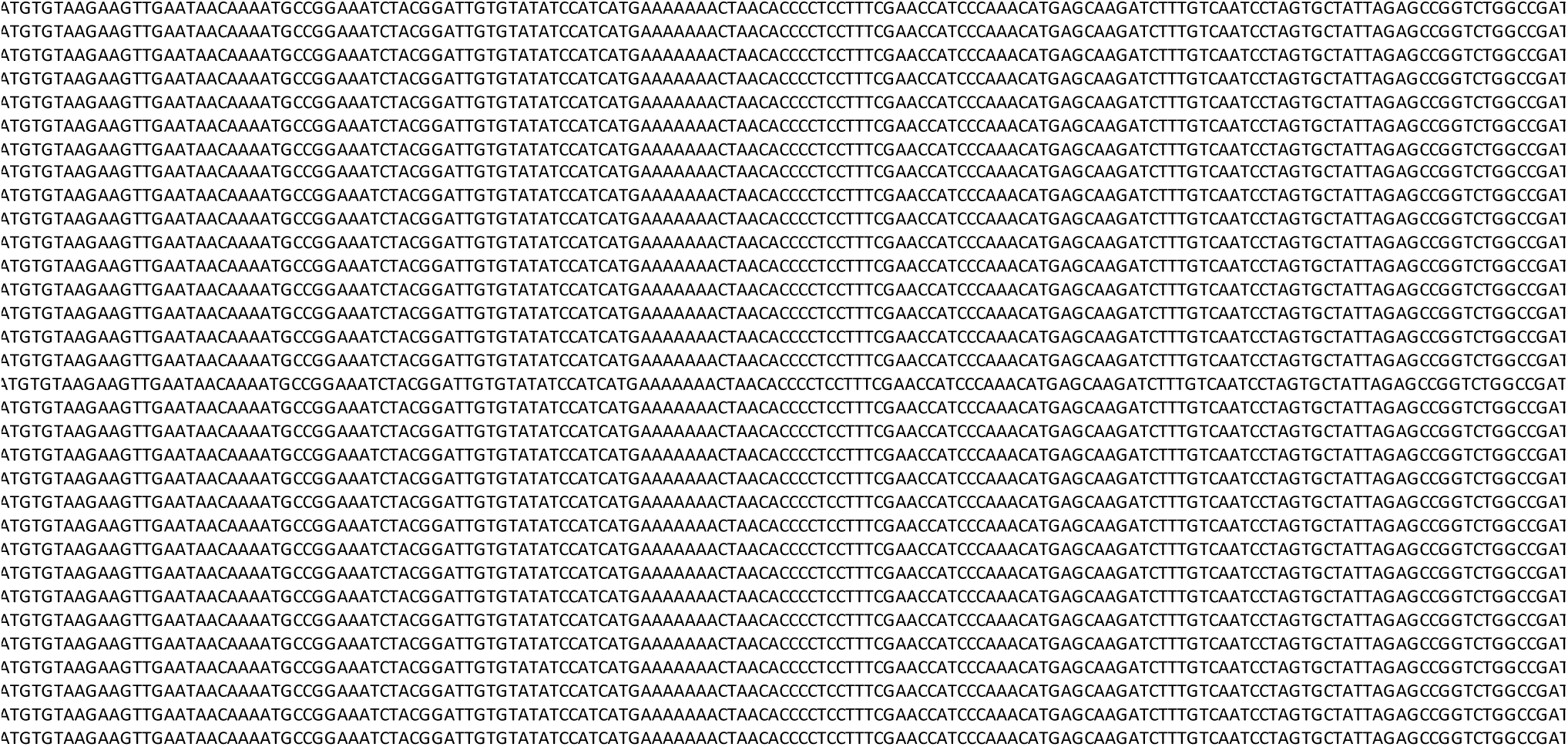
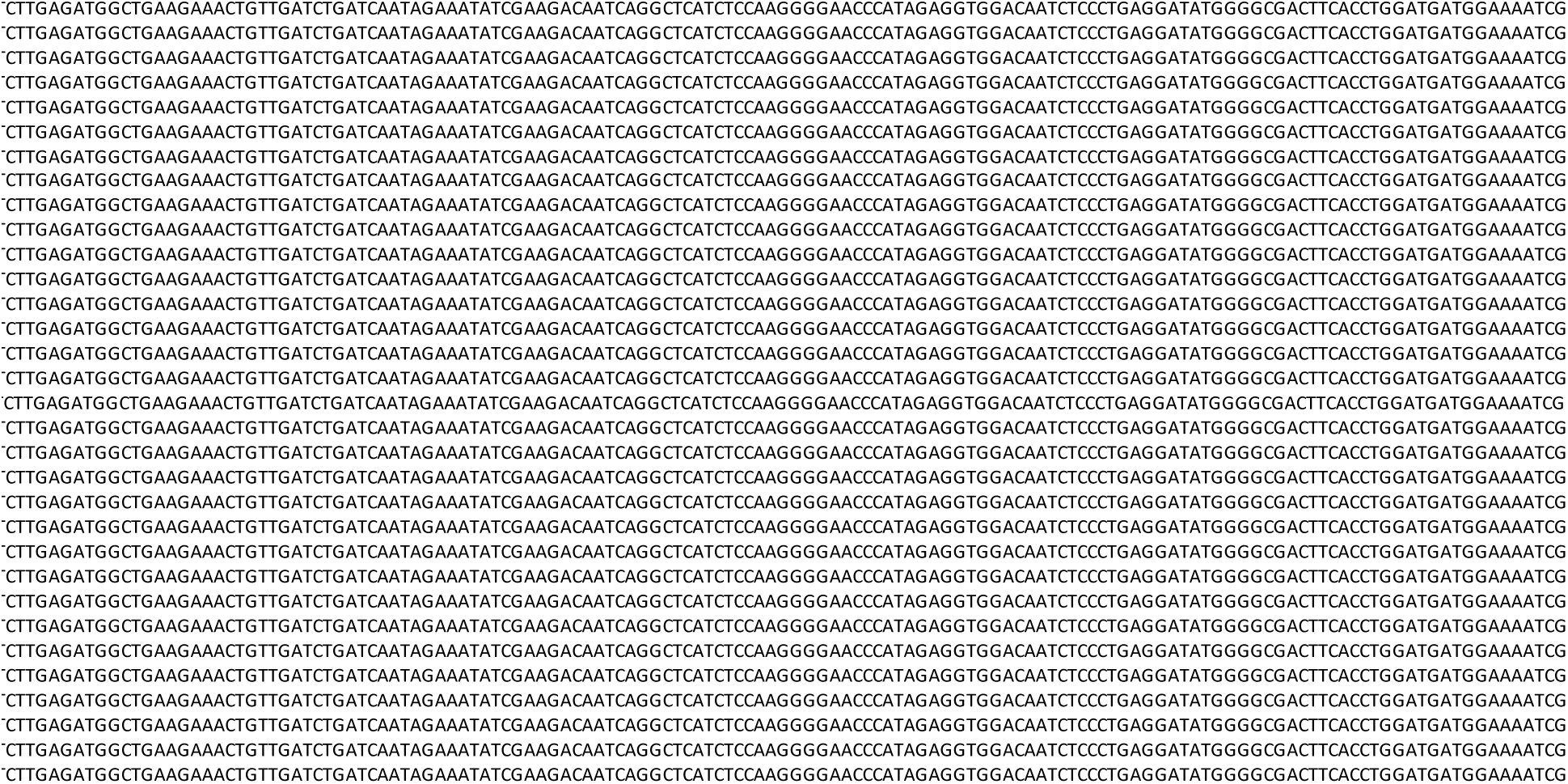
Sequencing results for RVΔG-NPEST-4Cre(EnvA) (“intact”) and RVΔG-NPEST-4Cre(EnvA) (“revertant”), showing that the 3’ additions to the nucleoprotein were as intended in the final stocks used for monosynaptic tracing experiments. We sequenced the complete region of the 3’ addition to the nucleoprotein region from 32 viral particles for each of the two high-titer, EnvA-enveloped preparations of RVΔG-NPEST- 4Cre(EnvA) and RVΔG-N*PEST-4Cre(EnvA) that we made for our *in vivo* experiments. For RVΔG-NPEST-4Cre(EnvA), none of the 32 clones had any mutations in the 3’ addition; one clone had a point mutation in the transcriptional termination signal of the nucleoprotein gene (from CATGAAAAAAA to CATAAAAAAAA). For RVΔG-N*PEST-4Cre(EnvA), 30 of the 32 clones had no mutations in the sequenced region; one clone had a synonymous mutation in the N gene (Asn436, from AAC to AAT), and one other clone had a synonymous mutation in the PEST region (Pro5, from CCG to CCA) after the stop codon.

**Supplementary File S8:**
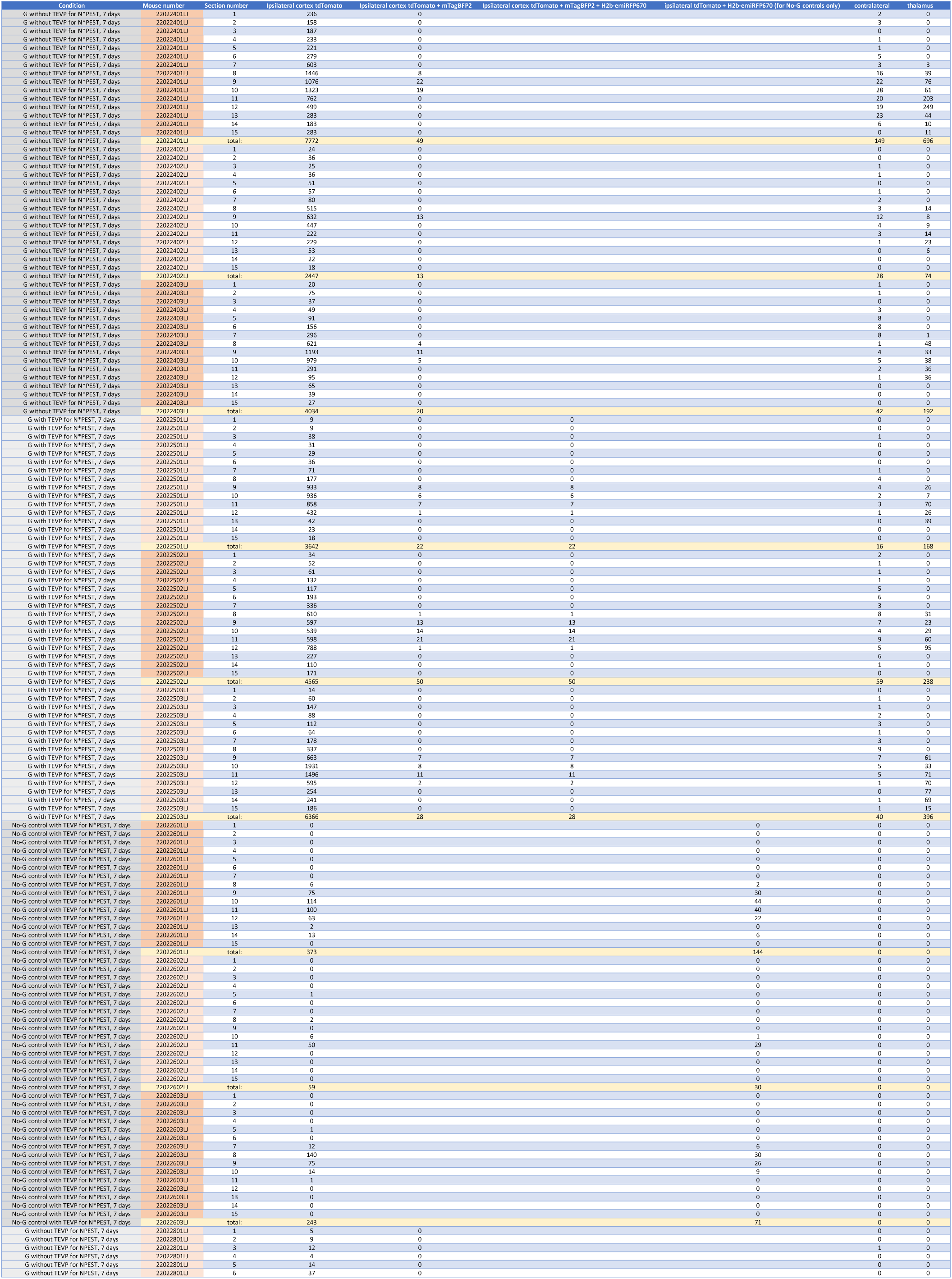
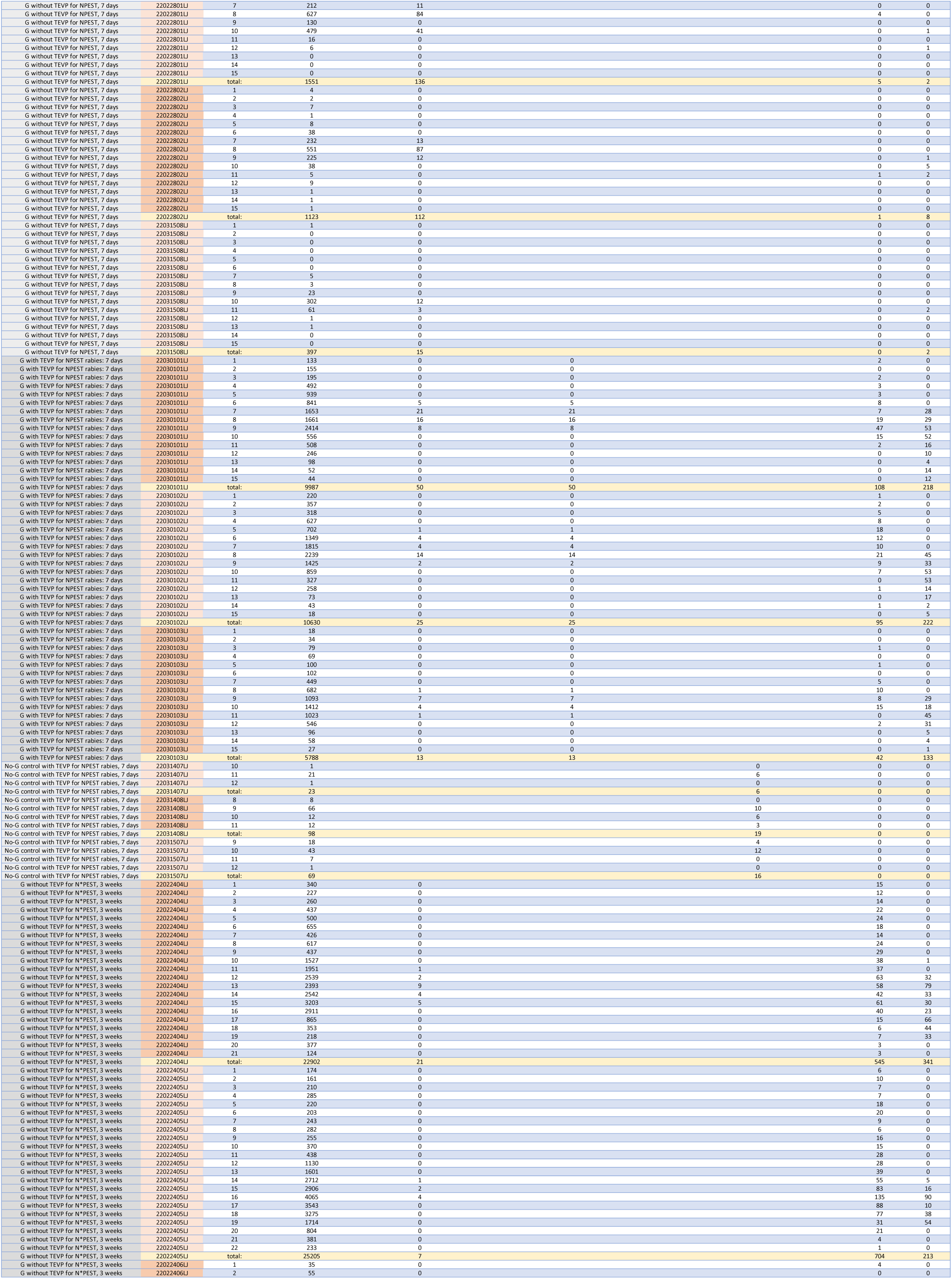
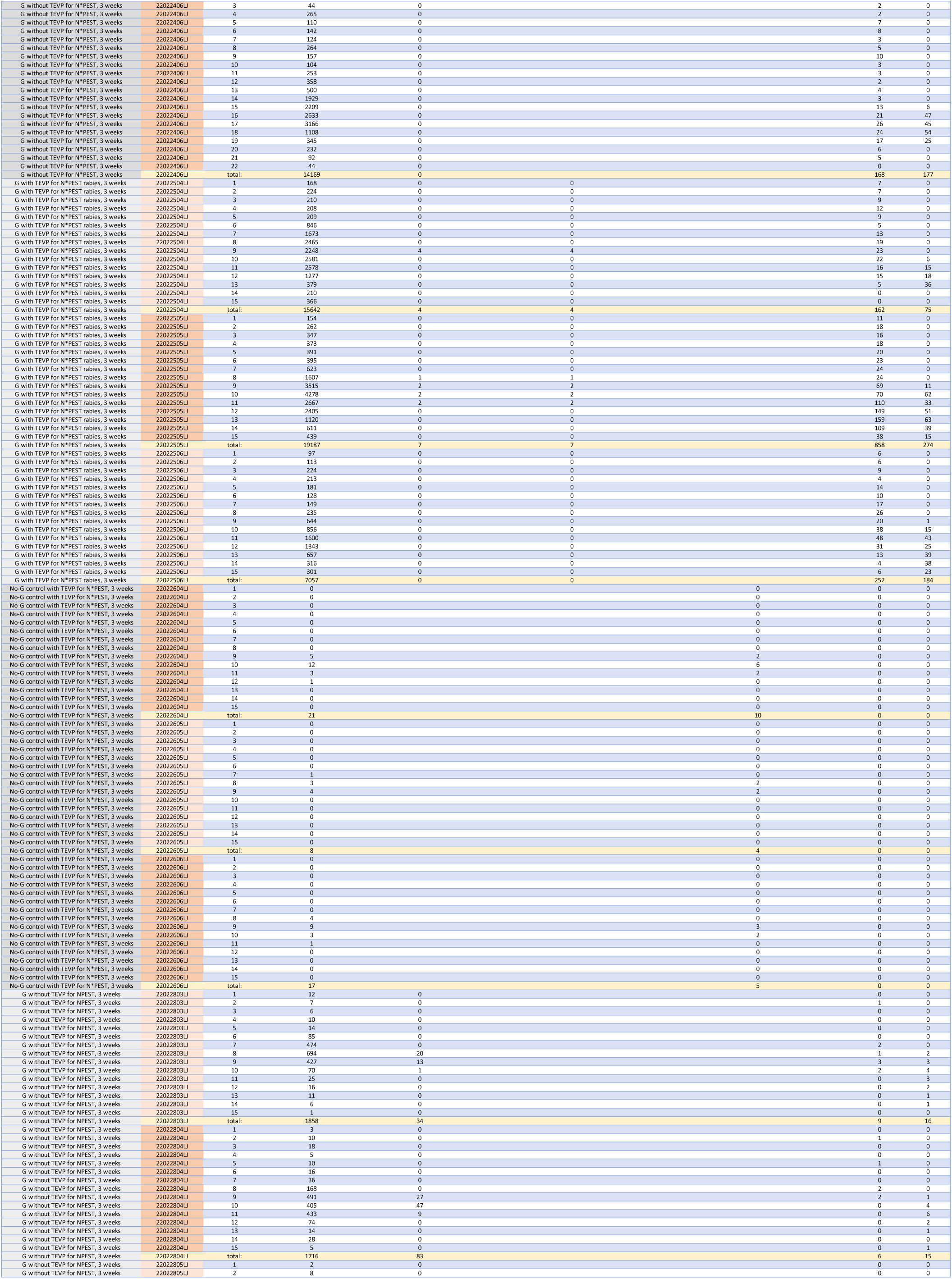
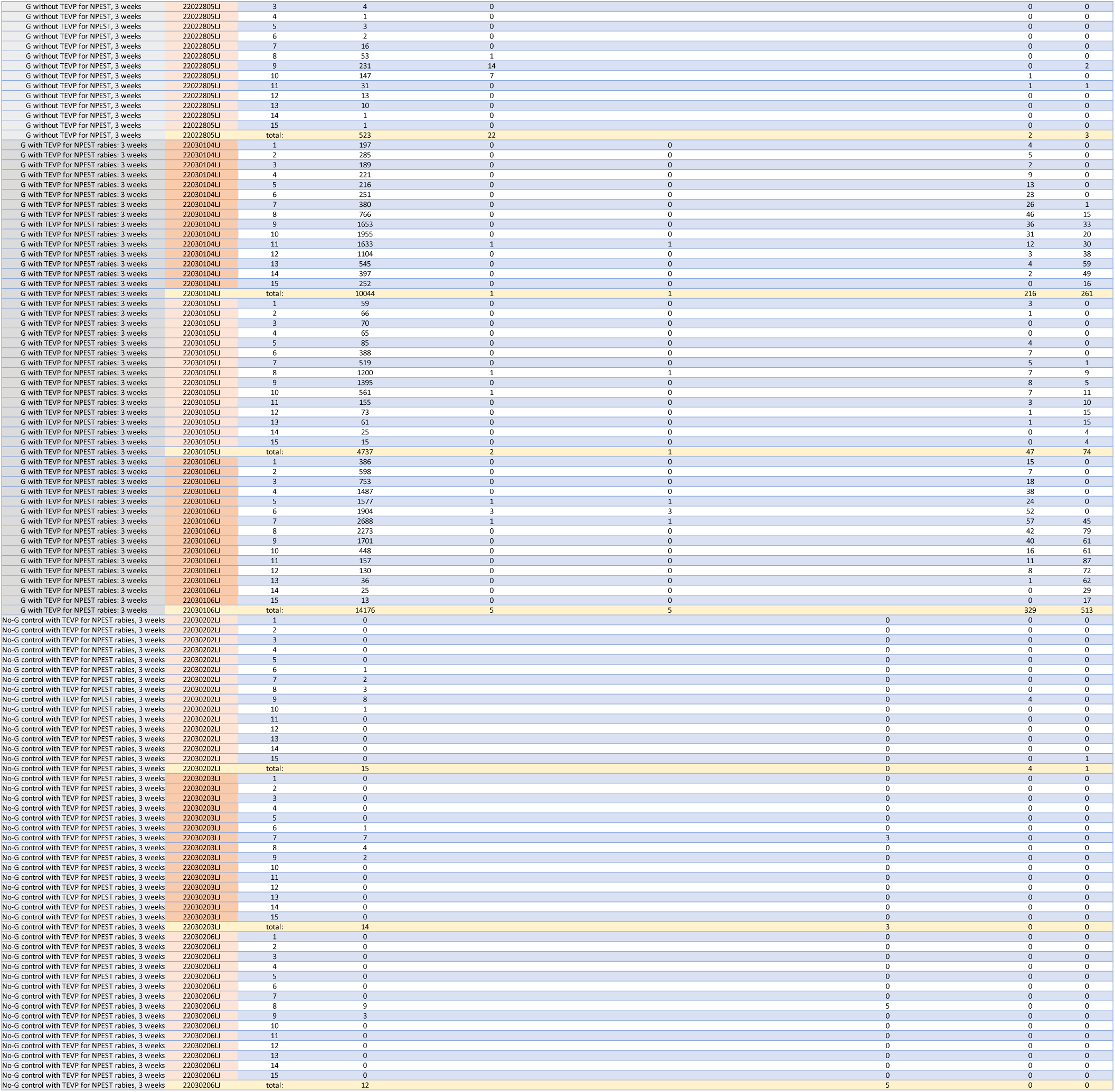
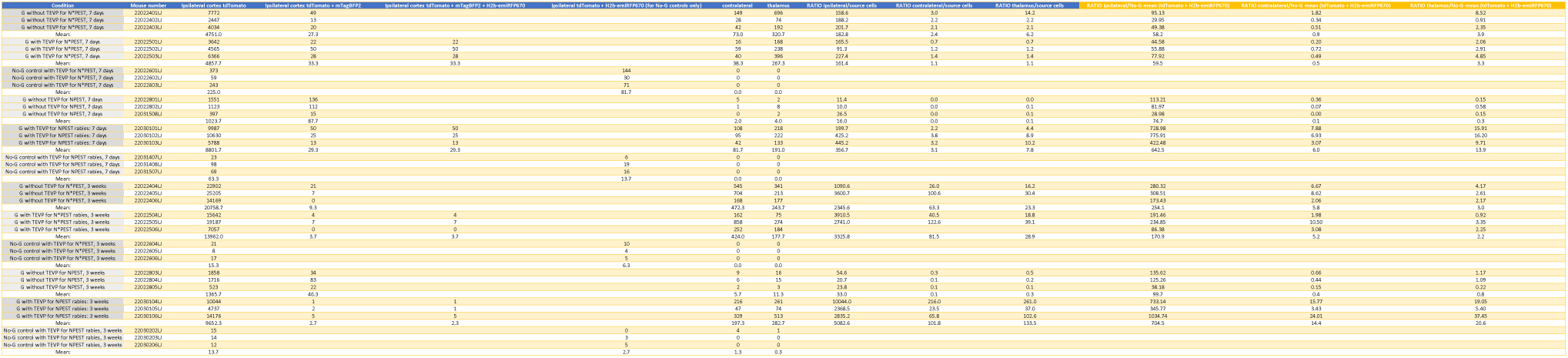
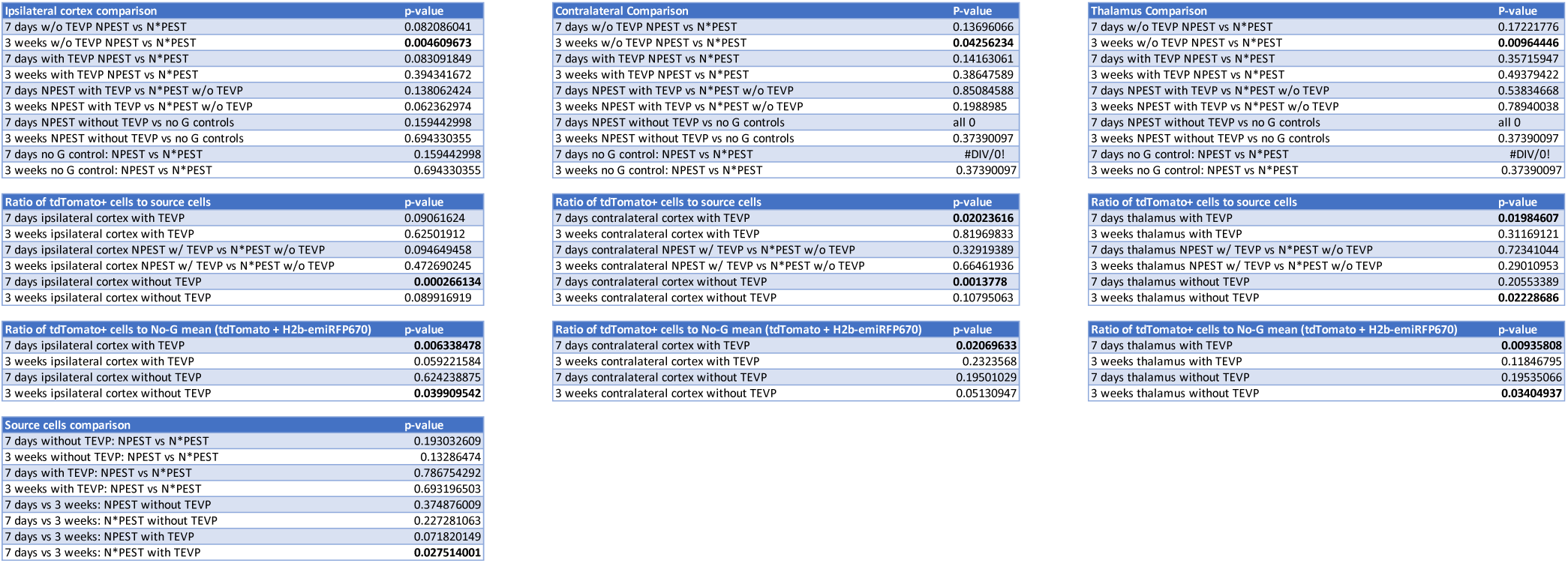
Counts and statistical analyses of labeled neurons in transsynaptic tracing experiments. Sheet 1: Detailed counts of tdTomato-labeled neurons found in every (6th) brain section for all mice, including cell numbers from ipsilateral cortex, contralateral cortex, and thalamus, as well as counts of source cells for each animal; Sheet 2: Total counts of labeled cells for each mouse in each condition, as well as the ratios of tdTomato+ cells to source cells and the average numbers for each condition; Sheet 3: P-values of comparisons using single-factor ANOVAs.

**Supplementary File S9:**
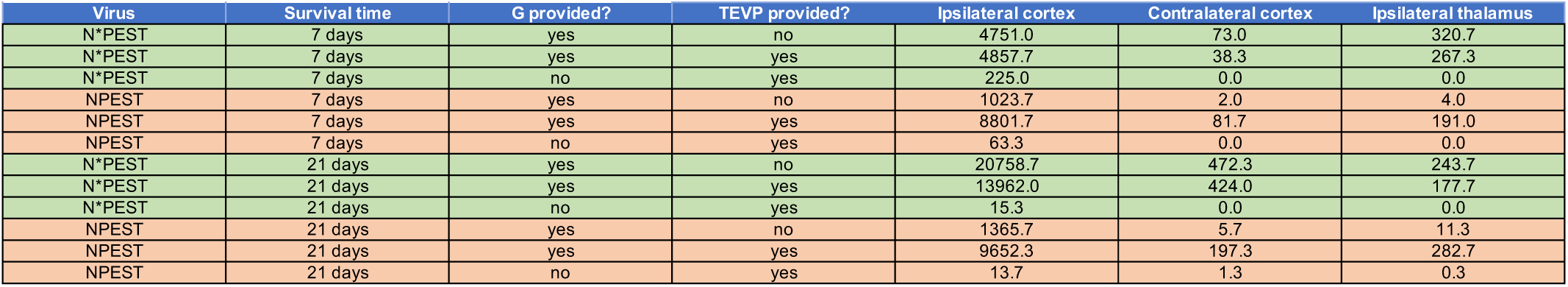
Summary table of mean tdTomato+ cell counts for all conditions. Mean numbers of tdTomato-positive cells in all three quantified brain regions (ipsilateral cortex, contralateral cortex, and thalamus) for each condition in the *in vivo* experiments using the NPEST and N*PEST viruses (cf. Figures 4 & 5 and Supplementary File S8). Numbers are average totals of labeled cells found in a given mouse brain when examining every sixth 50 µm section (see Methods).

## Notes

### Summary of Updates

Minor changes to Supplementary Information: addition of several more graphs to Supplementary Figure S9, and addition of a summary table (Supplementary File S9).

## REFERENCES

1. G. Ugolini, H. G. Kuypers, P. L. Strick, Transneuronal transfer of herpes virus from peripheral nerves to cortex and brainstem. Science 243, 89–91 (1989).

2. G. Ugolini, Specificity of rabies virus as a transneuronal tracer of motor networks: transfer from hypoglossal motoneurons to connected second-order and higher order central nervous system cell groups. J Comp Neurol 356, 457–480 (1995).

3. R. M. Kelly, P. L. Strick, Rabies as a transneuronal tracer of circuits in the central nervous system. J Neurosci Methods 103, 63–71 (2000).

4. I. R. Wickersham, S. Finke, K. K. Conzelmann, E. M. Callaway, Retrograde neuronal tracing with a deletion-mutant rabies virus. Nature Methods 4, 47–49 (2007).

5. I. R. Wickersham et al., Monosynaptic restriction of transsynaptic tracing from single, genetically targeted neurons. Neuron 53, 639–647 (2007).

6. D. Atasoy, Y. Aponte, H. H. Su, S. M. Sternson, A FLEX switch targets Channelrhodopsin-2 to multiple cell types for imaging and long-range circuit mapping. The Journal of neuroscience : the official journal of the Society for Neuroscience 28, 7025–7030 (2008).

7. K. T. Beier et al., Anterograde or retrograde transsynaptic labeling of CNS neurons with vesicular stomatitis virus vectors. P Natl Acad Sci USA 10.1073/pnas.1110854108 (2011).

8. L. Lo, D. J. Anderson, A Cre-dependent, anterograde transsynaptic viral tracer for mapping output pathways of genetically marked neurons. Neuron 72, 938–950 (2011).

9. L. A. Schwarz et al., Viral-genetic tracing of the input-output organization of a central noradrenaline circuit. Nature 10.1038/nature14600 (2015).

10. B. E. Deverman et al., Cre-dependent selection yields AAV variants for widespread gene transfer to the adult brain. Nat Biotechnol 34, 204–209 (2016).

11. D. G. Tervo et al., A Designer AAV Variant Permits Efficient Retrograde Access to Projection Neurons. Neuron 92, 372–382 (2016).

12. J. Dimidschstein et al., A viral strategy for targeting and manipulating interneurons across vertebrate species. Nat Neurosci 19, 1743–1749 (2016).

13. W. B. Zeng et al., Anterograde monosynaptic transneuronal tracers derived from herpes simplex virus 1 strain H129. Mol Neurodegener 12, 38 (2017).

14. L. Luo, E. M. Callaway, K. Svoboda, Genetic Dissection of Neural Circuits: A Decade of Progress. Neuron 98, 865 (2018).

15. S. Ravindra Kumar et al., Multiplexed Cre-dependent selection yields systemic AAVs for targeting distinct brain cell types. Nat Methods 17, 541–550 (2020).

16. J. K. Mich et al., Functional enhancer elements drive subclass-selective expression from mouse to primate neocortex. Cell Rep 34, 108754 (2021).

17. V. Augustine et al., Hierarchical neural architecture underlying thirst regulation. Nature 555, 204–209 (2018).

18. J. Kohl et al., Functional circuit architecture underlying parental behaviour. Nature 556, 326–331 (2018).

19. D. A. Evans et al., A synaptic threshold mechanism for computing escape decisions. Nature 558, 590–594 (2018).

20. M. M. Kaelberer et al., A gut-brain neural circuit for nutrient sensory transduction. Science 361 (2018).

21. S. Chatterjee et al., Nontoxic, double-deletion-mutant rabies viral vectors for retrograde targeting of projection neurons. Nat Neurosci 21, 638–646 (2018).

22. T. R. Reardon et al., Rabies Virus CVS-N2c(DeltaG) Strain Enhances Retrograde Synaptic Transfer and Neuronal Viability. Neuron 89, 711–724 (2016).

23. K. Morimoto, D. C. Hooper, S. Spitsin, H. Koprowski, B. Dietzschold, Pathogenicity of different rabies virus variants inversely correlates with apoptosis and rabies virus glycoprotein expression in infected primary neuron cultures. J Virol 73, 510–518 (1999).

24. L. Jin et al., Long-term labeling and imaging of synaptically-connected neuronal networks in vivo using nontoxic, double-deletion-mutant rabies viruses. bioRxiv10.1101/2021.12.04.471186, 2021.2012.2004.471186 (2021).

25. E. Ciabatti, A. Gonzalez-Rueda, L. Mariotti, F. Morgese, M. Tripodi, Life-Long Genetic and Functional Access to Neural Circuits Using Self-Inactivating Rabies Virus. Cell 170, 382–392 e314 (2017).

26. S. Rogers, R. Wells, M. Rechsteiner, Amino acid sequences common to rapidly degraded proteins: the PEST hypothesis. Science 234, 364–368 (1986).

27. D. A. Steinhauer, J. J. Holland, Rapid evolution of RNA viruses. Annu Rev Microbiol 41, 409–433 (1987).

28. D. A. Steinhauer, J. C. de la Torre, J. J. Holland, High nucleotide substitution error frequencies in clonal pools of vesicular stomatitis virus. J Virol 63, 2063–2071 (1989).

29. J. J. Holland, J. C. De La Torre, D. A. Steinhauer, RNA virus populations as quasispecies. Curr Top Microbiol Immunol 176, 1–20 (1992).

30. E. C. Holmes, C. H. Woelk, R. Kassis, H. Bourhy, Genetic constraints and the adaptive evolution of rabies virus in nature. Virology 292, 247–257 (2002).

31. G. M. Jenkins, A. Rambaut, O. G. Pybus, E. C. Holmes, Rates of molecular evolution in RNA viruses: a quantitative phylogenetic analysis. J Mol Evol 54, 156–165 (2002).

32. M. Combe, R. Sanjuan, Variation in RNA virus mutation rates across host cells. PLoS Pathog 10, e1003855 (2014).

33. I. R. Wickersham, H. A. Sullivan, H. S. Seung, Production of glycoprotein-deleted rabies viruses for monosynaptic tracing and high-level gene expression in neurons. Nature protocols 5, 595–606 (2010).

34. F. Osakada, E. M. Callaway, Design and generation of recombinant rabies virus vectors. Nat Protoc 8, 1583–1601 (2013).

35. I. R. Wickersham, H. A. Sullivan, Rabies viral vectors for monosynaptic tracing and targeted transgene expression in neurons. Cold Spring Harb Protoc 2015, 375–385 (2015).

36. A. P. Weible et al., Transgenic targeting of recombinant rabies virus reveals monosynaptic connectivity of specific neurons. The Journal of neuroscience : the official journal of the Society for Neuroscience 30, 16509–16513 (2010).

37. M. O. Carneiro et al., Pacific biosciences sequencing technology for genotyping and variation discovery in human data. BMC Genomics 13, 375 (2012).

38. C. Wirblich, M. J. Schnell, Rabies virus (RV) glycoprotein expression levels are not critical for pathogenicity of RV. J Virol 85, 697–704 (2011).

39. L. Tao et al., Molecular basis of neurovirulence of flury rabies virus vaccine strains: importance of the polymerase and the glycoprotein R333Q mutation. J Virol 84, 8926–8936 (2010).

40. C. Prehaud, S. Lay, B. Dietzschold, M. Lafon, Glycoprotein of nonpathogenic rabies viruses is a key determinant of human cell apoptosis. J Virol 77, 10537–10547 (2003).

41. L. Madisen et al., A toolbox of Cre-dependent optogenetic transgenic mice for light- induced activation and silencing. Nature neuroscience 15, 793–802 (2012).

42. N. C. Shaner et al., Improved monomeric red, orange and yellow fluorescent proteins derived from Discosoma sp. red fluorescent protein. Nat Biotechnol 22, 1567–1572 (2004).

43. M. Drobizhev, N. S. Makarov, S. E. Tillo, T. E. Hughes, A. Rebane, Two-photon absorption properties of fluorescent proteins. Nat Methods 8, 393–399 (2011).

44. K. W. Huang, B. L. Sabatini, Single-Cell Analysis of Neuroinflammatory Responses Following Intracranial Injection of G-Deleted Rabies Viruses. Front Cell Neurosci 14, 65 (2020).

45. M. Prosniak, D. C. Hooper, B. Dietzschold, H. Koprowski, Effect of rabies virus infection on gene expression in mouse brain. Proc Natl Acad Sci U S A 98, 2758–2763 (2001).

46. L. Madisen et al., Transgenic mice for intersectional targeting of neural sensors and effectors with high specificity and performance. Neuron 85, 942–958 (2015).

47. K. Liu et al., Lhx6-positive GABA-releasing neurons of the zona incerta promote sleep. Nature 548, 582–587 (2017).

48. T. K. Lavin, L. Jin, I. R. Wickersham, Monosynaptic tracing: a step-by-step protocol. J Chem Neuroanat 10.1016/j.jchemneu.2019.101661, 101661 (2019).

49. T. K. Lavin, L. Jin, N. E. Lea, I. R. Wickersham, Monosynaptic Tracing Success Depends Critically on Helper Virus Concentrations. Front Synaptic Neurosci 12, 6 (2020).

50. M. E. Matlashov et al., A set of monomeric near-infrared fluorescent proteins for multicolor imaging across scales. Nat Commun 11, 239 (2020).

51. E. Ciabatti et al., Genomic stability of Self-inactivating Rabies. bioRxiv 10.1101/2020.09.19.304683, 2020.2009.2019.304683 (2020).

52. K. P. Dalton, J. K. Rose, Vesicular stomatitis virus glycoprotein containing the entire green fluorescent protein on its cytoplasmic domain is incorporated efficiently into virus particles. Virology 279, 414–421 (2001).

53. W. P. Duprex, F. M. Collins, B. K. Rima, Modulating the function of the measles virus RNA-dependent RNA polymerase by insertion of green fluorescent protein into the open reading frame. J Virol 76, 7322–7328 (2002).

54. S. Finke, K. Brzozka, K. K. Conzelmann, Tracking fluorescence-labeled rabies virus: enhanced green fluorescent protein-tagged phosphoprotein p supports virus gene expression and formation of infectious particles. J Virol 78, 12333–12343 (2004).

55. M. L. Koser et al., Rabies virus nucleoprotein as a carrier for foreign antigens. Proc Natl Acad Sci U S A 101, 9405–9410 (2004).

56. D. D. Brown et al., Rational attenuation of a morbillivirus by modulating the activity of the RNA-dependent RNA polymerase. J Virol 79, 14330–14338 (2005).

57. S. C. Das, D. Nayak, Y. Zhou, A. K. Pattnaik, Visualization of intracellular transport of vesicular stomatitis virus nucleocapsids in living cells. J Virol 80, 6368–6377 (2006).

58. Y. Klingen, K. K. Conzelmann, S. Finke, Double-labeled rabies virus: live tracking of enveloped virus transport. Journal of virology 82, 237–245 (2008).

59. S. C. Das, D. Panda, D. Nayak, A. K. Pattnaik, Biarsenical labeling of vesicular stomatitis virus encoding tetracysteine-tagged m protein allows dynamic imaging of m protein and virus uncoating in infected cells. J Virol 83, 2611–2622 (2009).

60. A. C. Marriott, C. A. Hornsey, Reverse genetics system for Chandipura virus: tagging the viral matrix protein with green fluorescent protein. Virus Res 160, 166–172 (2011).

61. T. K. Soh, S. P. Whelan, Tracking the Fate of Genetically Distinct Vesicular Stomatitis Virus Matrix Proteins Highlights the Role for Late Domains in Assembly. J Virol 89, 11750–11760 (2015).

62. J. Nikolic, A. Civas, Z. Lama, C. Lagaudriere-Gesbert, D. Blondel, Rabies Virus Infection Induces the Formation of Stress Granules Closely Connected to the Viral Factories. PLoS Pathog 12, e1005942 (2016).

63. J. B. Case, et al., Replication-competent vesicular stomatitis virus vaccine vector protects against SARS-CoV-2-mediated pathogenesis. bioRxiv 10.1101/2020.07.09.196386 (2020).

64. E. A. Gomme, C. Wirblich, S. Addya, G. F. Rall, M. J. Schnell, Immune clearance of attenuated rabies virus results in neuronal survival with altered gene expression. PLoS Pathog 8, e1002971 (2012).

65. T. J. Wiktor, B. Dietzschold, R. N. Leamnson, H. Koprowski, Induction and biological properties of defective interfering particles of rabies virus. J Virol 21, 626–635 (1977).

66. A. Kawai, S. Matsumoto, Interfering and noninterfering defective particles generated by a rabies small plaque variant virus. Virology 76, 60–71 (1977).

67. H. F. Clark, N. F. Parks, W. H. Wunner, Defective interfering particles of fixed rabies viruses: lack of correlation with attenuation or auto-interference in mice. J Gen Virol 52, 245–258 (1981).

68. M. Faber et al., Overexpression of the rabies virus glycoprotein results in enhancement of apoptosis and antiviral immune response. J Virol 76, 3374–3381 (2002).

69. X. Chen, Y. C. Sun, G. M. Church, J. H. Lee, A. M. Zador, Efficient in situ barcode sequencing using padlock probe-based BaristaSeq. Nucleic Acids Res 46, e22 (2018).

70. X. Chen, et al., High-Throughput Mapping of Long-Range Neuronal Projection Using In Situ Sequencing. Cell 179, 772–786 e719 (2019).

71. Y. C. Sun et al., Integrating barcoded neuroanatomy with spatial transcriptional profiling enables identification of gene correlates of projections. Nat Neurosci 24, 873–885 (2021).

72. I. R. Wickersham et al., Lentiviral vectors for retrograde delivery of recombinases and transactivators. Cold Spring Harb Protoc 2015, 368–374 (2015).

73. L. Madisen et al., A robust and high-throughput Cre reporting and characterization system for the whole mouse brain. Nature neuroscience 13, 133–140 (2010).

## REFERENCES FOR SUPPLEMENTARY METHODS

1. Marr RA, et al. (2004) Neprilysin regulates amyloid Beta peptide levels. J Mol Neurosci 22(1-2):5–11.

2. Niwa H, Yamamura K, & Miyazaki J (1991) Efficient selection for high-expression transfectants with a novel eukaryotic vector. Gene 108(2):193–199.

3. Atasoy D, Aponte Y, Su HH, & Sternson SM (2008) A FLEX switch targets Channelrhodopsin-2 to multiple cell types for imaging and long-range circuit mapping. The Journal of neuroscience : the official journal of the Society for Neuroscience 28(28):7025–7030.

4. Subach OM, Cranfill PJ, Davidson MW, & Verkhusha VV (2011) An enhanced monomeric blue fluorescent protein with the high chemical stability of the chromophore. PLoS One 6(12):e28674.

5. Shaner NC, et al. (2004) Improved monomeric red, orange and yellow fluorescent proteins derived from Discosoma sp. red fluorescent protein. Nat Biotechnol 22(12):1567–1572.

6. Turan S, Kuehle J, Schambach A, Baum C, & Bode J (2010) Multiplexing RMCE: versatile extensions of the Flp-recombinase-mediated cassette-exchange technology. J Mol Biol 402(1):52–69.

7. Matsuda T & Cepko CL (2004) Electroporation and RNA interference in the rodent retina in vivo and in vitro. P Natl Acad Sci USA 101(1):16–22.

8. Bates P, Young JA, & Varmus HE (1993) A receptor for subgroup A Rous sarcoma virus is related to the low density lipoprotein receptor. Cell 74(6):1043–1051.

9. Raymond CS & Soriano P (2007) High-efficiency FLP and PhiC31 site-specific recombination in mammalian cells. PLoS One 2(1):e162.

10. Yusa K, Zhou L, Li MA, Bradley A, & Craig NL (2011) A hyperactive piggyBac transposase for mammalian applications. Proc Natl Acad Sci U S A 108(4):1531–1536.

11. Gray DC, Mahrus S, & Wells JA (2010) Activation of specific apoptotic caspases with an engineered small-molecule-activated protease. Cell 142(4):637–646.

12. Yang CF, et al. (2013) Sexually dimorphic neurons in the ventromedial hypothalamus govern mating in both sexes and aggression in males. Cell 153(4):896–909.

13. Kapust RB, et al. (2001) Tobacco etch virus protease: mechanism of autolysis and rational design of stable mutants with wild-type catalytic proficiency. Protein Eng 14(12):993–1000.

14. Gallardo HF, Tan C, & Sadelain M (1997) The internal ribosomal entry site of the encephalomyocarditis virus enables reliable coexpression of two transgenes in human primary T lymphocytes. Gene Ther 4(10):1115–1119.

15. Matlashov ME, et al. (2020) A set of monomeric near-infrared fluorescent proteins for multicolor imaging across scales. Nat Commun 11(1):239.

16. Liu K, et al. (2017) Lhx6-positive GABA-releasing neurons of the zona incerta promote sleep. Nature 548(7669):582–587.

17. Ciabatti E, Gonzalez-Rueda A, Mariotti L, Morgese F, & Tripodi M (2017) Life-Long Genetic and Functional Access to Neural Circuits Using Self-Inactivating Rabies Virus. Cell 170(2):382–392 e314.

18. Chatterjee S, et al. (2018) Nontoxic, double-deletion-mutant rabies viral vectors for retrograde targeting of projection neurons. Nat Neurosci 21(4):638–646.

19. Wickersham IR, et al. (2015) Lentiviral vectors for retrograde delivery of recombinases and transactivators. Cold Spring Harb Protoc 2015(4):368–374.

20. Sullivan HA & Wickersham IR (2015) Concentration and purification of rabies viral and lentiviral vectors. Cold Spring Harb Protoc 2015(4):386–391.

21. Lavin TK, Jin L, & Wickersham IR (2019) Monosynaptic tracing: a step-by-step protocol. J Chem Neuroanat:101661.

22. Tervo DG, et al. (2016) A Designer AAV Variant Permits Efficient Retrograde Access to Projection Neurons. Neuron 92(2):372–382.

23. Wickersham IR & Sullivan HA (2015) Rabies viral vectors for monosynaptic tracing and targeted transgene expression in neurons. Cold Spring Harb Protoc 2015(4):375–385.

24. Wickersham IR, Sullivan HA, & Seung HS (2010) Production of glycoprotein-deleted rabies viruses for monosynaptic tracing and high-level gene expression in neurons. Nature protocols 5(3):595–606.

25. Madisen L, et al. (2010) A robust and high-throughput Cre reporting and characterization system for the whole mouse brain. Nature neuroscience 13(1):133–140.

26. Madisen L, et al. (2012) A toolbox of Cre-dependent optogenetic transgenic mice for light-induced activation and silencing. Nature neuroscience 15(5):793–802.

